# A Bayesian Framework for Detecting Gene Expression Outliers in Individual Samples

**DOI:** 10.1101/662338

**Authors:** John Vivian, Jordan Eizenga, Holly C. Beale, Olena Morozova-Vaske, Benedict Paten

## Abstract

**Objective:** Many antineoplastics are designed to target upregulated genes, but quantifying upregulation in a single patient sample requires an appropriate set of samples for comparison. In cancer, the most natural comparison set is unaffected samples from the matching tissue, but there are often too few available unaffected samples to overcome high inter-sample variance. Moreover, some cancer samples have misidentified tissues or origin, or even composite-tissue phenotypes. Even if an appropriate comparison set can be identified, most differential expression tools are not designed to accommodate comparing to a single patient sample.

**Materials and Methods:** We propose a Bayesian statistical framework for gene expression outlier detection in single samples. Our method uses all available data to produce a consensus background distribution for each gene of interest without requiring the researcher to manually select a comparison set. The consensus distribution can then be used to quantify over- and under-expression.

**Results:** We demonstrate this method on both simulated and real gene expression data. We show that it can robustly quantify overexpression, even when the set of comparison samples lacks ideally matched tissues samples. Further, our results show that the method can identify appropriate comparison sets from samples of mixed lineage and rediscover numerous known gene-cancer expression patterns.

**Conclusions:** This exploratory method is suitable for identifying expression outliers from comparative RNA-seq analysis for individual samples and Treehouse, a pediatric precision medicine group that leverages RNA-seq to identify potential therapeutic leads for patients, plans to explore this method for processing their pediatric cohort.

## BACKGROUND

RNA-seq has been used in the cancer field for a number of purposes: to examine differences between tumor and normal tissue, to classify cancers for diagnostics, and--with the advent of single cell RNA-seq--to characterize tumor heterogeneity ^1–6^. Recently, precision medicine researchers have also begun exploring RNA-seq’s potential to aid in target selection and drug repositioning by identifying clinically actionable aberrations in tumor samples ^7–10^. Clinical studies have demonstrated actionable findings for up to 50% of patients through RNA-seq analysis, particularly for pediatric patients who often do not possess actionable coding DNA mutations ^11–15^. This has led to efforts like Treehouse, a precision medicine initiative for pediatric cancer that evaluates the utility of RNA-seq analysis to inform clinical interpretation. Treehouse has created a large compendium of open-access cancer gene expression data, which is incorporated into their analysis ^16–18^.

Protocols for such precision medicine initiatives involve identifying up-regulated druggable gene targets as therapeutic leads. Differential expression is commonly used to identify up- and down-regulation of genes between two groups of samples. However, most differential expression tools operate best under experimental conditions where both groups consist of several technical replicates, or lacking that, biological replicates ^19–22^. Thus, most existing tools are poorly suited to the clinical setting, where one group consists of only a single biological replicate from one patient (N-of-1), and the other comparison group is a library of diverse potential comparison samples. In particular, none of the existing methods have any way of suggesting what an appropriate subset of the sample library should be used for comparison. This limitation is especially acute in cancer, where uncertainty as to the cell of origin, histological complexity, and metastasis can make it difficult to identify the appropriate reference tissues for a sample ^23^. While some work exists to address statistical uncertainty of working with N-of-1 samples ^24^, we focus on solving the second problem, which is the principled selection of an appropriate comparison set.

Existing N-of-1 protocols compare targeted genes in an N-of-1 sample to an outlier cutoff generated from a large compendium of either cancer samples or unaffected tissue in order to determine whether a gene is up-regulated ^16,25–27^. While this outlier cutoff method is fast, there are some notable drawbacks. Applying a cutoff binarizes data, which makes it difficult to meaningfully rank outliers or to be aware of samples just short of meeting the cutoff. This cutoff method is also intended for Gaussian distributions, which is empirically common for gene expression within a tissue group, but not typical when considering the distribution of expression across tissues ^25^.

Ultimately, the most difficult problem is justifying the choice of what samples constitute the comparison set that generates the cutoff, since different comparison sets will identify different genes as outliers. Many comparison datasets are small (almost half of The Cancer Genome Atlas’ (TCGA) normal tissues have 10 or fewer samples), so they lack the statistical power to characterize the variability of the expression landscape in the normal tissue on their own ^28^. This power can be increased by also including samples from different tissues, but including tissues with larger sample sizes can drown out the information from the matched tissue. In addition, it is unclear which other tissues should be included in the pooled comparison set.

These concerns led us to propose a new approach for identifying outliers for N-of-1 samples. In contrast to previous methods, our method adaptively constructs a meaningful comparison set and avoids selection bias by automatically weighting the background sets to generate a consensus distribution of expression. It then uses the consensus distribution to quantify over-expression for genes of interest.

## Materials and Methods

The core of our method is a Bayesian statistical model for the N-of-1 sample’s gene expression. The model implicitly assumes that the sample’s gene expression can be approximated by a convex mixture of the gene expression of the background datasets. The coefficients of this mixture are shared across genes, much like a linear model in which each data point is the vector of expressions for a gene across the background datasets. In addition, we model expression for each gene from each background dataset as a random variable itself. This allows us to incorporate statistical uncertainty from certain background sets’ small sample size directly in the model without drowning out any background set’s contribution through pooling **(Figure 1)**.

**Figure 1.**
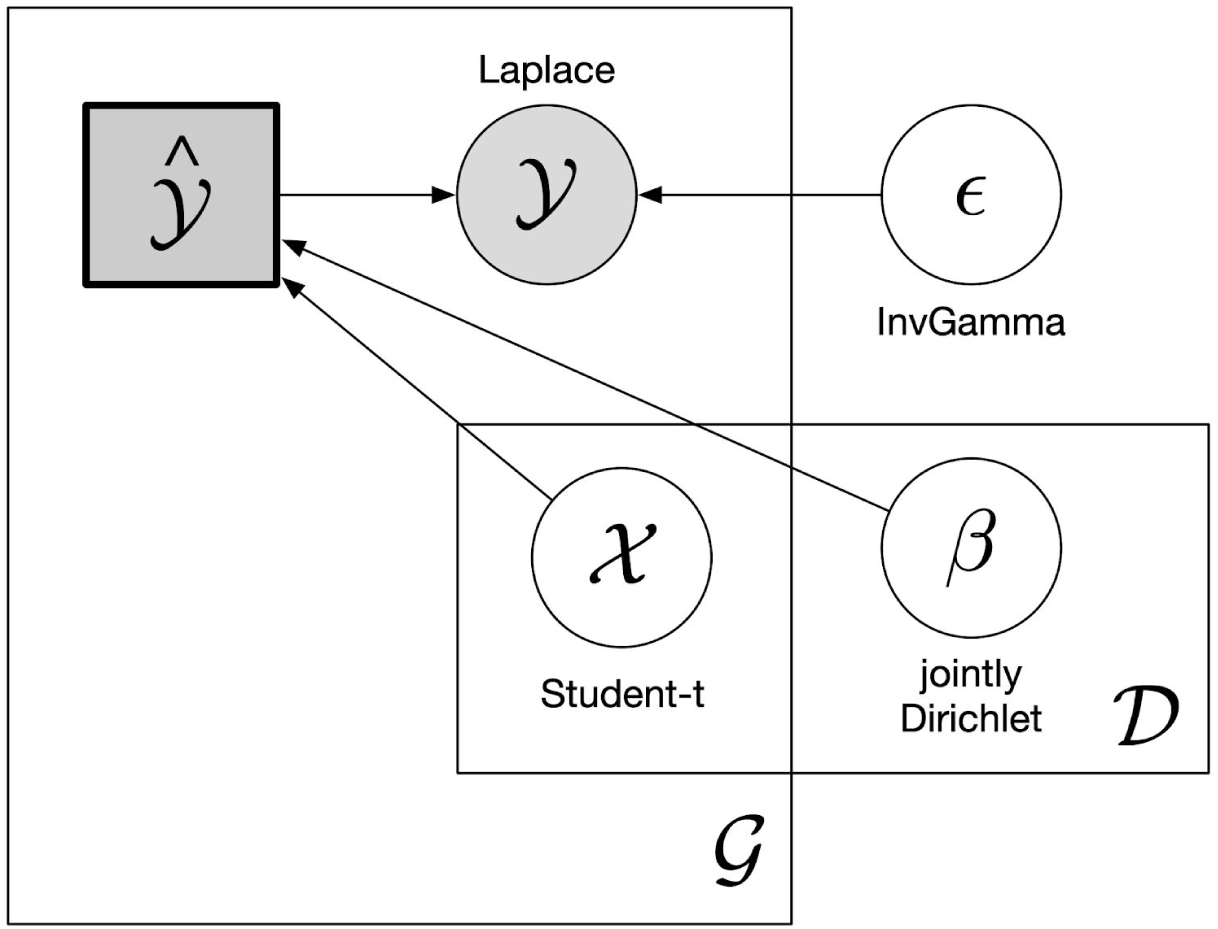
Bayesian plate notation of the model. The plates (G) and (D) stand for Gene and Dataset respectively. *x* represents gene expression for one background dataset and is multiplied by the dot product of *β*^*T*^ to produce the convex combination *ŷ.* We specify “jointly Dirichlet” as there is not one Dirichlet distribution per background dataset. Instead, each background dataset is one component of the Dirichlet *α* vector. We add Laplacian error *ϵ* to the expected expression *ŷ.* when modeling the observed expression of a new sample *y*.

### Model Specification

Suppose we have a *n* background datasets for expression, which we will call *X*_1_,…, *X*_*n*_. Within each background set *d* = 1,2, …,*n*, we model the expression of gene *g* as a normal-distributed random variable:

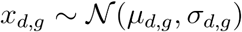

The parameters of each of these normal distributions are distributed according to a shallow but proper normal-inverse gamma prior:

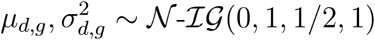

Next we assume there is another unobserved random variable 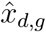 from the same distribution 𝒩(*µ*_*d,g*_, *σ*_*d,g*_). Conceptually, this corresponds to the expression value that the background distribution would influence the N-of-1 sample to take for that gene. We model the expected expression *ŷ*_*g*_ from a new sample as a convex combination of the unobserved expression values across the datasets.

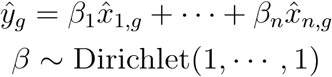

Note that *β* is shared across all genes. A Dirichlet distribution was chosen for *β* to enforce the convexity constraint. Finally, we model the observed expression of the new sample *y*_*g*_, which adds Laplacian error to the expected expression : *ŷ*_*g*_

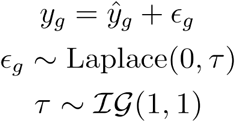

The distribution of the model error *ϵ* is shared between genes and incorporates uncertainty into the posterior to account for variance generated by a poor match of the N-of-1 to any particular background group, weak model fitting, and biological and technical noise. We use a Laplace distribution instead of the more conventional normal because we are interested in identifying expression outliers. The Laplace distribution is heavier-tailed, so it will fit to outliers less aggressively and thereby preserve their outlier status.

This model fits the data well for most cases. However, it behaves poorly on genes that have large variances in the background dataset. The reason is that the *ϵ* parameter has a uniform scale across genes. This causes expression outliers that are modest relative to the variance appear to be more significant. To address this limitation, we normalize the background datasets for variance, but not for location. This normalization step must be incorporated into the model specification, since it is not known *a priori* which background datasets the model will learn to be important, and different background datasets have different variances. This leads to the following equation:

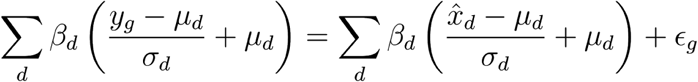

which simplifies to

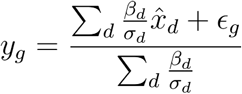

The model can be explored using Markov chain Monte Carlo (MCMC) to obtain samples for *y*_*g*_ that approximate its posterior distribution. If we have an observed expression value for a gene of interest (from the N-of-1 cancer sample), we can compare it to the sampled values. The proportion of sampled values for *y*_*g*_ that are greater (or lesser) than the observed value is an estimate of the posterior predictive p-value for this expression value. The posterior predictive p-value can be seen as a measure of how much of an outlier the expression is given the expectations of the comparison set.

The model is implemented in PyMC3 and each N-of-1 sample is trained using the No-U-turn MCMC sampler ^29,30^. Due to the computational burden of sampling from the model, we employ a couple computational tricks to reduce runtime. First, we integrate out the *µ*_*d,g*_ and *σ*_*d,g*_ parameters so that each 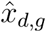 is distributed according to a posterior predictive Student-t. Given our choice of a Dirichlet distribution for *β*, most of the background datasets are assigned 0 weight which means it is inefficient to include all *n* background datasets for every training run. Instead, background datasets are heuristically ranked for similarity to the N-of-1 sample by a combination of analysis of variance (ANOVA) and pairwise distance, and then iteratively added until the posterior predictive p-values converge to Pearson correlation > 0.99 **(S6: Background Dataset Selection)**. The model is available as a Python package for convenience, a Docker container for reproducibility, and a Toil workflow for scalability. The software also provides comprehensive output to aid users in interpreting model results **(S10: Software Engineering)**.

## Results

### TCGA & GTEx Validation

We ran the model on 977 TCGA tumor samples — spanning ten different tissues that had corresponding normals in The Genotype-Tissue Expression Consortium (GTEx) — using normals from GTEx and TCGA as different background datasets **(Figure 2, S2: Assignment of Model Weight to Background Datasets)** ^31^. For every group of samples within a tissue type, the matched tissue in GTEx or TCGA-normal was afforded a majority of the model weight with only two groups of samples receiving less than 60% of all total weight: bladder and stomach. Dimensionality reduction reveals that bladder and stomach samples tend to cluster near other tissue groups that the model assigns weight to **(S2.1.1: Dimensionality Reduction of Low-Weight Samples)**. Despite GTEx and TCGA being independent projects and with no attempts to correct for batch effects, the model identifies the corresponding tissue for most samples.

**Figure 2.**
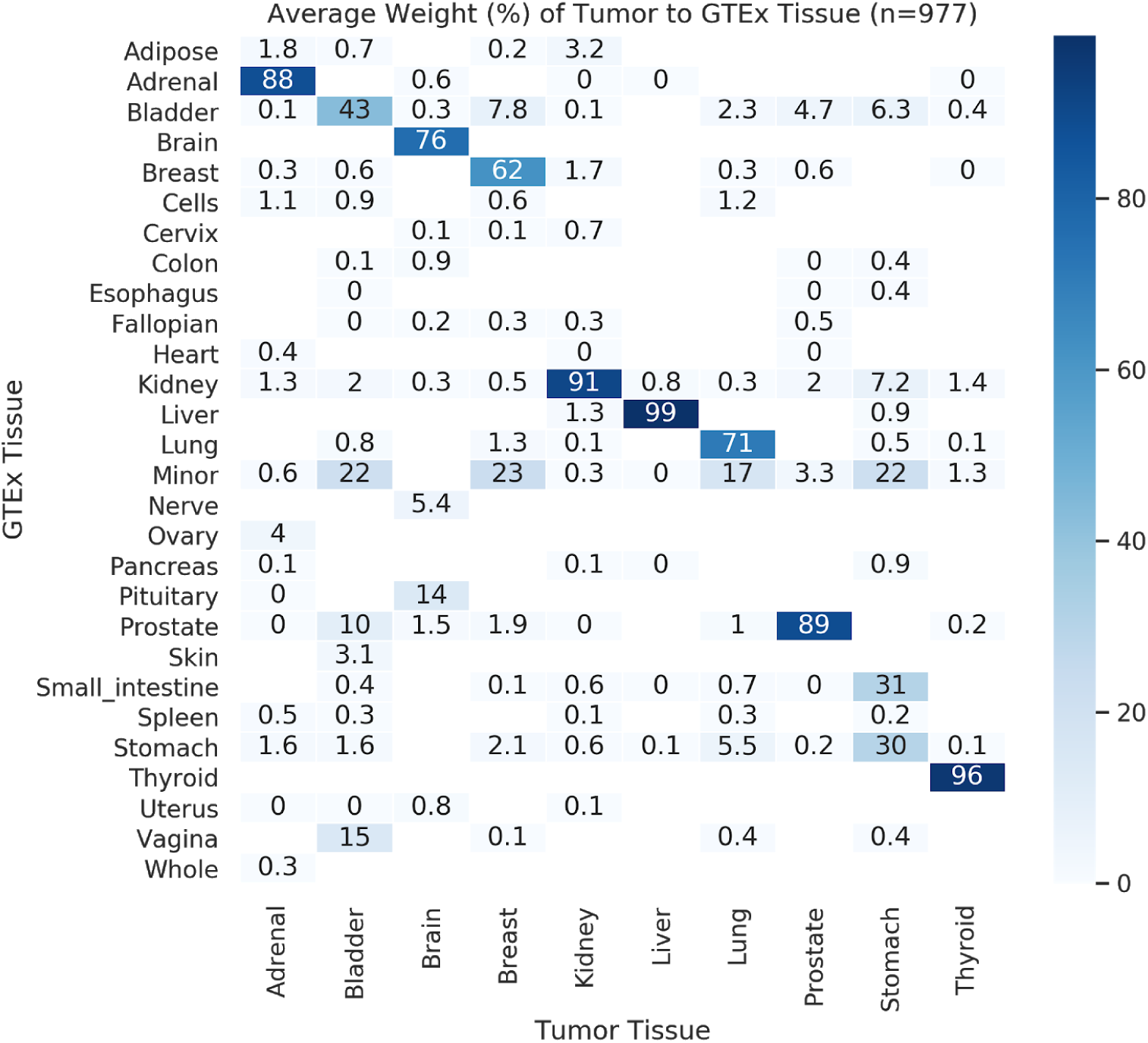
HeatMap of the average model weight assigned to GTEx tissues across tumor subtypes in TCGA. The model assigns a majority of weight to the matched tissue in GTEx for every tumor subtype. Only two tumor tissues, bladder and stomach, received less than 60% of the average model weight. GTEx has only 9 bladder samples and PCA shows those bladder samples cluster on top of minor salivary gland and vagina samples which helps explain its lower average weight relative to other tumor types **(Supplement: Dimensionality Reductions of Low-Weight Samples)**.

### Negative Control

As a negative control experiment, we ran 100 samples across ten GTEx tissues using three different backgrounds: TCGA tumor, TCGA normal, and GTEx. Our expectation was that there would be relatively few outliers when either normal (non-cancer) dataset is used as the background comparison set relative to TCGA-tumor. **Figure 3A** shows that when either GTEx or TCGA-normals were used as the background dataset, the gene p-values shrink towards the middle and outliers are rarely identified. The model tends to assign almost all weight to the N-of-1’s matched tissue in GTEx or TCGA-normal, and the N-of-1 does not deviate significantly from other samples in that tissue group, with few exceptions.

### Testing Model Robustness by Removing Matched Tissues

In most cases, our method is robust to situations in which there is no obvious matched normal tissue **(Figure 3B)**. To demonstrate this, we used our method with a comparison dataset in which we artificially removed the tissue matched to the sample, and then we compared the results with the restricted dataset to the results we obtain with the full dataset. The model will often go from assigning almost all of the weight to the matched normal tissue to distributing it among several other phenotypically similar tissues. However, in most tissues the model largely compensates for the missing data in the final results: the p-values remain highly correlated to those produced with the full dataset **(S4: Effect of Removing Matched Normal on Gene P-values)**. That said, the p-values do move slightly away from the tails, indicating lower power to detect outliers.

**Figure 3.**
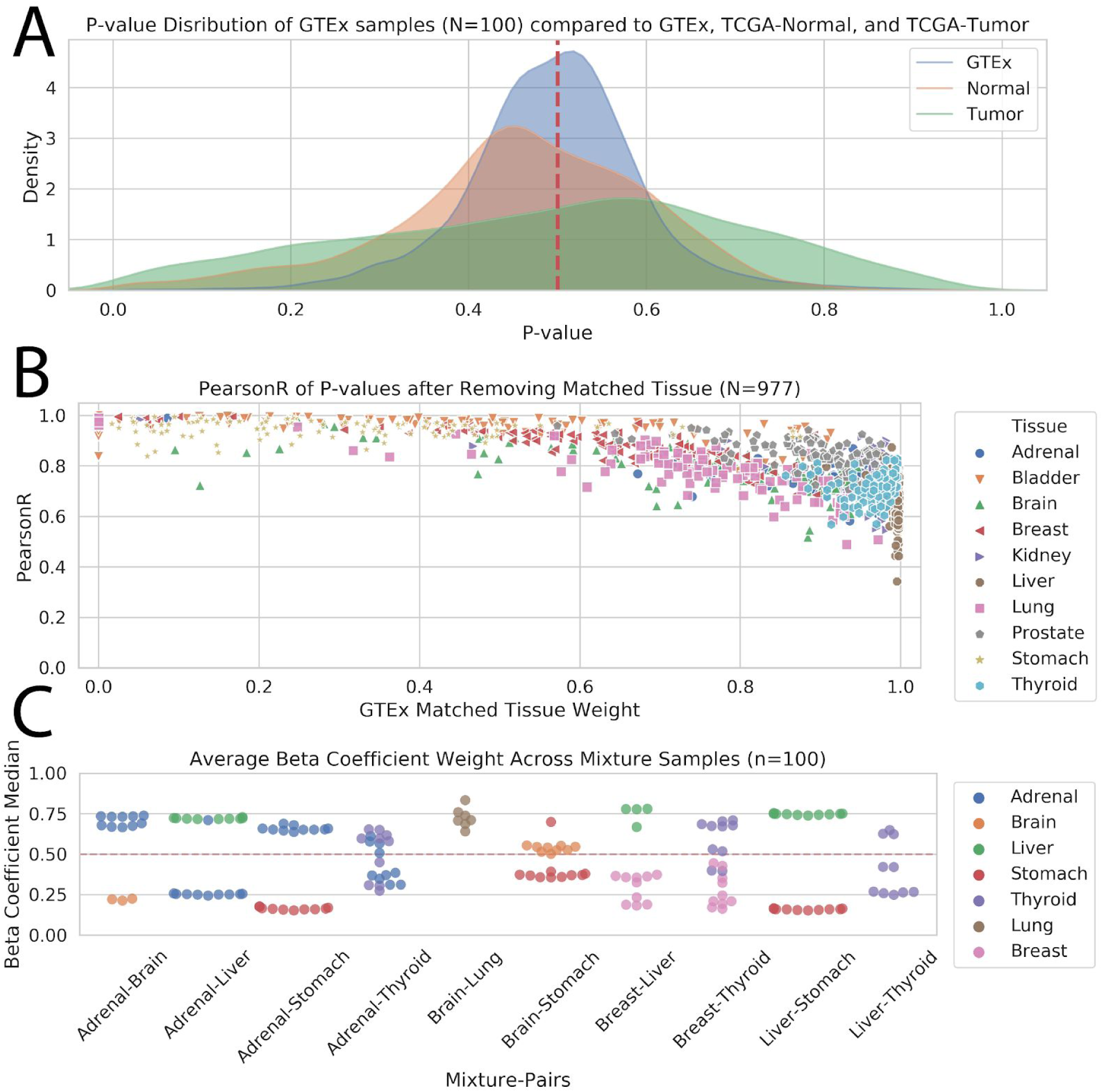
Different aspects of model robustness. **(A)** *Robustness to false positives.*Negative control experiment where 100 GTEx samples were run using GTEx, TCGA-normal, and TCGA-tumor as different backgrounds. Since posterior predictive p-values measure how different the observed result is from model expectations, we assume that using normal tissues as backgrounds for GTEx N-of-1 samples will result in a peak around 0.5 and very few outliers compared to when TCGA-Tumor is used as a background. **(B)** *Robustness to imperfect comparison sets.*The effect that removing a matched tissue has on the gene p-values the model generates as measured by Pearson correlation. The x-axis is the weight assigned to the matched tissue by the model. Gene p-values are relatively consistent even when a matched tissue is removed, particularly if the model can redistribute that weight to tissues of similar phenotypes. **(C)** *Robustness to mixed lineage samples.*Average model weight of mixture samples generated from random pairings of GTEx tissues. Samples were generated by averaging gene expression between randomly sampled subsets of each tissue group. In most cases, the two tissues used to generate the mixture are assigned the majority of the model weight, which is the expected result. Three sets of mixture samples — Adrenal-Brain, Brain-Lung, and Liver-Thyroid — do not get the same result. PCA of those mixture samples, in the context of similar tissues, shows the generated mixture samples happen to cluster closer to other tissues than one or both of the tissues used to generate the samples **(S3.7: Dimensionality Reductions of Mismatched Mixture Samples)**. In these circumstances, we expect the model to assign weight to those tissues that are more similar to the mixture samples.

### Mixture Simulation

We used a simulation to validate the method’s ability to identify comparison sets in tumors of non-specific lineage. Simulated N-of-1 samples were created by randomly selecting tissue pairs from GTEx then averaging gene expression between random samples from those tissue pairs. PCA of the mixture samples show a tight cluster in between the two clusters for the contributing tissues **(S3: Mixture Simulations)**. Mixture samples were run through the model and the weights from the two contributing tissues were collected **(Figure 3C).** Ideally, 50% of the model weight should be assigned to each of the contributing tissues used to generate those mixture samples, which is true for a majority of the tissue pairs. We would not want the model to split weight evenly between the two contributing tissues if the generated mixtures happen to be more similar to other tissue types in the background dataset. For mixture samples that did not match to the tissues used to generate them, dimensionality reduction clearly shows that other tissues happen to cluster closer to the mixtures than one or both of the contributing tissues **(S3.7: Dimensionality Reductions of Mismatched Mixture Samples)**.

### Upregulated Gene Outlier Counts Across Tumor Subtypes

Gene amplification and overexpression are a common hallmark of cancer cells, resulting from extra copies of a locus (amplicon) as well as other genetic and epigenetic changes. In many cases, these changes occur in genes that are specific to their tissue of origin ^32–34^. Many of these commonly mutated genes can be targeted by existing drugs ^35^. Eighty-five such druggable genes, mostly receptor tyrosine kinases, were curated and provided to us by Treehouse. We calculated p-values for these genes using our method across the 977 TCGA samples (**Figure 4)**. Genes with p-values below a cutoff (< 0.05) that also appeared in more than a third of the tumor samples within a subtype were all identified as known biomarkers in the literature (**S5.3: Literature Corroboration of Outlier Findings).** These include AURKA in both bladder and breast cancer ^36–38^, AURKB, CDK4, EGFR, and PDGFRA in brain cancers and gliomas ^39–43^, MET in thyroid carcinomas and gastric cancers ^44–47^, and ROS1 in lung cancers ^48^.

**Figure 4.**
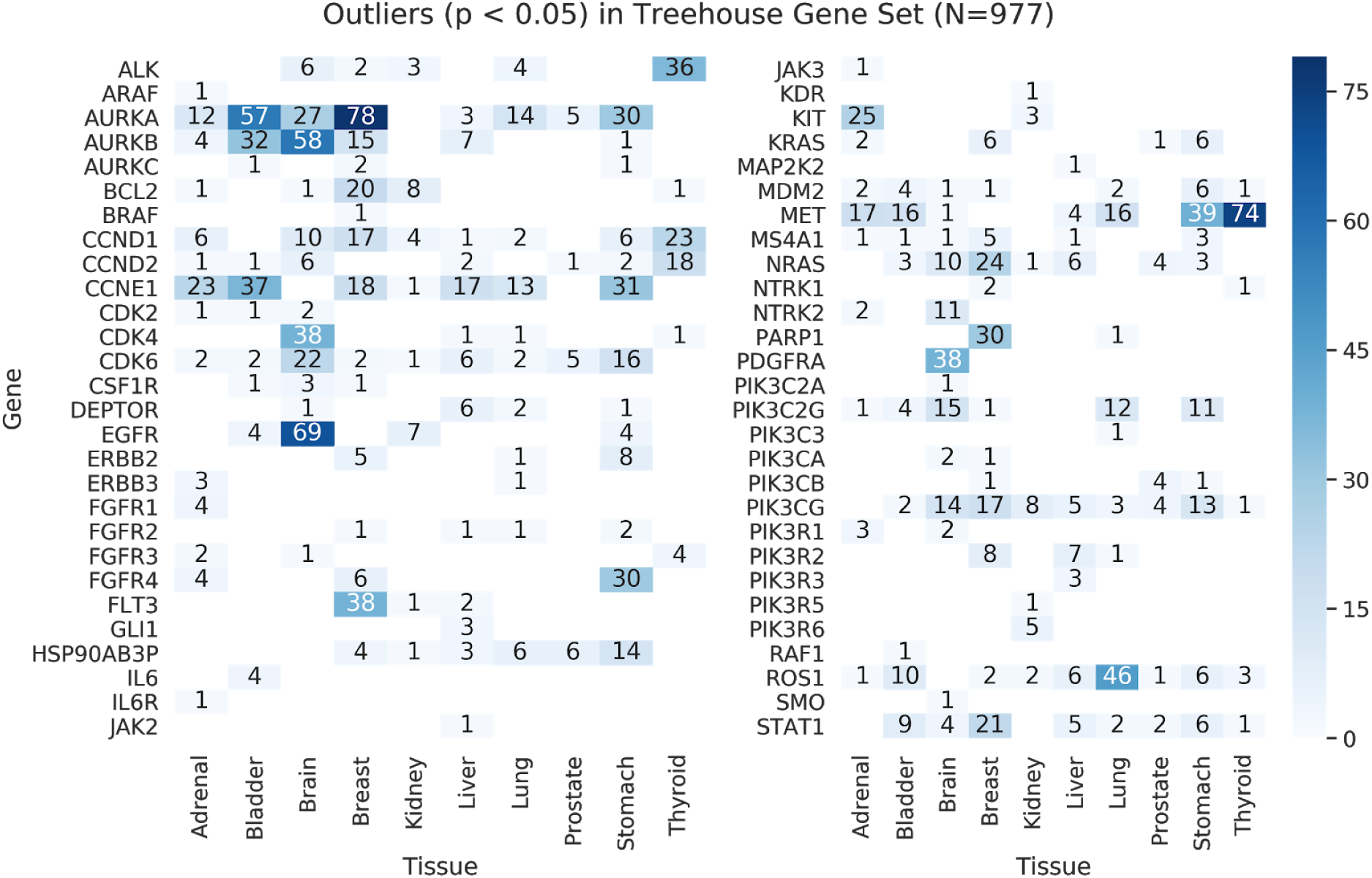
Outlier counts given a p-value cutoff of 0.05. Eighty-five “druggable” genes curated by Treehouse were used as the target gene set for this analysis. All genes with counts for more than half of the samples within a tumor subtype were identified as known tumor biomarkers within that subtype in the literature (**S5.3: Literature Corroboration of Outlier Findings).**

### Exploring Results For a Single Sample

To illustrate how our method is used in practice, we demonstrate the model on a single sample rather than summary statistics over many samples. **Figure 5** compares our method on a random tumor sample from TCGA to Treehouse’s standard practice approach of pooling normal samples and applying a cutoff based on inner-quartile range on a selection of 85 cancer genes which could be targeted by an available therapy ^49,50^. The random sample is labeled thyroid carcinoma in TCGA. Over 8,000 samples from GTEx dataset were used as the comparative normal dataset, categorized by tissue type. The model automatically weights each tissue group and assigns a majority of the weight to thyroid tissue in GTEx. Where the pan-normal cutoff method returns a binary classification for each of the selected genes, our method returns a posterior predictive p-value generated from a distribution informed by the background datasets that are most similar to the N-of-1 sample. Where there is disagreement in outlier classification between the two methods (PIK3CB and CCND2), the posterior distribution can be examined in the context of the highest weighted background dataset(s) to clearly understand how the p-value was generated. For example, our method does not identify PIK3CB as an outlier (given a p-value cutoff of 0.05), because the method down-weights non-thyroid tissues, which have lower average expression for this gene than normal thyroid tissue.

**Figure 5.**
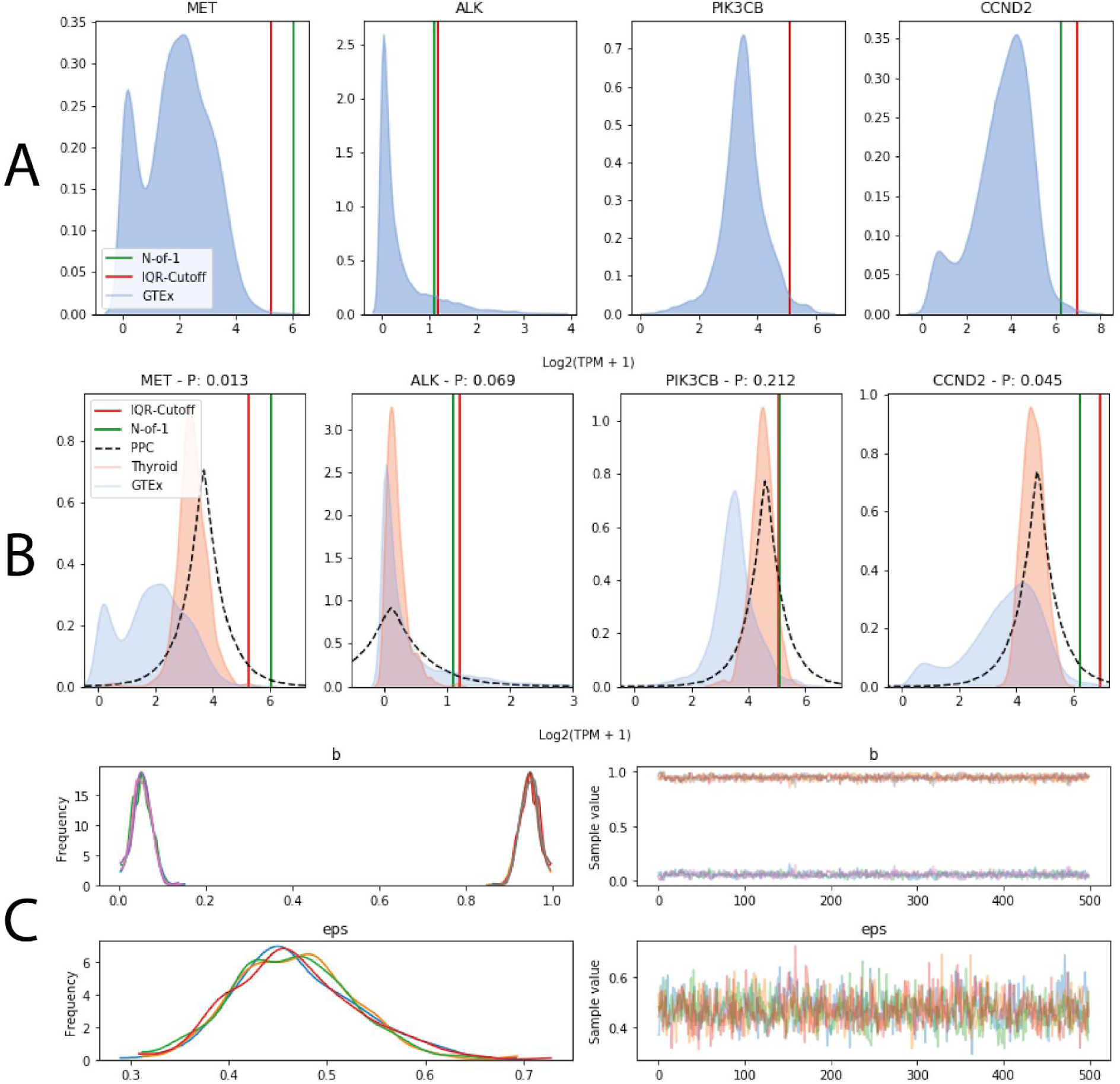
Comparing results between a pan-normal cutoff approach and our model for a single sample. (**A**) Gene expression distributions for 4 genes using the GTEx dataset, overlaid with the gene expression value from our N-of-1 tumor in green and an outlier cutoff in red generated from Tukey’s method (Q3 + IQR * 1.5). (**B**) Same plots as (A) along with the gene expression distribution for Thyroid in GTEx (orange) and the posterior predictive distribution (dotted black). The posterior distribution closely matches the GTEx Thyroid distribution because the model assigned almost all of the weight to that tissue. Posterior predictive p-values for each gene were added to the plot titles which is based on the relative proportion of samples in the posterior distribution greater than the value from the N-of-1 sample. (**C**) Trace plots for *β* and *ϵ* from the MCMC sampling step. *β* is a Dirichlet distribution and sums to 1 and in this case assigned essentially all model weight to Thyroid in GTEx.

## Discussion

### Method Contribution

Our method avoids selection bias introduced by having to choose a single comparison background dataset. It also provides continuous p-values for genes that can be ranked and also avoid missing borderline cases that would be ignored by existing cutoff methods. Moreover, in addition to under- and overexpression, the model quantifies the similarity of the analyst’s sample to background comparison sets. Researchers can use this feature as a diagnostic — diffuse weight distributions suggest that the model did not identify a strong set of matches among the background datasets. The model has also been demonstrated to be robust to false positives, incomplete comparison datasets, and when analyzing samples of mixed lineage.

These benefits do come with a tradeoff in computation — calculating outliers via other methods can be very fast whereas the runtime of this method quadratically increases with the number of genes and datasets. The method is only appropriate for analyzing small targeted gene sets — fewer than ∼200 ideally — due to the way model complexity scales. After ∼200 genes it is better to parallelize multiple runs for a single sample, which is facilitated by a Toil-based version of the workflow that makes scaling trivial and allows the method to run hundreds of samples in parallel on both standard local and cloud-based clusters ^51^. The software provides intermediate output at every step in the workflow so users can validate model convergence, assess the model’s similarity metrics for background datasets to the N-of-1, and examine every model parameter to reproduce how p-values were calculated for every gene.

### Limitations

The model makes certain mathematical assumptions that we know to be unrealistic. First, it implicitly assumes independence between the N-of-1 sample’s genes. It also assumes that the sample’s gene expression can be approximated by a convex mixture of the gene expression of the background datasets, which aids interpretability at the expense of descriptive accuracy.

These limitations could be addressed by extending the model. To incorporate correlation between genes, random variables for each *x*_*d,g*_ could be replaced with multivariate distributions that are shared between all genes belonging to that group of coexpressed genes, where groups could be formed through a clustering process or according to existing annotations. Prior knowledge could also be used to introduce non-independence between genes into the model. For instance, the independent error terms could be replaced by errors that are structured according to the Laplacian of a gene interaction network ^52^.

### Extensibility to Single-Cell RNA-seq

Our model could be theoretically extended to single cell RNA-seq analysis in addition to bulk. However, without any modifications to the existing model, this would require training the model for each cell, which would be computationally expensive. A faster alternative would be to cluster the cells and sample a small number of representative cells from each cluster to run through the model to get summary information about each of the single-cell clusters. However, the high levels of technical noise, biological variability between cells, and dropouts, may require too many training genes to obtain robust estimates of the model parameters ^53^. Finally, the distribution of the random variable *x*_*d,g*_ would need to be replaced with a more appropriate distribution to model single-cell expression, such as a beta-Poisson mixture model ^54^.

## Conclusions

As clinicians have begun to demonstrate that RNA-seq analysis can produce actionable findings for cancer patients, it is necessary to have informed and principled analytic tools for an individual patient. The method we have proposed detects gene expression outliers among a panel of target genes. It also provides additional information for researchers to explore and validate the results through examination of the model’s parameters. For portability, scalability, and reproducibility, we have made this open-source tool available as a Python package, Docker container, and Toil workflow available at https://github.com/jvivian/gene-outlier-detection/.

## Declarations

### Ethics approval and consent to participate

Not applicable

### Consent for publication

Not applicable

### Availability of data and material

- Raw expression data used in these experiments is available at UCSC Xena
  - https://toil.xenahubs.net
- All data used to produce every figure and experiment is publicly available on UCSC
  - http://courtyard.gi.ucsc.edu/~jvivian/outlier-paper/
- Our model’s code is open-source and available on GitHub
  - https://github.com/jvivian/gene-outlier-detection

## Authors’ contributions

JV, JE, and BP wrote the manuscript and developed the method. JV wrote the software implementation, documentation, and supplement. HB, OMV, and BP oversaw project details.

## Acknowledgements

We thank the members of the Computational Genomics Laboratory at the UC Santa Cruz Genomics Institute for numerous conversations that helped shape this work as well as the Treehouse organization for their collaborative efforts.

