## Supplement for "A Bayesian Framework for Detecting Gene Expression Outliers in Individual Samples"

John Vivian

June 2019

### 1 Introduction

For reproducibility, we have included all code used to generate figures in the paper here along with additional experiments (section 8, section 9). We have also hosted all data used in this paper at a public URL `http://courtyard.gi.ucsc.edu/~jvivian/outlier-paper/`. The **expression-matrices** subdirectory contains HD5 matrices with metadata and gene expression information for GTEx, TCGA tumors, and TCGA normals. In Python, these matrices can be opened with Pandas, e.g. (`df = pd.read_hdf('tumor.hd5')`). The **experiments** subdirectory contains compressed archives of data generated for figures in the paper as well as the supplement. Each section in the supplement explictly references a URL to the data used to generate figures in that section.

| Tissue | Tumor Subtype | Number of Samples |
|---|---|---|
| Adrenal | Adrenocortical Carcrinoma | 77 |
| Bladder | Bladder Urothelial Carcinoma | 100 |
| Brain | Brain Lower Grade Glioma | 100 |
| Breast | Breast Invasive Carcinoma | 100 |
| Kidney | Kidney Renal Clear Cell Carcinoma | 100 |
| Liver | Liver Hepatocellular Carcinoma | 100 |
| Lung | Lung Adenocarcinoma | 100 |
| Prostate | Prostate Adenocarcinoma | 100 |
| Stomach | Stomach Adenocarcinoma | 100 |
| Thyroid | Thyroid Carcinoma | 100 |

Table 1: Tissues and corresponding tumor subtypes for the 977 samples used to generate data for section 2 and section 5. 100 samples were chosen randomly from each subtype, except for adrenocortical carcinoma which only has 77 samples.

#### 2 Assignment of Model Weight to Background Datasets

This section explores assignment of model weight to different background datasets in GTEx across ~1,000 TCGA tumor samples. Despite no attempt to correct for batch effects and GTEx and TCGA being completely separate projects with different laboratory protocols, the model is able to accurately assign model weight to the appropriate tissue in GTEx for a majority of samples across ten different tumor types.

All data used to generate the following section is avilable at http://courtyard.gi.ucsc.edu/~jvivian/outlier-paper/experiments/gtex-1000.tar.gz.

```
In [1]: import os

        import matplotlib.pyplot as plt
        import pandas as pd
        import seaborn as sns
        from sklearn.decomposition import pca

        sns.set(font_scale=1.0)
        sns.set_style("whitegrid")

        class Weights:
            def __init__(self, tumor_path: pd.DataFrame, sample_dir: str):
                self.tumor_path = tumor_path
                self.tumor = self._load_tumor()
                self.genes = self.tumor.columns[5:]
                self.sample_dir = sample_dir
                self.df = self._weight_df()
                self.tissue_perc = self._perc_df()
                self.subtype_perc = self._perc_df("subtype")
                self.num_samples = len(self.df["sample"].unique())

            def _weight_df(self) -> pd.DataFrame:
                """
```

```python
        Creates DataFrame of sample weights from a directory of samples
        Columns: tissue, normal_tissue, weight, sample_id

        Returns:
            DataFrame of Weights
        """
        # DataFrame: cols=tissue, normal_tissue, weight
        weights = []
        tissues = self.tumor.tissue
        subtypes = self.tumor.subtype
        for sample in os.listdir(self.sample_dir):
            sample_tissue = tissues.loc[sample]
            sample_subtype = subtypes.loc[sample]
            w = pd.read_csv(
                os.path.join(self.sample_dir, sample, "weights.tsv"), sep="\t"
            )
            w.columns = ["normal_tissue", "Median", "std"]
            w["tissue"] = sample_tissue
            w["subtype"] = sample_subtype
            w["sample"] = sample
            # Add zero weight if missing
            if sample_tissue not in w.normal_tissue.values:
                w.loc[len(w)] = [
                    sample_tissue,
                    0,
                    0,
                    sample_tissue,
                    sample_subtype,
                    sample,
                ]
            weights.append(w.drop("std", axis=1))

        return pd.concat(weights).reset_index(drop=True)

    def _perc_df(self, group="tissue") -> pd.DataFrame:
        """
        Converts DataFrame of weights into a DataFrame of percentages

        Returns:
            Weight percentage DataFrame
        """
        c = self.df.groupby([group, "normal_tissue"])["Median"].sum().rename("count")
        perc = c / c.groupby(level=0).sum() * 100
        return perc.reset_index()

    def _load_tumor(self):
        print(f"Reading in {self.tumor_path}")
        if self.tumor_path.endswith(".csv"):
            df = pd.read_csv(self.tumor_path, index_col=0)
        elif self.tumor_path.endswith(".tsv"):
            df = pd.read_csv(self.tumor_path, sep="\t", index_col=0)
        else:
            try:
                df = pd.read_hdf(self.tumor_path)
```

```
        except Exception as e:
            print(e)
            raise RuntimeError(f"Failed to open DataFrame: {self.tumor_path}")
    return df

def plot_match_scatter(self, out_dir: str = None):
    """
    Scatterplot of samples by tissue and their matched tissue model weight

    Args:
        out_dir: Optional output directory

    Returns:
        Plot axes object
    """
    df = self.df
    # Subset for matched-tissue samples
    df = df[df.normal_tissue == df.tissue].sort_values("tissue")

    f, ax = plt.subplots(figsize=(12, 4))
    sns.swarmplot(data=df, x="tissue", y="Median")
    plt.xticks(rotation=45)
    plt.xlabel("Tissue")
    plt.ylabel("GTEx Matched Tissue Weight")
    plt.title("TCGA Tumor Samples and Model Weight for GTEx Matched Tissue")
    if out_dir:
        plt.savefig(os.path.join(out_dir, "matched_weight_scatter.svg"))
    return ax

def plot_perc_heatmap(self, out_dir: str = None, subtype=False):
    """
    Heatmap of weight percentages

    Args:
        out_dir: Optional output directory

    Returns:
        Plot axes object
    """
    df = self.subtype_perc if subtype else self.tissue_perc
    column = "subtype" if subtype else "tissue"
    f, ax = plt.subplots(figsize=(12, 7))
    perc_heat = df.pivot(index="normal_tissue", columns=column, values="count")
    sns.heatmap(
        perc_heat.apply(lambda x: round(x, 1)),
        cmap="Blues",
        annot=True,
        linewidths=0.5,
    )
    plt.xlabel("Tumor Tissue")
    plt.ylabel("GTEx Tissue")
    title = f"Average Weight (%) of Tumor to GTEx Tissue (n={self.num_samples})"
    plt.title(title)
    if out_dir:
```

```
            plt.savefig(os.path.join(out_dir, "weight_perc_heatmap.svg"))
        return ax

    def plot_pca_nearby_tissues(self, background_path: str, tissues, tumor_tissue):
        """
        PCA of nearby tissues

        Args:
            background_path: Path to background dataframe
            tissues: Tissues to include in PCA
            tumor_tissue: Label for tumor tissue

        Returns:
            Plot axes object
        """
        df = pd.read_hdf(background_path)
        tumor = self.tumor
        tumor = tumor[tumor.tissue == tumor_tissue]
        tumor["tissue"] = f"{tumor_tissue}-Tumor"

        sub = df[df.tissue.isin(tissues)]
        pca_df = pd.concat([tumor, sub]).dropna(axis=1)
        embedding = pca.PCA(n_components=2).fit_transform(pca_df[self.genes])
        embedding = pd.DataFrame(embedding)
        embedding.columns = ["PCA1", "PCA2"]
        embedding["tissue"] = list(pca_df["tissue"])

        f, ax = plt.subplots(figsize=(8, 8))
        sns.scatterplot(
            data=embedding, x="PCA1", y="PCA2", hue="tissue", style="tissue"
        )
        plt.title(f"PCA of {tumor_tissue} and Nearby GTEx Tissues")
        return ax
```

#### 2.1 Weight plots for GTEx as background dataset

```
In [2]: w = Weights(
            tumor_path='/mnt/data/expression/tumor.hd5',
            sample_dir='/mnt/normsd-outlier-runs/gtex-1000/'
        )
```

```
Reading in /mnt/data/expression/tumor.hd5
```

```
In [3]: out_dir = '/mnt/figures/defense-figures'
        w.plot_match_scatter();
        plt.tight_layout()
        plt.savefig(
            os.path.join(out_dir, 'TCGA-Samples-and-Model-Weight.png'),
            dpi=300,
            transparent=True
        )
```

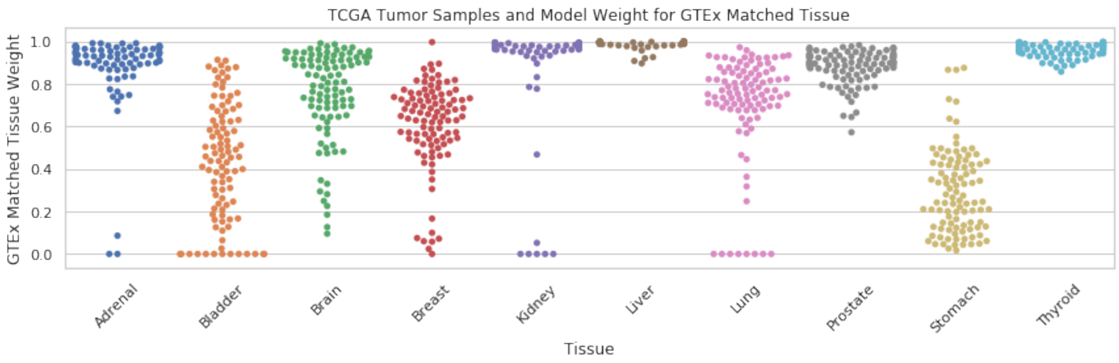

```
In [4]: w.plot_perc_heatmap();
        plt.tight_layout()
        plt.savefig(
            os.path.join(out_dir, 'GTEx-1000-weight-heatmap.png'),
            dpi=300,
            transparent=True
        )
```

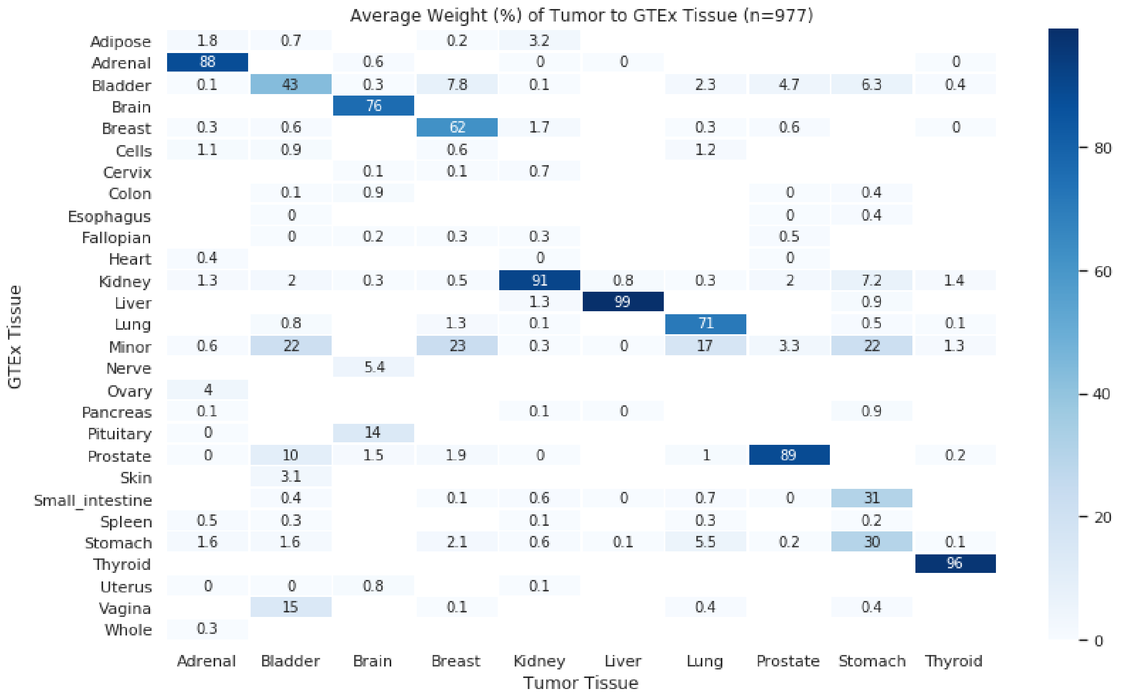

##### 2.1.1 Dimensionality Reduction of Low-Weight Samples

This section explores low model weight assignment to bladder and stomach samples. The subsections below show dimensionality reduction plots of samples from two tissues – bladder and stomach – that the model had difficulty assigning the correct weight to. Since the model assigns weights based on tissues that are most similar to the N-of-1 sample, it can only perform as well as the data provided. Performing PCA of bladder tumor samples alongside tissues the model assigned weight to reveal why the model has difficulty associating the normal bladder tissue in GTEx to the bladder tumor samples. GTEx possesses only 9 bladder samples and these samples cluster into two distinct groups – one near the bladder tumor samples and the other near vagina and minor salivary gland samples, which are the other two groups that were assigned a significant percentage of weight. Finally, it is apparent why the model struggled to correctly assign GTEx stomach as the primary weight for stomach cancer samples. GTEx stomach samples are split into two sub-populations and small intestine, which is proximal to and phenotypically similar to stomach tissue, clusters closely to stomach tumor samples.

**PCA of Bladder**

```
In [6]: w.plot_pca_nearby_tissues(
            background_path='/mnt/data/outlier/gtex.hd5',
            tissues=['Bladder', 'Minor', 'Vagina'],
            tumor_tissue='Bladder'
        );
        plt.tight_layout()
        plt.savefig(os.path.join(out_dir, 'PCA-Bladder.png'), dpi=300, transparent=True)
```

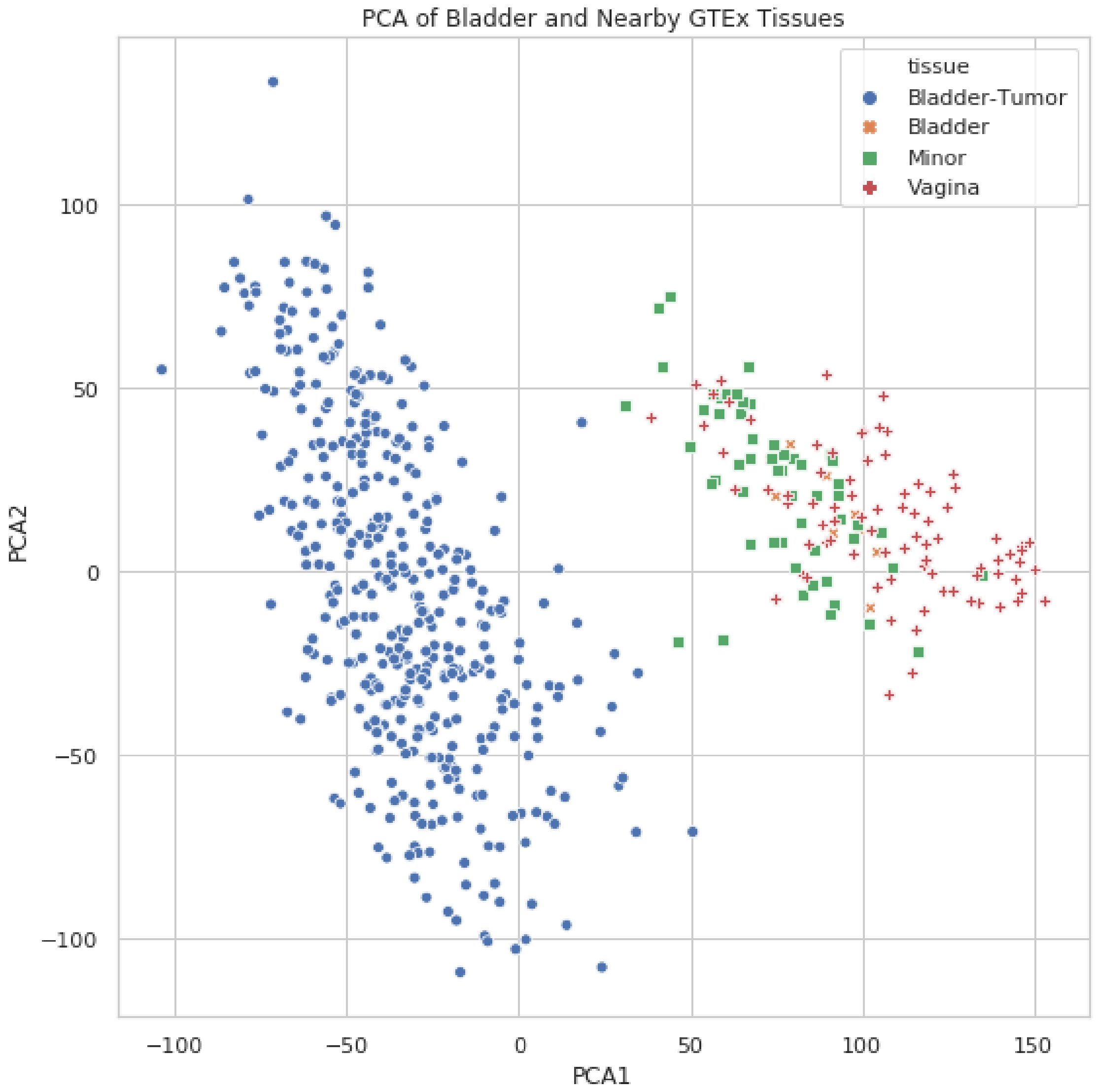

**PCA of Stomach**

```
In [7]: w.plot_pca_nearby_tissues(
            background_path='/mnt/data/outlier/gtex.hd5',
            tissues=['Stomach', 'Small_intestine', 'Minor'],
            tumor_tissue='Stomach'
        );
        plt.tight_layout()
        plt.savefig(os.path.join(out_dir, 'PCA-Stomach.png'), dpi=300, transparent=True)
```

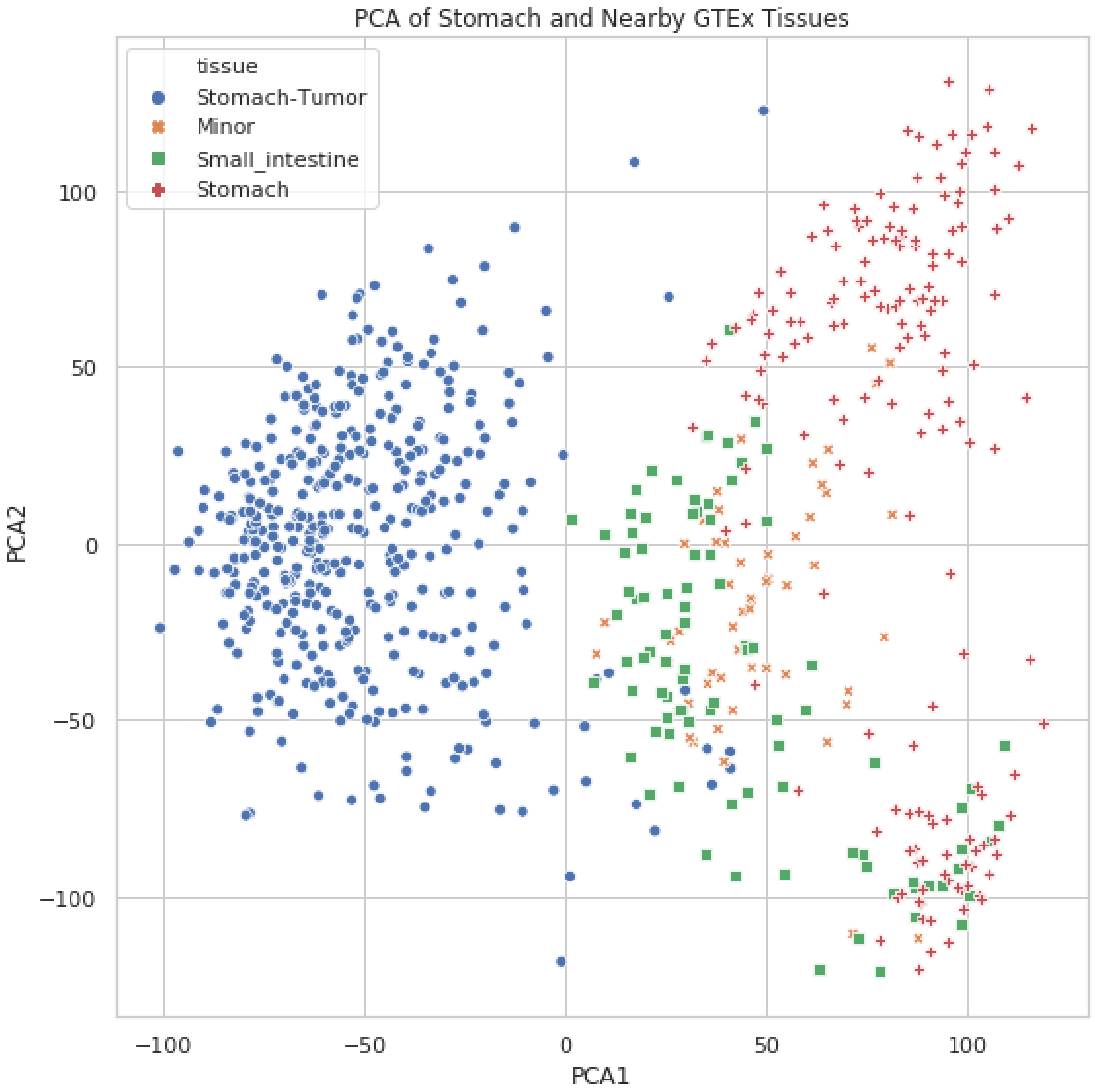

#### 3 Mixture Simulations

We validate the model's ability to assign weights to samples of mixed phenotypes by simulating mixture samples from pairs of different tissues and then checking if the model splits the weights between those two contributing tissues.

All data used to generate the following section is avilable at http://courtyard.gi.ucsc.edu/~jvivian/outlier-paper/experiments/mixture-simulation.tar.gz. This archive contains both the results from running the model on the mixture samples as well as the simulated matrices.

In [16]:
```
import os
from itertools import combinations

import matplotlib.pyplot as plt
import numpy as np
```

```python
import pandas as pd
import seaborn as sns
from sklearn.decomposition import pca

sns.set_style("whitegrid")

class Mixture:
    def __init__(self, df_path):
        self.df_path = df_path
        self.df = self._load_df(df_path)
        self.genes = list(self.df.columns[5:])

    @staticmethod
    def _load_df(path):
        """Loads DataFrame"""
        print(f"Reading in {path}")
        if path.endswith(".csv"):
            df = pd.read_csv(path, index_col=0)
        elif path.endswith(".tsv"):
            df = pd.read_csv(path, sep="\t", index_col=0)
        else:
            try:
                df = pd.read_hdf(path)
            except Exception as e:
                print(e)
                raise RuntimeError(f"Failed to open DataFrame: {path}")
        return df

    def select_mixture_tissues(self, number_of_pairs=10):
        """Generates N tissue pairs"""
        # Choose a tissue subset with relevant matches to TCGA
        tissues_to_include = [
            "Adrenal",
            "Bladder",
            "Brain",
            "Breast",
            "Kidney",
            "Liver",
            "Lung",
            "Prostate",
            "Stomach",
            "Thyroid",
        ]
        df = self.df[self.df.tissue.isin(tissues_to_include)]
        df = df.groupby("tissue").filter(lambda x: len(x) > 100)
        tissues = list(combinations(df.tissue.unique(), 2))
        ix = np.random.choice(range(len(tissues)), number_of_pairs, replace=False)

        pairs = []
        for i in ix:
            pairs.append(tissues[i])
        return pairs
```

```python
def mixture_matrix(self, t1, t2, out_dir):
    """Create mixture_matrix for one tissue pair"""
    df = self.df
    df1 = df[df.tissue == t1]
    df2 = df[df.tissue == t2]
    path = os.path.join(out_dir, f"{t1}-{t2}.hd5")
    if os.path.exists(path):
        return pd.read_hdf(path)

    samples = []
    for i in range(50):
        s1 = np.random.choice(df1.index, 100, replace=False)
        s2 = np.random.choice(df2.index, 100, replace=False)

        g1 = df1.loc[s1]
        g2 = df2.loc[s2]

        vec = pd.concat([g1, g2])[self.genes].median()
        samples.append(vec)
    mixture_matrix = pd.concat(samples, axis=1).T
    mixture_matrix.index = [f"{t1}-{t2}-{i}" for i in range(50)]

    mixture_matrix.to_hdf(path, key="exp")
    return mixture_matrix

def mixture_matrices(self, tissues, out_dir):
    """Create mixture matrices for all tissue pairs"""
    matrices = {}
    for t1, t2 in tissues:
        label = f"{t1}-{t2}"
        print(f"Creating mixture matrix for {label}")
        matrices[label] = self.mixture_matrix(t1, t2, out_dir)
    return matrices

def manifest(self, tissues, matrix_dir, output_dir):
    """Create manifest for outlier Toil run"""
    path = os.path.join(output_dir, "manifest.tsv")
    with open(path, "w") as f:
        f.write("id\tsample\n")
        for t1, t2 in tissues:
            label = f"{t1}-{t2}"
            for i in range(10):
                matrix_path = os.path.join(matrix_dir, f"{label}.hd5")
                f.write(f"{label}-{i}\t{matrix_path}\n")

@staticmethod
def weight_df(sample_dir: str, only_contributing_tissues=True) -> pd.DataFrame:
    """Creates DataFrame from weights across samples"""
    samples = os.listdir(sample_dir)
    weights = []
    for sample in samples:
        t1, t2 = sample.split("-")[:-1]
        label = f"{t1}-{t2}"
        path = os.path.join(sample_dir, sample, "weights.tsv")
```

```
            w = pd.read_csv(path, sep="\t")
            w.columns = ["tissue", "Median", "std"]
            w["sample"] = sample
            w["label"] = label
            if only_contributing_tissues:
                w = w[w.tissue.isin([t1, t2])]
            weights.append(w.drop("std", axis=1))
        weights = pd.concat(weights).reset_index(drop=True)
        return weights.sort_values("label")

    def plot_mixture_pca(self, matrix_dir: str):
        """PCA of mixture samples and the tissues used to generate them"""
        os.listdir(matrix_dir)
        f, ax = plt.subplots(4, 3, figsize=(12, 4 * 4))
        ax = ax.flatten()
        for i, matrix in enumerate(os.listdir(matrix_dir)):
            t1, t2 = os.path.splitext(matrix)[0].split("-")
            mix_df = pd.read_hdf(os.path.join(matrix_dir, matrix))
            mix_df["tissue"] = "Mixture"
            sub = self.df[self.df.tissue.isin([t1, t2])]
            pca_df = pd.concat([sub, mix_df]).dropna(axis=1)
            embedding = pca.PCA(n_components=2).fit_transform(pca_df[self.genes])
            embedding = pd.DataFrame(embedding)
            embedding.columns = ["PCA1", "PCA2"]
            embedding["tissue"] = list(pca_df["tissue"])
            sns.scatterplot(
                data=embedding,
                x="PCA1",
                y="PCA2",
                hue="tissue",
                style="tissue",
                ax=ax[i],
            )
            ax[i].set_title(f"{t1}-{t2}")
        for i in [10, 11]:
            f.delaxes(ax.flatten()[i])
        plt.tight_layout()
        return ax

    def plot_pca_nearby_tissues(self, matrix_path: str, tissues):
        st_df = pd.read_hdf(matrix_path)
        label = os.path.splitext(os.path.basename(matrix_path))[0]
        st_df["tissue"] = "Mixture"
        sub = self.df[self.df.tissue.isin(tissues)]
        pca_df = pd.concat([st_df, sub]).dropna(axis=1)
        embedding = pca.PCA(n_components=2).fit_transform(pca_df[self.genes])
        embedding = pd.DataFrame(embedding)
        embedding.columns = ["PCA1", "PCA2"]
        embedding["tissue"] = list(pca_df["tissue"])

        f, ax = plt.subplots(figsize=(8, 8))
        sns.scatterplot(
            data=embedding, x="PCA1", y="PCA2", hue="tissue", style="tissue"
        )
```

```
            plt.title(f"PCA of {label} Mixtures and Nearby Tissues")
            return ax

        @staticmethod
        def plot_weight_swarm(wdf: pd.DataFrame):
            """Swarmplot of weights for the tissues that comprise the mixture"""
            f, ax = plt.subplots(figsize=(12, 4))
            sns.swarmplot(data=wdf, x="label", y="Median", hue="tissue", size=8, alpha=0.75)
            plt.ylim([0, 1])
            plt.xticks(rotation=45)
            plt.legend(bbox_to_anchor=(1.01, 1))
            plt.axhline(0.5, c="r", alpha=0.5, ls="--")
            plt.xlabel("Mixture-Pairs")
            plt.ylabel("Beta Coefficient Median")
            plt.title(f"Average Beta Coefficient Weight Across Mixture Samples (n=100)")
            return ax

        @staticmethod
        def plot_weight_boxplot(wdf, label):
            """Boxplot of weigts for a single label"""
            weights = wdf[wdf.label == label]
            f, ax = plt.subplots(figsize=(8, 4))
            sns.boxplot(data=weights, x="tissue", y="Median")
            plt.title(f"Boxplot of Weights for {label} Mixtures")
            plt.xlabel("Tissue")
            plt.ylabel("Average Weight")
            plt.ylim([0, 1])
            return ax
```

```
In [7]: m = Mixture('/mnt/data/outlier/gtex.hd5')
```

```
Reading in /mnt/data/outlier/gtex.hd5
```

##### 3.1 Select Tissue Pairs

```
In [3]: tissues = m.select_mixture_tissues()
        tissues
```

```
Out[3]: [('Adrenal', 'Brain'),
         ('Breast', 'Liver'),
         ('Breast', 'Thyroid'),
         ('Adrenal', 'Stomach'),
         ('Liver', 'Thyroid'),
         ('Brain', 'Lung'),
         ('Adrenal', 'Liver'),
         ('Liver', 'Stomach'),
         ('Adrenal', 'Thyroid'),
         ('Brain', 'Stomach')]
```

##### 3.2 Create Simulated Sample Matrices

```
In [4]: matrix_dir = '/mnt/normsd-outlier-runs/mixture-simulation/matrices'
        matrices = m.mixture_matrices(tissues, matrix_dir)
```

```
Creating mixture matrix for Adrenal-Brain
Creating mixture matrix for Breast-Liver
Creating mixture matrix for Breast-Thyroid
Creating mixture matrix for Adrenal-Stomach
Creating mixture matrix for Liver-Thyroid
Creating mixture matrix for Brain-Lung
Creating mixture matrix for Adrenal-Liver
Creating mixture matrix for Liver-Stomach
Creating mixture matrix for Adrenal-Thyroid
Creating mixture matrix for Brain-Stomach
```

##### 3.3 PCA of Mixture Samples

Run PCA on all mixture samples and the two tissues used to generate the mixture samples. Since this is PCA, the mixture samples should be situated approximately between the clusters of the two contributing tissues.

```
In [5]: m.plot_mixture_pca(matrix_dir);
```

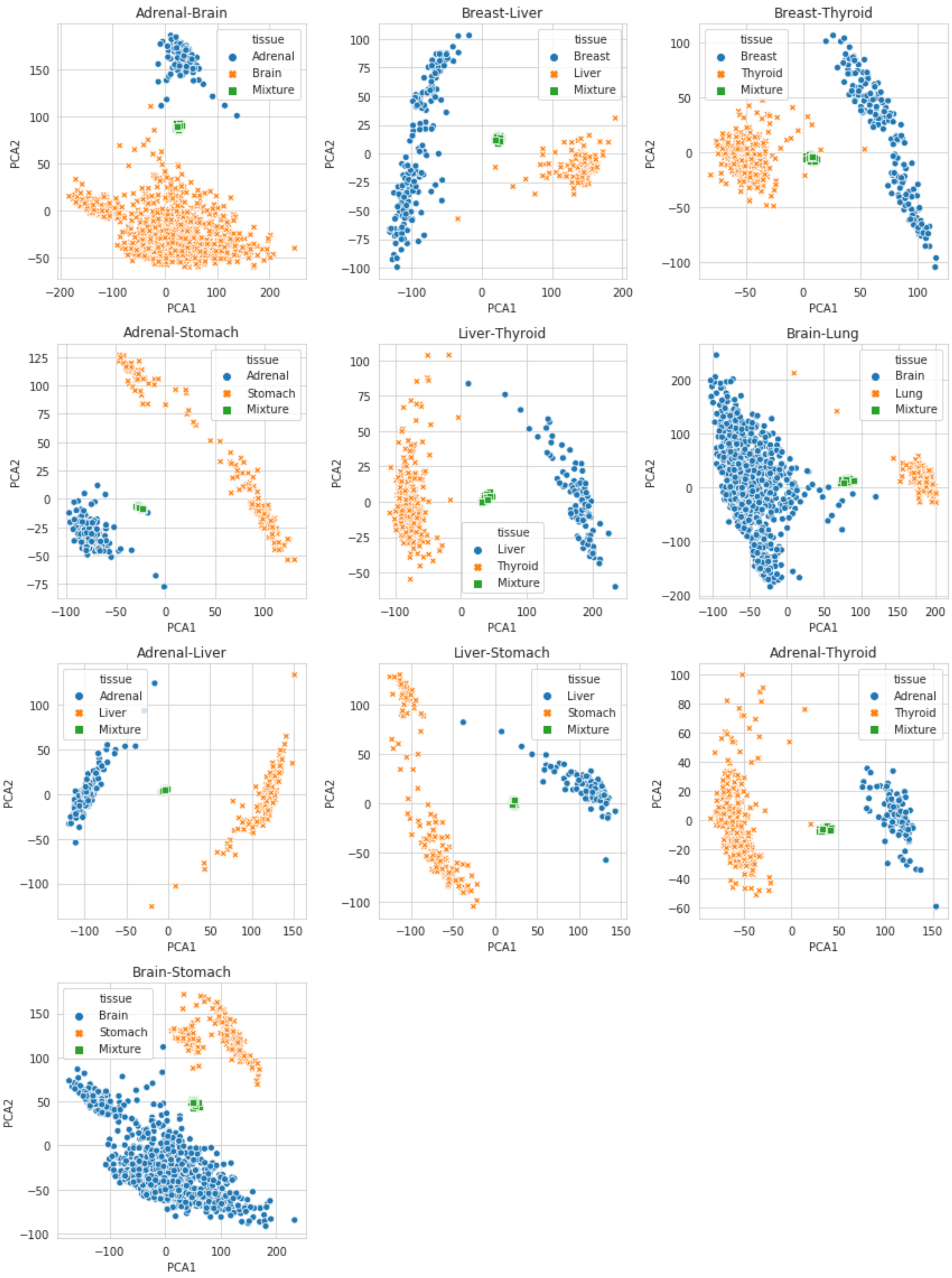

#### 3.4 Generate Manifest for Run

Create a manifest for the Toil version of the workflow

```
In [6]: out_dir = '/mnt/normsd-outlier-runs/mixture-simulation/'
        m.manifest(tissues, matrix_dir, out_dir)
```

##### 3.5 Run Samples

Command used to run mixture samples

```
#!/usr/bin/env bash
source activate toil
python /mnt/gene-outlier-detection/toil/toil-outlier-detection.py \
    --sample /mnt/data/outlier/tumor.hd5 \
    --background /mnt/data/outlier/gtex.hd5 \
    --manifest /mnt/normsd-outlier-runs/mixture-simulation/manifest.tsv \
    --out-dir /mnt/normsd-outlier-runs/mixture-simulation/output \
    --gene-list /mnt/data/outlier/drug-genes.txt \
    --group tissue \
    --col-skip 5 \
    --num-backgrounds 4 \
    --max-genes 125 \
    --workDir /mnt/ \
    /mnt/jobStore
```

##### 3.6 Model Weights for Contributing Tissues

After running the mixture samples through the model, we can plot the weights assigned to the two tissues used to generate the mixture samples. Ideally, the model will assign split weight between each of the two contributing tissues.

```
In [8]: sample_dir = '/mnt/normsd-outlier-runs/mixture-simulation/output/'
        wdf = m.weight_df(sample_dir)
        m.plot_weight_swarm(wdf);
        plt.tight_layout()
        plt.savefig(
            '/mnt/figures/Mixture-Simulation/average-weight.png',
            dpi=300,
            transparent=True
        )
```

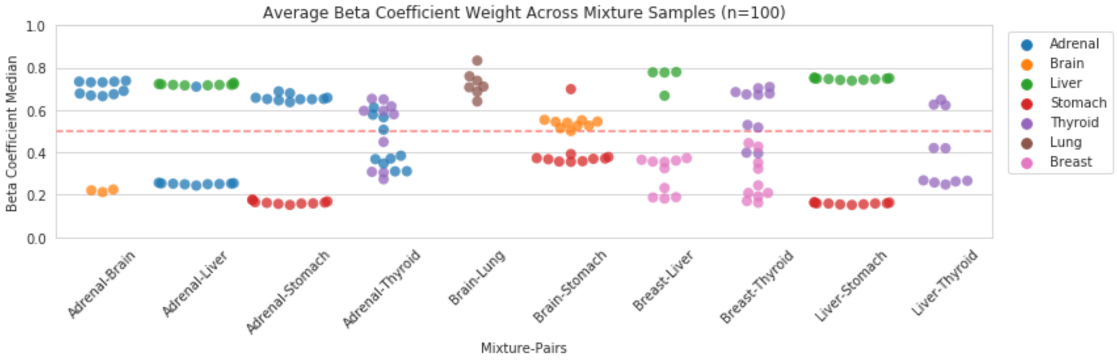

##### 3.7 Dimensionality Reductions of Mismatched Mixture Samples

Some of the mixture samples did not quite meet our expectation of being assigned a majority of the model weight. We can contextualize these results by plotting dimensionality reductions of those mixture samples alongside tissues that the model assigned weight to.

```
In [30]: sample_dir = '/mnt/normsd-outlier-runs/mixture-simulation/output/'
         matrix_dir = '/mnt/normsd-outlier-runs/mixture-simulation/matrices'
         wdf = Mixture.weight_df(sample_dir, only_contributing_tissues=False)
```

###### 3.7.1 Brain-Lung

Brain-Lung simulated mixture samples cluster more closely with prostate and breast tissue than brain or lung, which is reflected in the model's assignment of weight.

**PCA**

```
In [21]: tissues = ['Brain', 'Lung', 'Breast', 'Prostate']
         matrix_path = os.path.join(matrix_dir, 'Brain-Lung.hd5')
         m.plot_pca_nearby_tissues(matrix_path, tissues);
```

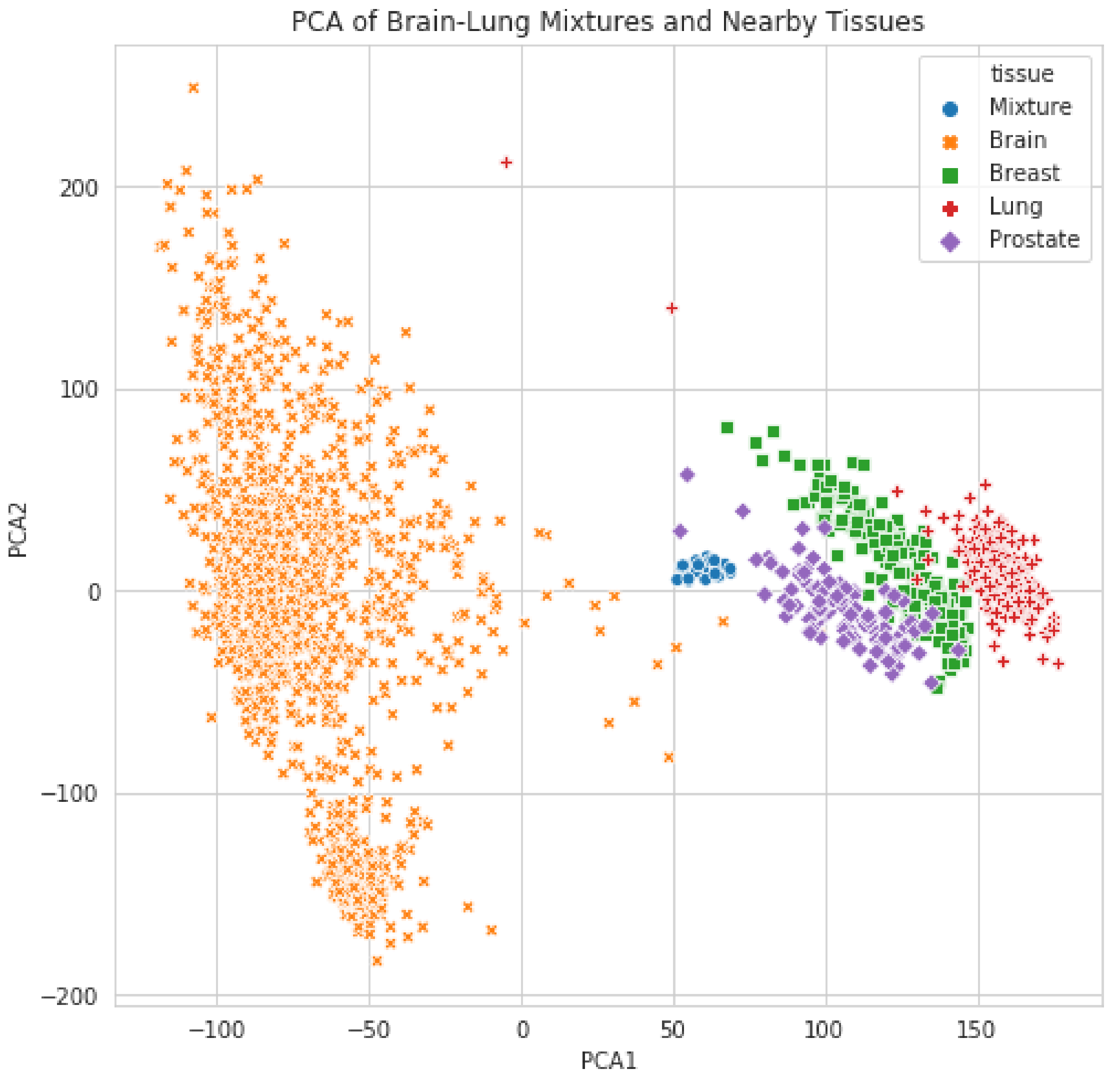

**Model Weight**

```
In [19]: m.plot_weight_boxplot(wdf.sort_values('tissue'), label='Brain-Lung');
```

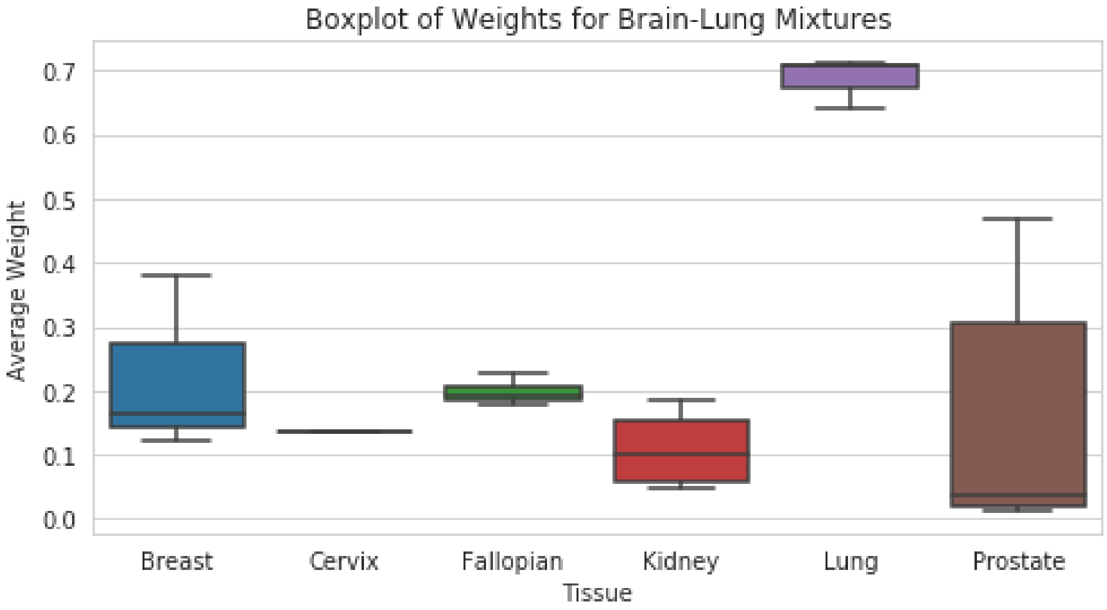

##### 3.7.2 Adrenal-Brain

Simulated Adrenal-Brain samples cluster near pituitary and kidney tissues. Adrenal tissue is also more homogenous than brain which might explain why the model prefers assign weight to adrenal samples over brain.

**PCA**

```
In [33]: tissues = ['Adrenal', 'Brain', 'Kidney', 'Pituitary']
         matrix_path = os.path.join(matrix_dir, 'Adrenal-Brain.hd5')
         m.plot_pca_nearby_tissues(matrix_path, tissues);
```

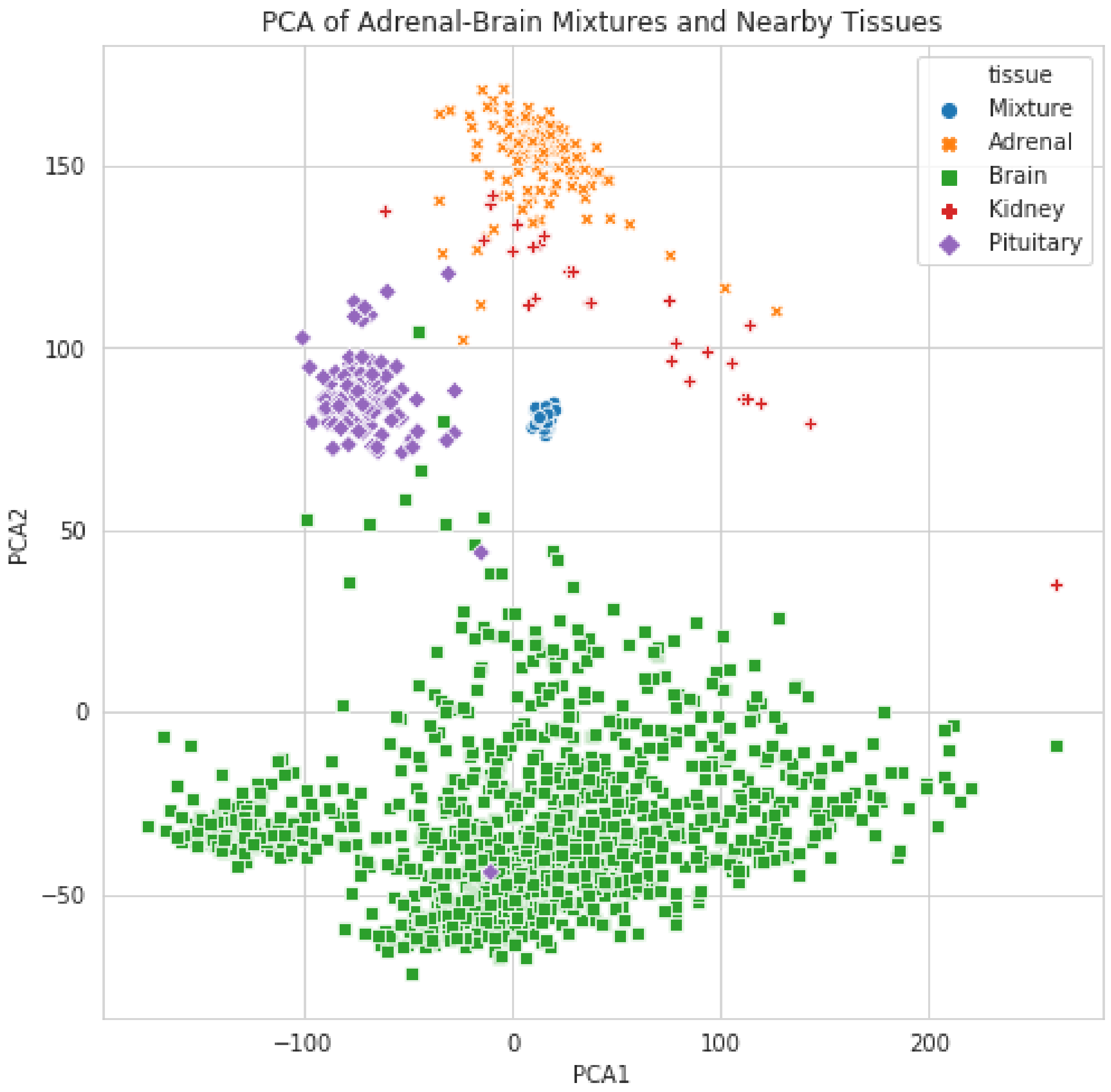

**Model Weight**

```
In [31]: m.plot_weight_boxplot(wdf.sort_values('tissue'), label='Adrenal-Brain');
```

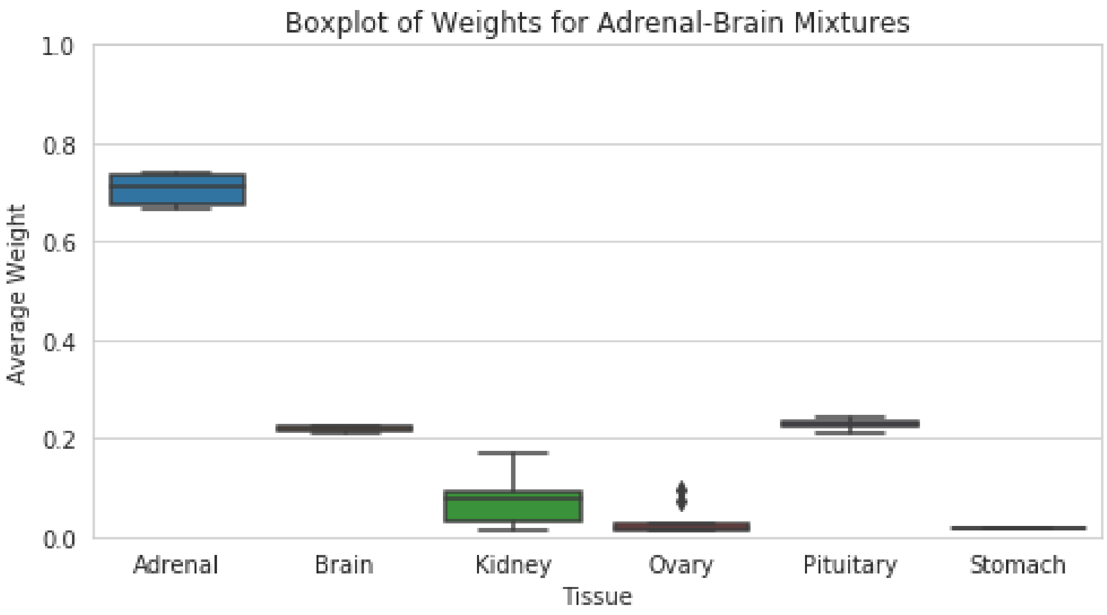

##### 3.7.3 Liver-Thyroid

Liver-Thyroid mixture samples don't get weight assigned to liver, but to thyroid, kidney, and minor salivary gland tissues. Although more proximal to liver than minor salivary gland across the first two principal components, minor samples are likely closer to the mixture samples along another component axis.

###### PCA

```
In [24]: tissues = ['Thyroid', 'Liver', 'Kidney', 'Minor']
         matrix_path = os.path.join(matrix_dir, 'Liver-Thyroid.hd5')
         m.plot_pca_nearby_tissues(matrix_path, tissues);
```

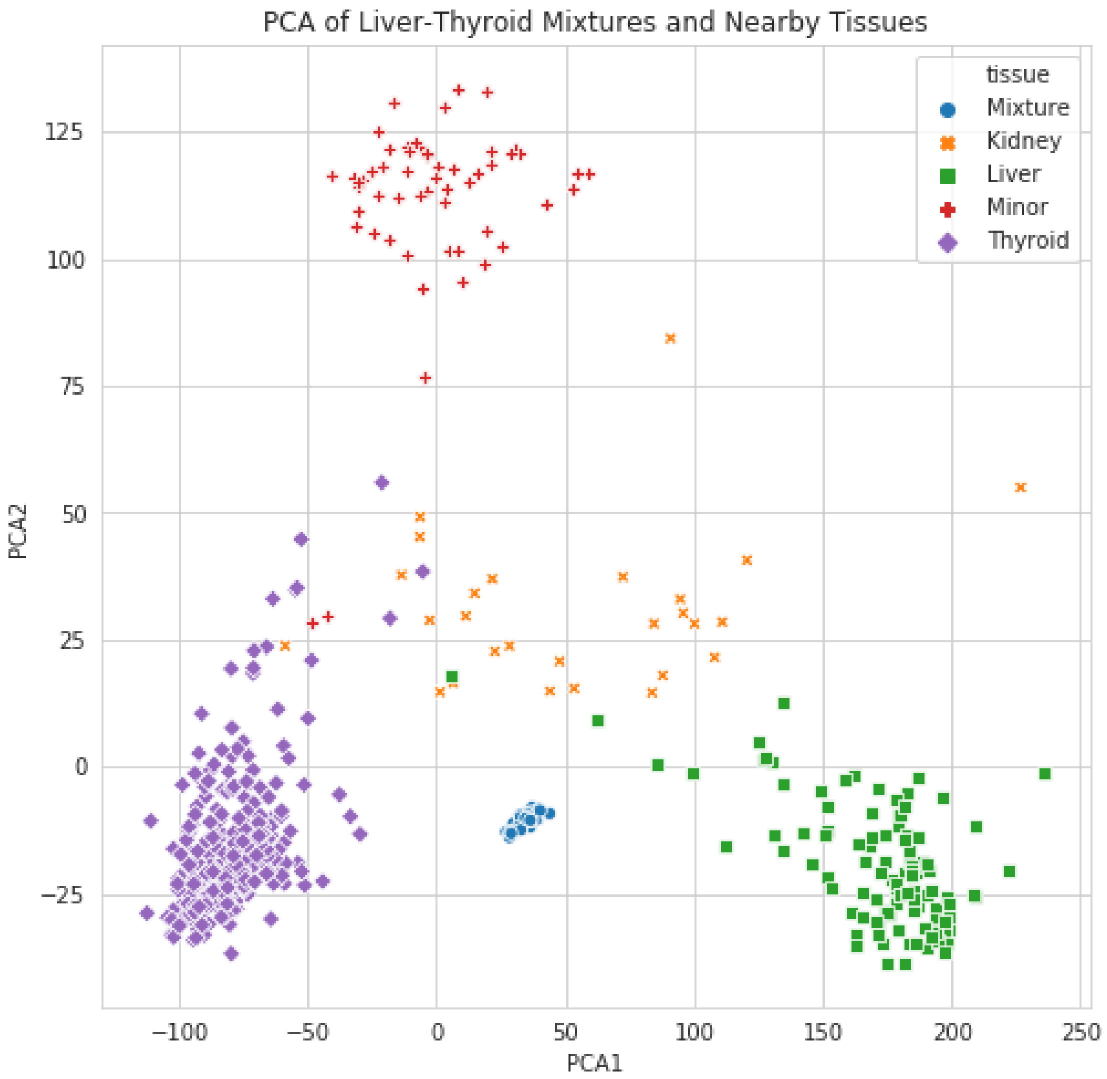

**Model Weight**

```
In [23]: m.plot_weight_boxplot(wdf.sort_values('tissue'), label='Liver-Thyroid');
```

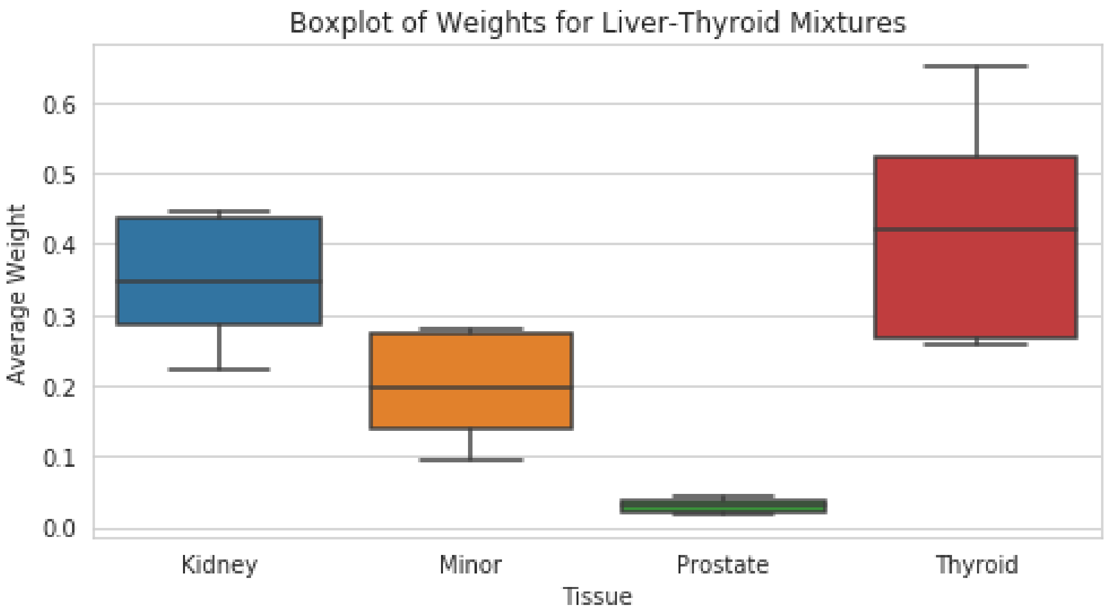

#### 4 Effect of Removing Matched Normal on Gene P-values

To test the robustness of the model, we explore the assocation between the model weight assigned to a particular normal tissue in the background for a sample and how removing that tissue effects the generated p-values.

All data used to generate the following section is avilable at http://courtyard.gi.ucsc.edu/~jvivian/outlier-paper/experiments/gtex-1000-match-removed.tar.gz. This data is compared directly to data in the **gtex-1000.tar.gz** archive in the same directory.

```
In [1]: import os
        import pandas as pd
        from scipy.stats import pearsonr

        class MatchRemove:
            def __init__(self, background_path):
                self.df = self._load_df(background_path)

            @staticmethod
            def _load_df(path):
                """Loads DataFrame"""
                print(f"Reading in {path}")
                if path.endswith(".csv"):
                    df = pd.read_csv(path, index_col=0)
                elif path.endswith(".tsv"):
                    df = pd.read_csv(path, sep="\t", index_col=0)
                else:
                    try:
                        df = pd.read_hdf(path)
```

```
            except Exception as e:
                print(e)
                raise RuntimeError(f"Failed to open DataFrame: {path}")
        return df

    def create_backgrounds(self, tissues, out_dir):
        for tissue in tissues:
            sub = self.df[self.df.tissue != tissue]
            out = os.path.join(out_dir, f"gtex-{tissue}-removed.hd5")
            sub.to_hdf(out, key="exp")

    def manifest(self, samples, sample_path, background_dir, out_dir):
        df = self._load_df(sample_path)
        tissues = df.tissue
        out_dir = os.path.join(out_dir, "manifest.tsv")
        with open(out_dir, "w") as f:
            f.write("sample\tbackground\n")
            for sample in samples:
                tissue = tissues.loc[sample]
                bg_path = os.path.join(background_dir, f"gtex-{tissue}-removed.hd5")
                f.write(f"{sample}\t{bg_path}\n")

    @staticmethod
    def weight_pearsonR_df(out_dir, mr_dir, sample_path):
        samples = os.listdir(mr_dir)
        df = MatchRemove._load_df(sample_path)
        tissues = df.tissue
        rows = []
        for sample in samples:
            tissue = tissues.loc[sample]
            # Collect median weight
            weight_path = os.path.join(out_dir, sample, "weights.tsv")
            weights = pd.read_csv(weight_path, sep="\t", index_col=0)
            try:
                w = weights.loc[tissue].Median
            except KeyError:
                w = 0
            # Collect pvals and calculate PearsonR
            norm_path = os.path.join(out_dir, sample, "pvals.tsv")
            norm_pvals = pd.read_csv(norm_path, sep="\t", index_col=0)
            mr_path = os.path.join(mr_dir, sample, "pvals.tsv")
            mr_pvals = pd.read_csv(mr_path, sep="\t", index_col=0)
            pvals = pd.concat([norm_pvals, mr_pvals], axis=1).dropna()
            pvals.columns = ["norm", "mr"]
            r, _ = pearsonr(pvals["norm"], pvals["mr"])
            rows.append([w, r, tissue, sample])
        columns = ["Weight", "PearsonR", "Tissue", "Sample"]
        return pd.DataFrame(rows, columns=columns)
```

```
In [2]: m = MatchRemove('/mnt/data/outlier/gtex.hd5')
```

```
Reading in /mnt/data/outlier/gtex.hd5
```

#### 4.1 Create Background Dataset for Every Tissue

This background dataset will have all samples belonging to a particular tissue removed

```
In [ ]: tissues = ["Adrenal", "Bladder", "Brain", "Breast", "Kidney",
                   "Liver", "Lung", "Prostate", "Stomach", "Thyroid"]
        background_dir = '/mnt/data/outlier/match-remove'
        m.create_backgrounds(tissues, background_dir)
```

#### 4.2 Create Manifest for Samples

Manifest will associate a sample with the appropriate background dataset that has its matched tissue in GTEx removed

```
In [ ]: samples = os.listdir('/mnt/normsd-outlier-runs/gtex-1000/')
        m.manifest(
            samples=samples,
            sample_path='/mnt/data/outlier/tumor.hd5',
            background_dir=background_dir,
            out_dir='/mnt/data/outlier/match-remove/'
        )
```

#### 4.3 Run Samples

```
#!/usr/bin/env bash
source activate toil
python /mnt/gene-outlier-detection/toil/toil-outlier-detection.py \
    --sample /mnt/data/outlier/tumor.hd5 \
    --background /mnt/data/outlier/gtex.hd5 \
    --gene-list /mnt/data/outlier/drug-genes.txt \
    --manifest /mnt/normsd-outlier-runs/variable-backgrounds/manifest.tsv \
    --out-dir /mnt/normsd-outlier-runs/variable-backgrounds/output/ \
    --group tissue \
    --col-skip 5 \
    --num-backgrounds 1 \
    --max-genes 105 \
    --workDir /mnt/ \
    --disable-iter \
    /mnt/jobStore
```

#### 4.4 Calculate Pearson Correlations

```
In [3]: out_dir = '/mnt/normsd-outlier-runs/gtex-1000/'
        mr_dir = '/mnt/normsd-outlier-runs/gtex-1000-match-removed/'
        sample_path = '/mnt/data/outlier/tumor.hd5'

        df = m.weight_pearsonR_df(
            out_dir=out_dir,
            mr_dir=mr_dir,
            sample_path=sample_path
        )
        df = df.sort_values('Tissue')
        df.head()
```

```
Reading in /mnt/data/outlier/tumor.hd5
```

```
Out[3]:        Weight  PearsonR   Tissue          Sample
        394  0.939996  0.699236  Adrenal  TCGA-OR-A5JR-01
        125  0.935275  0.776023  Adrenal  TCGA-OR-A5KW-01
        632  0.970991  0.611433  Adrenal  TCGA-PK-A5H9-01
        635  0.961370  0.783986  Adrenal  TCGA-PK-A5HB-01
        127  0.897582  0.723726  Adrenal  TCGA-OR-A5K8-01
```

#### 4.5 Plot Weight by PearsonR

```
In [4]: import seaborn as sns
        import matplotlib.pyplot as plt
        sns.set(font_scale=1.25)
        sns.set_style('whitegrid')
```

```
In [5]: plt.figure(figsize=(12, 4))
        filled_markers = ('o', 'v', '^', '<', '>', '8', 's', 'p', '*', 'h', 'H', 'D', 'd')
        sns.scatterplot(
            data=df,
            x='Weight',
            y='PearsonR',
            hue='Tissue',
            style='Tissue',
            markers=filled_markers,
            alpha=0.9,
            s=60
        )
        plt.title(f'PearsonR of P-values after Removing Matched Tissue (N={len(df)})')
        plt.xlabel('GTEx Matched Tissue Weight')
        plt.ylim([0, 1.05])
        plt.xlim([-0.025, 1.025])
        plt.legend(bbox_to_anchor=(1.03, 1));
        plt.tight_layout()
        plt.savefig(
            '/mnt/figures/match-remove/weight-pearson.png',
            dpi=300,
            transparent=True
        )
```

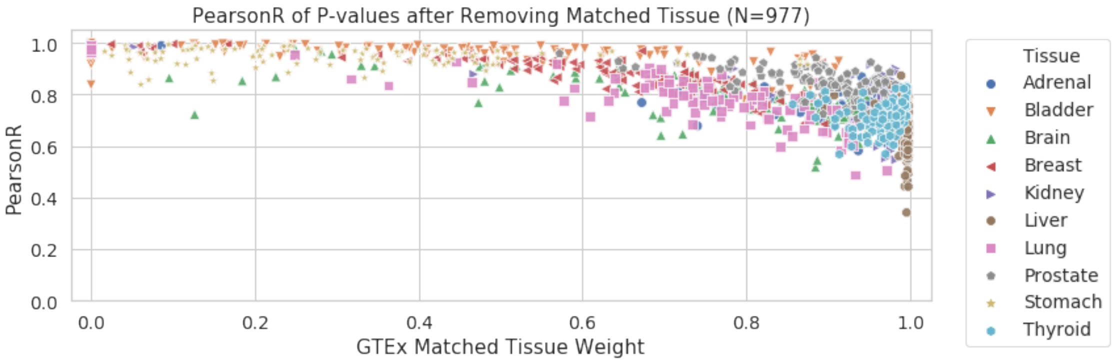

#### 5 Quantify Outliers Across Cancer Subtypes

Given an arbitrary p-value cutoff of 0.05, quantify up-regulated expression outliers across 10 tumor subtypes for ~1,000 TCGA samples when GTEx is used as the background comparison set. All data used to generate the following section is avilable at http://courtyard.gi.ucsc.edu/~jvivian/outlier-paper/experiments/gtex-1000.tar.gz.

In [1]:
```
import pandas as pd
import numpy as np
import os
import scipy.stats as st

import matplotlib.pyplot as plt
import seaborn as sns

sns.set(font_scale=1.20)
sns.set_style("whitegrid")

class Outliers:
    def __init__(self, sample_dir, sample_path):
        self.sample_dir = sample_dir
        self.sample_path = sample_path
        self.df = self._load_df(sample_path)
        self.tissue = self.df.tissue
        self.subtype = self.df.subtype
        self.pval_df = self._pval_df()

    def _pval_df(self) -> pd.DataFrame:
        pvals = []
        for sample_name in os.listdir(self.sample_dir):
            # Check if matched tissue was used in the weight
            w_path = os.path.join(self.sample_dir, sample_name, "weights.tsv")
            w = pd.read_csv(w_path, sep="\t", index_col=0)
            tissue = self.tissue.loc[sample_name]
            if not tissue in w.index:
                continue
            pval_path = os.path.join(self.sample_dir, sample_name, "pvals.tsv")
            p = pd.read_csv(pval_path, sep="\t")
            p["sample"] = sample_name
            p["tissue"] = self.tissue.loc[sample_name]
            p["subtype"] = self.subtype.loc[sample_name]
            pvals.append(p)
        pvals = pd.concat(pvals)
        return pvals.sort_values("tissue").reset_index(drop=True)

    @staticmethod
    def _load_df(path):
        """Loads DataFrame"""
        print(f"Reading in {path}")
        if path.endswith(".csv"):
            df = pd.read_csv(path, index_col=0)
        elif path.endswith(".tsv"):
            df = pd.read_csv(path, sep="\t", index_col=0)
```

```
            else:
                try:
                    df = pd.read_hdf(path)
                except Exception as e:
                    print(e)
                    raise RuntimeError(f"Failed to open DataFrame: {path}")
            return df
```

```
In [2]: drug_path = '/mnt/data/outlier/drug-genes.txt'
        drug_genes = sorted([x.strip() for x in open(drug_path).readlines()])
```

#### 5.1 Outlier Counts

```
In [3]: o = Outliers(
            sample_dir='/mnt/normsd-outlier-runs/gtex-1000/',
            sample_path='/mnt/data/outlier/tumor.hd5'
        )
```

```
Reading in /mnt/data/outlier/tumor.hd5
```

```
In [4]: pdf = o.pval_df
        pdf = pdf[pdf.Pval <= 0.05]
        pdf = pdf[pdf.Gene.isin(drug_genes)]
```

```
In [5]: counts = pdf.groupby('tissue')['Gene'].value_counts().reset_index(name='count')
```

```
In [6]: plt.figure(figsize=(12, 4))
        plt.hist(counts['count'], bins=20)
        plt.title('Gene outlier counts')
        plt.xlabel('Number of outliers')
```

```
Out[6]: Text(0.5, 0, 'Number of outliers')
```

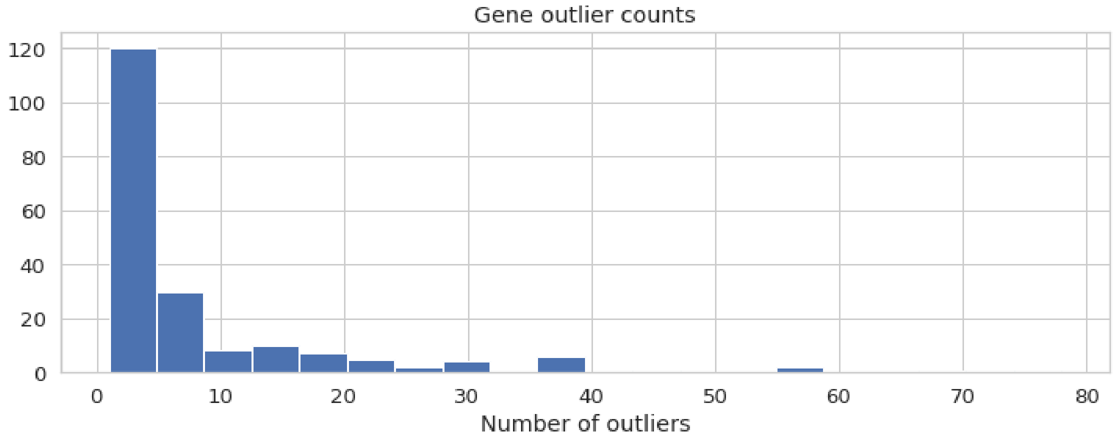

#### 5.2 Heatmap of Common Outlier Genes

```
In [7]: counts.columns = ['Tissue', 'Gene', 'Counts']
        counts.head()
```

```
Out[7]:      Tissue   Gene  Counts
        0   Adrenal    KIT      25
        1   Adrenal  CCNE1      23
        2   Adrenal    MET      17
        3   Adrenal  AURKA      12
        4   Adrenal  CCND1       6
```

```
In [8]: heatmap = counts.pivot(index='Gene', columns='Tissue', values='Counts')
```

First plot with genes with counts > 10

```
In [9]: heatmap = heatmap[heatmap.sum(axis=1) > 10]
```

```
In [10]: i = len(heatmap.index)//2
         genes1 = heatmap.index[:i]
         genes2 = heatmap.index[i:]
```

```
In [11]: f, ax = plt.subplots(1, 2, sharex=True, figsize=(12, 6))
         for i, genes in enumerate([genes1, genes2]):
             sub = heatmap[heatmap.index.isin(genes)]
             sns.heatmap(sub, ax=ax[i], cbar=i==1, cmap='Blues', vmin=0, vmax=79, annot=True)
         ax[1].set_ylabel(None)
         plt.tight_layout()
         out ='/mnt/figures/outlier-counts/outliers-talk.png'
         plt.savefig(out, dpi=300, transparent=True)
```

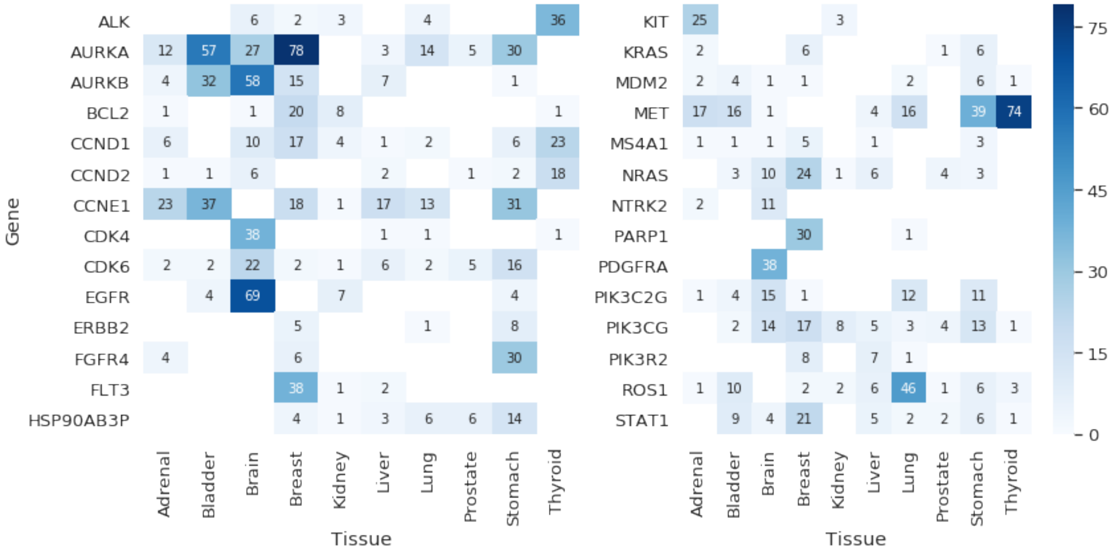

##### 5.2.1 Complete heatmap with all Treehouse genes

```
In [12]: heatmap = counts.pivot(index='Gene', columns='Tissue', values='Counts')
```

```
In [13]: i = len(heatmap.index)//2
         genes1 = heatmap.index[:i]
         genes2 = heatmap.index[i:]
```

```
In [14]: f, ax = plt.subplots(1, 2, sharex=True, figsize=(12, 8))
         for i, genes in enumerate([genes1, genes2]):
             sub = heatmap[heatmap.index.isin(genes)]
             sns.heatmap(sub, ax=ax[i], cbar=i==1, cmap='Blues', vmin=0, vmax=79, annot=True)
         ax[1].set_ylabel(None)
         plt.suptitle('Outliers (p < 0.05) in Treehouse Gene Set (N=977)')
         plt.tight_layout(rect=[0, 0.03, 1, 0.95])
         out ='/mnt/figures/outlier-counts/outliers-talk-all.png'
         plt.savefig(out, dpi=300, transparent=True)
```

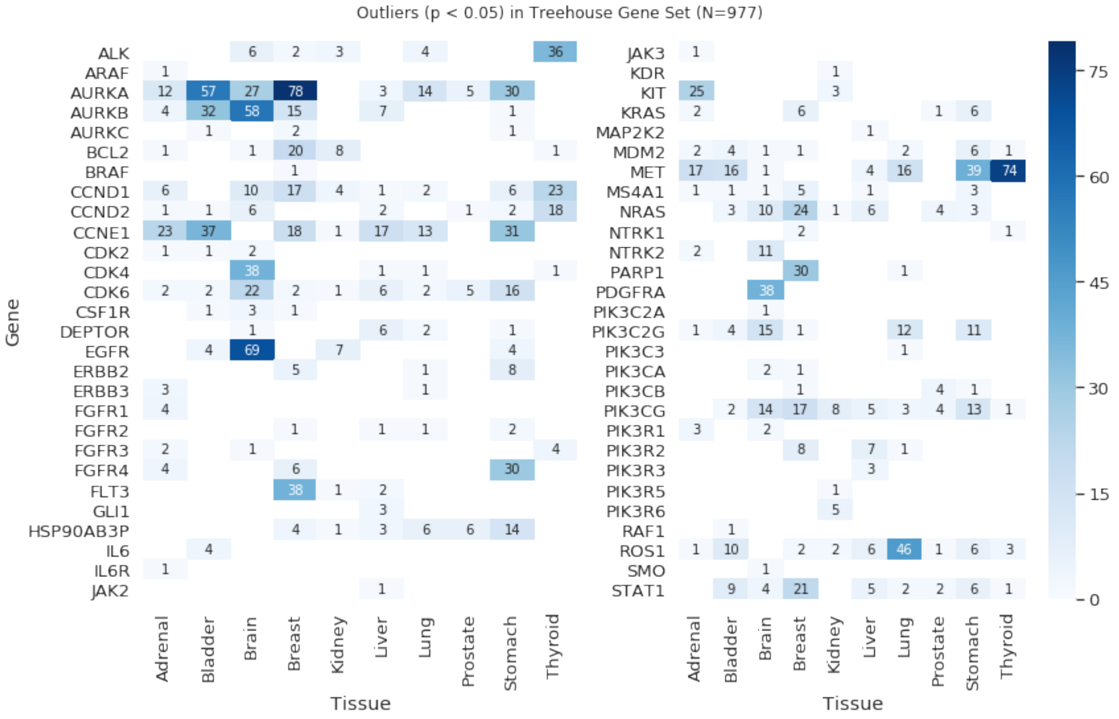

#### 5.3 Literature Corroboration of Outlier Findings

After quantifying outliers across the 10 different tumor subyptes, we assessed whether the high-count outliers we found were corroborated by literature as previously identified tumor markers for these cancer subtypes (Table 2). We found evidence for amplification of Aurora Kinase A (AURKA), a gene involved in microtubule formation, in bladder cancers, which has been used as a biomarker for early detection [1, 2]. AURKA is also frequently amplified in certain breast cancers [3]. Aurora Kinase B (AURKB) is often amplified in gliomas relative to normal brain tissue and is associated with histopathological grades [4]. Epidermal Growth Factor Receptor (EGFR) amplification is also commonly seen in both lower grade gliomas as well as high grade gliomas [5, 6, 7]. Many inhibitor drugs of EGFR also target Platelet Derived Growth Factor Receptor Alpha (PDGFRA), which is another gene often overexpressed in gliomas [8]. Both thyroid carcinomas and gastric cancers report high expression of the MET proto-oncogene [9, 10, 11, 12]. Finally, despite being more commonly known for forming fusion genes, ROS1 has also been identified as overexpressed in lung adenocarcinomas [13]. While RNA-seq quantification will still map reads from a fusion gene to the original gene, the high outlier count for ROS1 is likely due to weight assignment of lung tumors to minor salivary gland in GTEx which has no ROS1 expression. Lung adenocarcinomas with $< 15\%$ of model weight assigned to minor salivary gland reduced the percentage of lung samples that counted ROS1 as an outlier from 50% to 37%, which is more consistent with the literature values. Due to minor salivary gland samples clustering closely to several phenotypically distinct tissues (bladder, breast, lung, stomach), it would likely improve the overall analysis if removed from the GTEx cohort. Minor salivary gland was not removed for our analysis as it was discovered only after the performing the experiments and without a valid mathematical justification for removing it, would constitute an unprincipled decision.

We also see very little overexpression of HER2 (ERBB2) in breast cancers which is a relatively well-known overexpressed gene in a subset of breast cancers. This is due to the extreme gene expression variance seen in the normal tissue of breast in GTEx, which ranges from log2(0) to log2(10) transcripts per million. This means that HER2 in breast tumors would have to express HER2 $>$ log2(12) or log2(13) in order to count as an outlier. Based on the relatively large variance and bimodal features of many gene expression distributions in GTEx breast, it is possible that the GTEx breast dataset is comprised of several phenotypically distinct parts of the normal breast tissue. If GTEx breast tissues could be divided into sub-populations based on Clustering, it could potentially reduce the variance of many of these genes and improve the ability to detect outliers in breast tumor samples. We chose to use the unadulterated GTEx dataset for our analysis to keep things simple and easily reproducible.

| **Gene** | **Cancer** | **Paper** | **Author** | **Notes / Quotes** |
|---|---|---|---|---|
| AURKA | Bladder Urothelial Carcinoma | POTENTIAL NEW MARKERS IN THE EARLY DETECTION OF BLADDER CANCER | Samir P. Shirodkar and Vinata B. Lokeshwar | In a case control study, involving bladder cancer, normal individuals and patients with benign conditions, they reported 96.6% specificity and 87% sensitivity for AURKA-FISH test to detect bladder cancer. |
| AURKA | Bladder Urothelial Carcinoma | Quantitation of Aurora Kinase A Gene Copy Number in Urine Sediments and Bladder Cancer Detection | HS Park et al. | Forced overexpression of AURKA in urothelial cells induced amplification of centrosomes, chromosome missegregation, and aneuploidy, and natural overexpression was detectable in in situ lesions from patients with bladder cancer |
| AURKA | Bladder Urothelial Carcinoma | Quantitation of Aurora Kinase A Gene Copy Number in Urine Sediments and Bladder Cancer Detection | HS Park et al. | The Aurora kinase A (AURKA) gene, which encodes a key regulator of mitosis, is frequently amplified and/or overexpressed in cancer cells |
| AURKA | Breast Cancer | Aurora-A gene is frequently amplified in basal-like breast cancer | S Staff et al. | AURKA amplification (found in 21%) showed an association with basal-like tumor phenotype (p=0.046) |
| AURKA | Stomach Cancer | AURKA regulates JAK2eSTAT3 activity in human gastric and esophageal cancers | A Katsha et al. | Aurora kinase A is a frequently amplified and overexpressed gene in upper gastrointestinal adenocarcinomas (UGCs). |
| AURKB | Brain Cancer | Kinesin family member 2C (KIF2C/MCAK) is a novel marker for prognosis in human gliomas | L Bie et al. | We found that KIF2C and AURKB genes were higher expression in glioma samples when were compared with normal brain tissues. KIF2C and AURKB expression were associated with histopathological grades. |

| | | | | |
|---|---|---|---|---|
| CCND1 | Thyroid Carcinoma | Tissue array for Tp53, C-myc, CCND1 gene over-expression in different tumors | GY Liu et al. | In this study, a significant difference was found in carcinomatous and paracancerous tissue samples, including those of stomach, rectum, thyroid gland, liver, and mammary gland. Our data indicate that CCND1 expression was significantly associated with carcinomatous change. |
| CCNE1 | Bladder Urothelial Carcinoma | A multi-stage genome-wide association study of bladder cancer identifies multiple susceptibility loci | N Rothman et al. | Cyclin E1 expression in bladder cancer has been associated with high-grade or muscle-invasive tumors and poor clinical outcome |
| CCNE1 | Bladder Urothelial Carcinoma | High-Throughput Tissue Microarray Analysis of Cyclin E Gene Amplification and Overexpression in Urinary Bladder Cancer | J Richter et al. | Overexpression of cyclin E by mechanisms other than amplification seems common and is characteristic to a subset of bladder carcinomas, especially at the stage of early invasion. |
| CCNE1 | Bladder Urothelial Carcinoma | Tumour-suppressive microRNA-144-5p directly targets CCNE1/2 as potential prognostic markers in bladder cancer | R Matsushita et al. | The patients with high CCNE1 or CCNE2 expression had lower overall survival probabilities than those with low expression (P=0.025 and P=0.032). |
| CDK4 | Brain Cancer | Combinations of genetic mutations in the adult neural stem cell compartment determine brain tumour phenotypes | Thomas S Jacques et al. | The $5'$ transcript of Cdk4 is significantly (P<0.001) upregulated in brain tumours, whereas there is a statistically non[U+2010]significant upregulation of p16/Ink4a and p19/Arf transcripts. |
| EGFR | Brain Cancer | Genetic Pathways to Primary and Secondary Glioblastoma | H Ohgaki and P Kleihues | They affect mainly the elderly and are genetically characterized by loss of heterozygosity 10q (70% of cases), EGFR amplification (36%), p16INK4a deletion (31%), and PTEN mutations (25%) |
| EGFR | Brain Cancer | Clinical and Molecular Characteristics of Malignant Transformation of Low-Grade Glioma in Children | A Broniscer et al. | Although EGFR and ERBB4 overexpression were seen in both LGGs and HGGs, they occurred more commonly in the latter group. |
| EGFR | Brain Cancer | Gene expression profiling of metastatic brain cancer | VM Zohrabian et al. | Studies have examined the differences in EGFR expression between primary and secondary glioblastoma multiforme (GBM), and have found that EGFR overexpression is associated with tumor growth and angiogenesis |

| | | | | |
|---|---|---|---|---|
| MET | Thyroid Carcinoma | Overexpression of the c-MET/HGF receptor gene in human thyroid carcinomas | MFR Di et al. | The receptor is barely detectable, however, in normal thyroids and in specimens from patients affected by non-neoplastic thyroid diseases. Now we report that the expression of the Met/HGF receptor is increased a hundred fold in 22 out of 41 human carcinomas derived from the thyroid follicular epithelium |
| MET | Stomach Cancer | MET Expression and Amplification in Patients with Localized Gastric Cancer | YY Janjigian et al. | Although high levels of MET protein and mRNA were commonly encountered (in 63% and 50% of resected tumor specimens, respectively), none of these tumors had MET gene amplification by FISH, and only 6.6% had evidence of MET tyrosine kinase activity by p-MET IHC. |
| MET | Stomach Cancer | Durable Complete Response of Metastatic Gastric Cancer with Anti-Met Therapy Followed by Resistance at Recurrence | DVT Catenacci et al. | The primary tumor had high MET gene polysomy and evidence for an autocrine production of hepatocyte growth factor, the growth factor ligand of Met. |
| MET | Stomach Cancer | Impact of MET amplification on gastric cancer: Possible roles as a novel prognostic marker and a potential therapeutic target | J Lee et al. | Although the definitive role of MET oncogene is yet to be determined in carcinogenesis of gastric cancer, overexpression and amplification of c-Met has been demonstrated in gastric cancer cell lines |
| PDGFRA | Brain Cancer | Amplification of KIT, PDGFRA, VEGFR2, and EGFR in Gliomas | Marjut Puputti et al. | In conclusion, besides glioblastoma, amplified KIT, PDGFRA, and VEGFR may also occur in lower-grade gliomas and in their recurrent tumors. |
| ROS1 | Lung Cancer | Targeting ROS1 with Anaplastic Lymphoma Kinase Inhibitors: A Promising Therapeutic Strategy for a Newly Defined Molecular Subset of Non–Small-Cell Lung Cancer | Leow PayChin et al. | Microarray analysis of NSCLC demonstrated significantly elevated ROS1 expressions in 20% to 30% of cases, and elevated ROS1 expression was found to be part of a molecular signature for lung adenocarcinoma subtype |

Table 2: References that corroborate outlier findings for all genes identified as outliers in over half of samples within a tumor subtype.

### 6 Negative Control Experiment

This section shows the p-value distributions of 100 GTex samples when different background datasets are used in the model. Comparing normal samples to other normals should result in a posterior predictive p-value distribution that is peaked around 0.5 (centered). Ergo, there should be far fewer outliers when GTEx and TCGA-normal are the backgrounds compared to TCGA-tumor. We expect the p-value distribution when tumor is the background to have longer tails that extend below 0.05 and above 0.95.

All data used to generate the following section is avilable at http://courtyard.gi.ucsc.edu/~jvivian/outlier-paper/experiments/negative-control.tar.gz.

```
In [16]: import pandas as pd
         import numpy as np
         import os
```

#### 6.1 Randomly select samples

```
In [14]: tissues = ["Adrenal", "Bladder", "Brain", "Breast", "Kidney",
                    "Liver", "Lung", "Prostate", "Stomach", "Thyroid"]
         gtex = pd.read_hdf('/mnt/data/outlier/gtex.hd5')
```

```
In [19]: samples = {}
         for tissue in tissues:
             gtex_samples = list(gtex[gtex.tissue == tissue].index)
             samples[tissue] = list(np.random.choice(gtex_samples, 10))
```

#### 6.2 Create manifest

```
In [22]: manifest_out = '/mnt/normsd-outlier-runs/negative-control/manifest.tsv'
         output_path = '/mnt/out'
         with open(manifest_out, 'w') as f:
             f.write('sample\tbackground\tout_dir\n')
             for k, v in samples.items():
                 for background in ['gtex', 'tumor', 'normal']:
                     background_path = f'/mnt/inputs/{background}.hd5'
                     experiment_dir = 'mnt/outputs/norm-sd-runs/negative-control'
                     out_path = os.path.join(experiment_dir, background)
                     for sample in v:
                         f.write(f'{sample}\t{background_path}\t{out_path}\n')
```

#### 6.3 Run Samples

```
#!/usr/bin/env bash
source activate toil
python toil-outlier-detection.py \
    --sample /mnt/inputs/gtex.hd5 \
    --background /mnt/inputs/gtex.hd5 \
    --manifest /mnt/inputs/norm-sd-runs/negative-control/manifest.tsv \
    --out-dir /mnt/outputs/norm-sd-runs/negative-control/ \
    --gene-list /mnt/inputs/drug-genes.txt \
    --group tissue \
    --col-skip 5 \
    --num-backgrounds 4 \
    --max-genes 125 \
    --workDir /mnt/ \
```

```
    --disable-iter \
    /mnt/jobStore
```

#### 6.4 Analysis

```
In [94]: class Outliers:
             def __init__(self, sample_dir, sample_path):
                 self.sample_dir = sample_dir
                 self.sample_path = sample_path
                 self.df = self._load_df(sample_path)
                 self.tissue = self.df.tissue
                 self.pval_df = self._pval_df()

             def _pval_df(self) -> pd.DataFrame:
                 pvals = []
                 for dataset in os.listdir(self.sample_dir):
                     if dataset == 'manifest.tsv':
                         continue
                     for sample_name in os.listdir(os.path.join(self.sample_dir, dataset)):
                         sample_path = os.path.join(self.sample_dir, dataset, sample_name)
                         pval_path = os.path.join(sample_path, "pvals.tsv")
                         p = pd.read_csv(pval_path, sep="\t")
                         p["sample"] = sample_name
                         p["tissue"] = self.tissue.loc[sample_name]
                         p["dataset"] = dataset.capitalize()
                         pvals.append(p)
                 pvals = pd.concat(pvals)
                 return pvals.sort_values("tissue").reset_index(drop=True)

             @staticmethod
             def _load_df(path):
                 """Loads DataFrame"""
                 print(f"Reading in {path}")
                 if path.endswith(".csv"):
                     df = pd.read_csv(path, index_col=0)
                 elif path.endswith(".tsv"):
                     df = pd.read_csv(path, sep="\t", index_col=0)
                 else:
                     try:
                         df = pd.read_hdf(path)
                     except Exception as e:
                         print(e)
                         raise RuntimeError(f"Failed to open DataFrame: {path}")
                 return df
```

```
In [95]: o = Outliers(
             sample_dir='/mnt/normsd-outlier-runs/negative-control/',
             sample_path='/mnt/data/outlier/gtex.hd5'
         )
```

```
Reading in /mnt/data/outlier/gtex.hd5
```

```
In [96]: drug_path = '/mnt/data/outlier/drug-genes.txt'
         drug_genes = sorted([x.strip() for x in open(drug_path).readlines()])
```

In [81]:
```python
import matplotlib.pyplot as plt
import seaborn as sns
sns.set(font_scale=1.25)
sns.set_style('whitegrid')
```

In [109]:
```python
pdf = o.pval_df
pdf = pdf[pdf.Gene.isin(drug_genes)]
pdf = pdf.sort_values(['tissue', 'dataset'])
pdf.head()
```

Out[109]:

| | Gene | Pval | sample | tissue | dataset |
|---|---|---|---|---|---|
| 56 | MAP2K1 | 0.359 | GTEX-R53T-0226-SM-48FEH | Adrenal | Gtex |
| 57 | MTOR | 0.359 | GTEX-R53T-0226-SM-48FEH | Adrenal | Gtex |
| 58 | HSP90AB1 | 0.363 | GTEX-R53T-0226-SM-48FEH | Adrenal | Gtex |
| 59 | FGFR4 | 0.370 | GTEX-R53T-0226-SM-48FEH | Adrenal | Gtex |
| 60 | PIK3CD | 0.377 | GTEX-R53T-0226-SM-48FEH | Adrenal | Gtex |

In [110]:
```python
plt.figure(figsize=(12, 4))
sns.kdeplot(pdf[pdf.dataset == 'Gtex'].Pval, label='GTEx', alpha=0.5, shade=True)
sns.kdeplot(pdf[pdf.dataset == 'Normal'].Pval, label='Normal', alpha=0.5, shade=True)
sns.kdeplot(pdf[pdf.dataset == 'Tumor'].Pval, label='Tumor', alpha=0.5, shade=True)
plt.title('P-value Disribution of GTEx samples (N=100) \
        compared to GTEx, TCGA-Normal, and TCGA-Tumor')
plt.xlabel('P-value')
plt.ylabel('Density')
plt.xlim([-0.05, 1.05])
plt.tight_layout()
plt.savefig(
    '/mnt/figures/negative-control/pval-dist.png',
    dpi=300,
    transparent=True
)
```

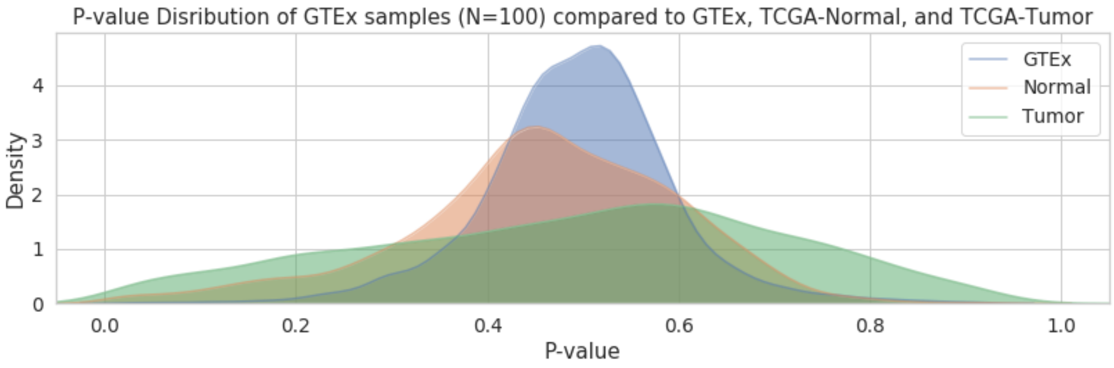

In [119]:
```python
plt.figure(figsize=(12, 4))
plt.axhline(0.5, c='r', ls='--', lw=3)
sns.boxplot(data=pdf, x='tissue', y='Pval', hue='dataset')
plt.legend(bbox_to_anchor=[1, 1])
plt.title('P-value Distributions for GTEx Samples by \
        Background Dataset and Tissue (N=100)');
plt.tight_layout()
```

```
plt.savefig(
    '/mnt/figures/negative-control/pval-dist-boxplot.png',
    dpi=300,
    transparent=True
)
```

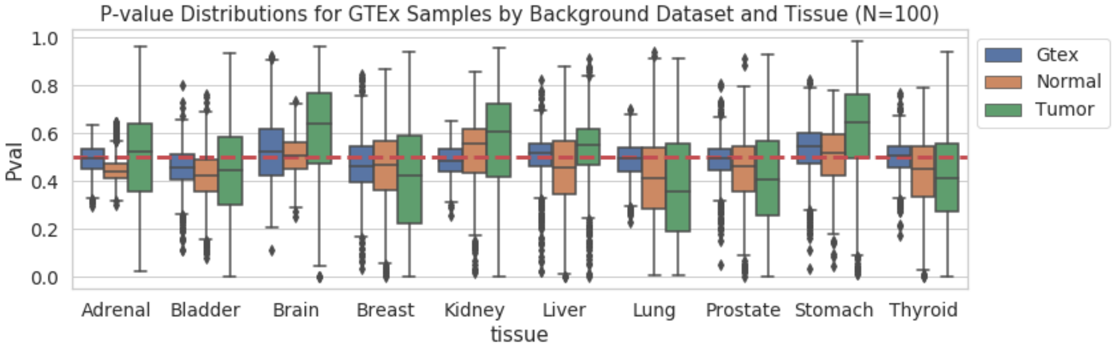

#### 7 Background Dataset Selection

Due to the choice of Dirichlet distribution for the $\beta$ coefficients, which results in most background tissues being assigned $\sim 0$ weight, it is reasonable to select a subset of background tissues to keep runtime low. Instead of choosing an arbitrary number of background datasets, the model can be iteratively run with more and more background datasets until the p-values converge. Background datasets are first ranked by selecting 10% of the genes via ANOVA F-value and then calculating the pairwise distance between the N-of-1 sample and the background datasets. The model is then run, starting with the most proximal dataset, and datasets are iteratively added until p-values converge to a Pearson correlation greater than 0.99. This can drastically reduce runtime for N-of-1 samples with a strong match to one of the background datasets and is flexible for samples that are more appropriately modeled by multiple background datasets. The complete method for ranking background sets is shown below.

Python implementation for ranking background datasets

```
def anova_distances(
    sample: pd.Series,
    df: pd.DataFrame,
    genes: List[str],
    group: str = "tissue",
    percent_genes=0.10,
):
    """
    Calculates distance to each group via pairwise distance using top N ANOVA genes

    Args:
        sample: n-of-1 sample. Gets own label
        df: background dataset
        genes: genes to use for pairwise distance
        group: Column to use as class discriminator
        percent_genes: Percent of ANOVA genes to use for pairwise distance

    Returns:
        DataFrame of pairwise distances
    """
    click.echo(f"Ranking background datasets by {group} via ANOVA")
    n_genes = int(percent_genes * len(genes))
    skb_genes = select_k_best_genes(df, genes, n=n_genes)
    dist = pairwise_distances(np.array(sample[skb_genes]).reshape(1, -1), df[skb_genes])
```

```
    dist = pd.DataFrame([dist.ravel(), df["tissue"]]).T
    dist.columns = ["Distance", "Group"]

    # Median by group and sort
    med_dist = (
        dist.groupby("Group").apply(lambda x: x["Distance"].median()).reset_index()
    )
    med_dist.columns = ["Group", "MedianDistance"]
    return med_dist.sort_values("MedianDistance").reset_index(drop=True)
```

### 8 Number of Genes Effect on Model Output

This section explores the effect of varying the number of training genes on different aspects of the model output.

Additional training genes help the model identify background datasets most similar to the N-of-1 sample so the model can appropriately assign coefficient weights and produce an appropriate posterior distribution. Choosing a subset of genes that accurately discriminate between the different background datasets allows the model appropriately assign a majority of the weight to the "correct" background dataset that is most similar to the N-of-1. A fast method for selecting genes that fits this definition is to calculate the ANOVA F-value for each background dataset using all available genes and selecting those with the highest F-values.

$$F = \frac{\text{variation between sample means}}{\text{variation within samples}}$$

ANOVA was compared with Chi2, which measures dependence between variables, and was found to produce model weights that were more consistent with the appropriate matched tissue. All data used to generate the following section is avilable at http://courtyard.gi.ucsc.edu/~jvivian/outlier-paper/experiments/variable-genes.tar.gz.

```
In [1]: import os
        import pickle

        import matplotlib.pyplot as plt
        import pandas as pd
        import seaborn as sns
        import numpy as np
        from scipy.stats import pearsonr

        sns.set_style("whitegrid")

        class Genes:
            def __init__(
                    self, sample_dir, background_path, sample_path, sample_name, genes_path
            ):
                self.sample_dir = sample_dir
                self.background_path = background_path
                self.sample_path = sample_path
                self.sample_name = sample_name
                self.bg = None
                self.sg = None
                self.df = None
                self.classes = None
                self.w = self._weight_df()
                self.p = self._pval_df()
```

```python
        self.traces = {}
        self.dfs = {}
        self.genes = [
            x.strip() for x in open(genes_path, "r").readlines() if not x.isspace()
        ]

    def _weight_df(self) -> pd.DataFrame:
        weights = []
        for subdir in os.listdir(self.sample_dir):
            max_genes = int(subdir)
            sample_name = os.listdir(os.path.join(self.sample_dir, subdir))[0]
            weight_path = os.path.join(
                self.sample_dir, subdir, sample_name, "weights.tsv"
            )
            w = pd.read_csv(weight_path, sep="\t")
            w.columns = ["tissue", "Median", "std"]
            w["sample"] = sample_name
            w["max_genes"] = int(max_genes)
            w["num_training_genes"] = int(max_genes) - 85
            weights.append(w.drop("std", axis=1))
        weights = pd.concat(weights).reset_index(drop=True)
        self.classes = sorted(weights.tissue)
        return weights.sort_values("max_genes")

    def _pval_df(self) -> pd.DataFrame:
        pvals = []
        for subdir in os.listdir(self.sample_dir):
            max_genes = int(subdir)
            sample_name = os.listdir(os.path.join(self.sample_dir, subdir))[0]
            pval_path = os.path.join(
                self.sample_dir,
                subdir,
                sample_name,
                "pvals.tsv"
            )
            p = pd.read_csv(pval_path, sep="\t")
            p["sample"] = sample_name
            p["max_genes"] = int(max_genes)
            p["num_training_genes"] = int(max_genes) - 85
            pvals.append(p)
        pvals = pd.concat(pvals).reset_index(drop=True)
        return pvals.sort_values("max_genes")

    def _load_df(self, path):
        if path in self.dfs:
            return self.dfs[path]
        print(f"Reading in {path}")
        if path.endswith(".csv"):
            df = pd.read_csv(path, index_col=0)
        elif path.endswith(".tsv"):
            df = pd.read_csv(path, sep="\t", index_col=0)
        else:
            try:
                df = pd.read_hdf(path)
```

```python
        except Exception as e:
            print(e)
            raise RuntimeError(f"Failed to open DataFrame: {path}")
    self.dfs[path] = df
    return df

@staticmethod
def _load_model(pkl_path):
    with open(pkl_path, "rb") as buff:
        data = pickle.load(buff)
    return data["model"], data["trace"]

def _df(self, gene, tissue) -> pd.DataFrame:
    p = []
    for i, subdir in enumerate(os.listdir(self.sample_dir)):
        max_genes = int(subdir)
        sample_name = os.listdir(os.path.join(self.sample_dir, subdir))[0]
        if subdir in self.traces:
            t = self.traces[subdir]
        else:
            if i == 0:
                print("Loading traces to extract posterior distribution")
            model_path = os.path.join(
                self.sample_dir, subdir, sample_name, "model.pkl"
            )
            m, t = self._load_model(model_path)
            self.traces[subdir] = t
        # Calculate PPC
        ppc = self._posterior_predictive_check(t, [gene])
        df = pd.DataFrame()
        df["ppc"] = ppc[gene]
        df["x"] = t[f"{gene}={tissue}"]
        df["max_genes"] = max_genes
        df["num_training_genes"] = int(max_genes) - 85
        p.append(df)
    p = pd.concat(p).reset_index(drop=True)
    return p.sort_values("max_genes")

def t_fits(self, backgrounds, group='tissue'):
    """
    StudentT distribution fits for every dataset/gene pair

    Args:
        df: Background dataframe to use in comparison
        genes: Genes to fit
        backgrounds: Background datasets to fit
        group: Column in background dataset to use as labels

    Returns:
        StudentT fits for every gene-dataset pair
    """
    df = self._load_df(self.background_path)
    genes = self.genes
    fits = {}
```

```python
        for gene in genes:
            for i, dataset in enumerate(backgrounds):
                # "intuitive" prior parameters
                prior_mean = 0.0
                prior_std_dev = 1.0
                pseudocounts = 1.0

                # convert to prior params of normal-inverse gamma
                kappa_0 = pseudocounts
                mu_0 = prior_mean
                alpha_0 = 0.5 * pseudocounts
                beta_0 = 0.5 / prior_std_dev ** 2

                # collect summary statistics for data
                observations = np.array(df[df[group] == dataset][gene])
                n = len(observations)
                obs_sum = np.sum(observations)
                obs_mean = obs_sum / n
                obs_ssd = np.sum(np.square(observations - obs_mean))

                # compute the posterior params
                kappa_n = kappa_0 + n
                mu_n = (kappa_0 * mu_0 + obs_sum) / (kappa_0 + n)
                alpha_n = alpha_0 + 0.5 * n
                beta_n = beta_0 + 0.5 * (
                        obs_ssd + kappa_0 * n * (obs_mean - mu_0) ** 2 / (kappa_0 + n)
                )

                # from https://www.seas.harvard.edu/courses/cs281/papers/murphy-2007.pdf,
                # Equation 110
                # convert to the params of a PyMC student-t (i.e. integrate out the prior)
                mu = mu_n
                nu = 2.0 * alpha_n
                lam = alpha_n * kappa_n / (beta_n * (kappa_n + 1.0))

                fits[f"{gene}={dataset}"] = (mu, nu, lam, np.sqrt(1 / lam))
        return fits

    def _posterior_predictive_check(self, trace, genes):
        """
        Posterior predictive check for a list of genes trained in the model

        Args:
            trace: PyMC3 trace
            fits: StudentT fits for background dataset/gene expression
            genes: List of genes of interest

        Returns:
            Dictionary of [genes, array of posterior sampling]
        """
        fits = self.t_fits(self.classes)
        d = {}
        for gene in genes:
            d[gene] = self._gene_ppc(trace, fits, gene)
```

```
        return d

    @staticmethod
    def _gene_ppc(trace, fits, gene: str) -> np.array:
        """
        Calculate posterior predictive for a gene

        Args:
            trace: PyMC3 Trace
            fits: StudentT fits for background dataset/gene expression
            gene: Gene of interest

        Returns:
            Random variates representing PPC of the gene
        """
        y_gene = [x for x in trace.varnames if x.startswith(f"{gene}=")]
        y, norm_term = 0, 0
        multiple_backgrounds = "b" in trace.varnames
        for i, y_name in enumerate(y_gene):
            nu, mu, lam, sd = fits[y_name]
            b = trace["b"][:, i] if multiple_backgrounds else 1
            y += (b / sd) * trace[y_name]
            norm_term += b / sd

        return np.random.laplace(loc=(y / norm_term), scale=(trace["eps"] / norm_term))

    def _pearson_pvalue_matrix(self):
        df = self.p
        matrix = []
        df = df[df.Gene.isin(self.genes)]
        ntg_vector = df.num_training_genes.unique()
        for i in ntg_vector:
            row = []
            d1 = df[df.num_training_genes == i]
            d1.index = d1.Gene
            for j in ntg_vector:
                d2 = df[df.num_training_genes == j]
                d2.index = d2.Gene
                # Combine and index by gene
                c = pd.concat([d1.Pval, d2.Pval], axis=1)
                c.columns = [0, 1]
                r, _ = pearsonr(c[0], c[1])
                row.append(r)
            matrix.append(row)
        matrix = pd.DataFrame(matrix, index=ntg_vector, columns=ntg_vector)
        return matrix.apply(lambda x: round(x, 4))

    def plot_genes_by_weights(self):
        _, ax = plt.subplots(figsize=(8, 4))
        sns.lineplot(data=self.w, x="num_training_genes", y="Median", hue="tissue")
        plt.xlabel("Number of Additional Training Genes")
        plt.ylabel("Median Beta Coefficient")
        plt.ylim([0, 1])
        plt.title("Effect of Adding Training Genes on Beta Weights")
```

```
        return ax

    def plot_x(self, gene, tissue):
        self.bg = self._load_df(self.background_path) if self.bg is None else self.bg
        self.sg = self._load_df(self.sample_path) if self.sg is None else self.sg
        df = self._df(gene, tissue)

        _, ax = plt.subplots(figsize=(8, 5))
        plt.axvline(self.sg.loc[self.sample_name][gene], label="N-of-1", c="r")
        gr = df.groupby("num_training_genes")["x"]
        for label, arr in gr:
            sns.kdeplot(arr, label=label)
        sns.kdeplot(self.bg[self.bg.tissue == tissue][gene], shade=True, label="Observed")
        plt.xlabel("Transcripts per Million (log2(TPM +1))")
        plt.ylabel("Density")
        plt.title(f"Effect of n Genes on x Distribution - {gene}")
        return ax

    def plot_posterior(self, gene, tissue):
        self.bg = self._load_df(self.background_path) if self.bg is None else self.bg
        self.sg = self._load_df(self.sample_path) if self.sg is None else self.sg
        df = self._df(gene, tissue)

        _, ax = plt.subplots(figsize=(8, 5))
        plt.axvline(self.sg.loc[self.sample_name][gene], label="N-of-1", c="r")
        gr = df.groupby("num_training_genes")["ppc"]
        for label, arr in gr:
            sns.kdeplot(arr, label=label)
        sns.kdeplot(self.bg[self.bg.tissue == tissue][gene], shade=True, label="Observed")
        plt.xlabel("Transcripts per Million (log2(TPM +1))")
        plt.ylabel("Density")
        plt.title(f"Effect of n Genes on Posterior Distribution - {gene}")
        return ax

    def plot_pval_heatmap(self):
        matrix = self._pearson_pvalue_matrix()
        f, ax = plt.subplots(figsize=(12, 4))
        sns.heatmap(matrix, cmap="Blues", annot=True, linewidths=0.5)
        plt.xlabel("Number of Additional Training Genes")
        plt.ylabel("Number of Additional Training Genes")
        plt.title(
            "Pearson Correlation of Gene P-values (n=85) as Training Genes are Added"
        )
        return ax
```

#### 8.1 Select Sample

Randomly select a tumor sample to run

```
In [19]: tumor = pd.read_hdf('/mnt/data/outlier/tumor.hd5')
         # sample_id = np.random.choice(tumor.index)
         # Selected randomly, but hardcoded here for reproducibility
         sample_id = 'TCGA-DJ-A2PX-01'
```

#### 8.2 Manifest

```
In [3]: out_dir = '/mnt/normsd-outlier-runs/variable-genes'
        path = os.path.join(out_dir, 'manifest.tsv')
        max_genes = [85, 95, 105, 115, 125, 150, 175, 200, 225, 250, 275, 300]
        with open(path, 'w') as f:
            f.write('id\tmax_genes\tout_dir\n')
            for x in max_genes:
                sample_dir = os.path.join(out_dir, str(x))
                f.write(f'{sample_id}\t{x}\t{sample_dir}\n')
```

#### 8.3 Run Samples

```
#!/usr/bin/env bash
source activate toil
python /mnt/gene-outlier-detection/toil/toil-outlier-detection.py \
    --sample /mnt/data/outlier/tumor.hd5 \
    --background /mnt/data/outlier/gtex.hd5 \
    --gene-list /mnt/data/outlier/drug-genes.txt \
    --manifest /mnt/normsd-outlier-runs/variable-genes/manifest.tsv \
    --out-dir /mnt/normsd-outlier-runs/variable-genes/output/ \
    --group tissue \
    --col-skip 5 \
    --num-backgrounds 4 \
    --max-genes 125 \
    --workDir /mnt/ \
    --disable-iter \
    /mnt/jobStore
```

#### 8.4 Genes by Weights

Visualize the effect of additional training genes on the weight that gets assigned to different background tissues.

```
In [2]: g = Genes(
            '/mnt/normsd-outlier-runs/variable-genes/output',
            '/mnt/data/outlier/gtex.hd5',
            '/mnt/data/outlier/tumor.hd5',
            'TCGA-DJ-A2PX-01',
            '/mnt/data/outlier/drug-genes.txt',
        )
```

```
In [3]: out_dir = '/mnt/figures/variable-genes'
        g.plot_genes_by_weights();
        plt.tight_layout()
        plt.savefig(os.path.join(out_dir, 'weights.png'), dpi=300, transparent=True)
```

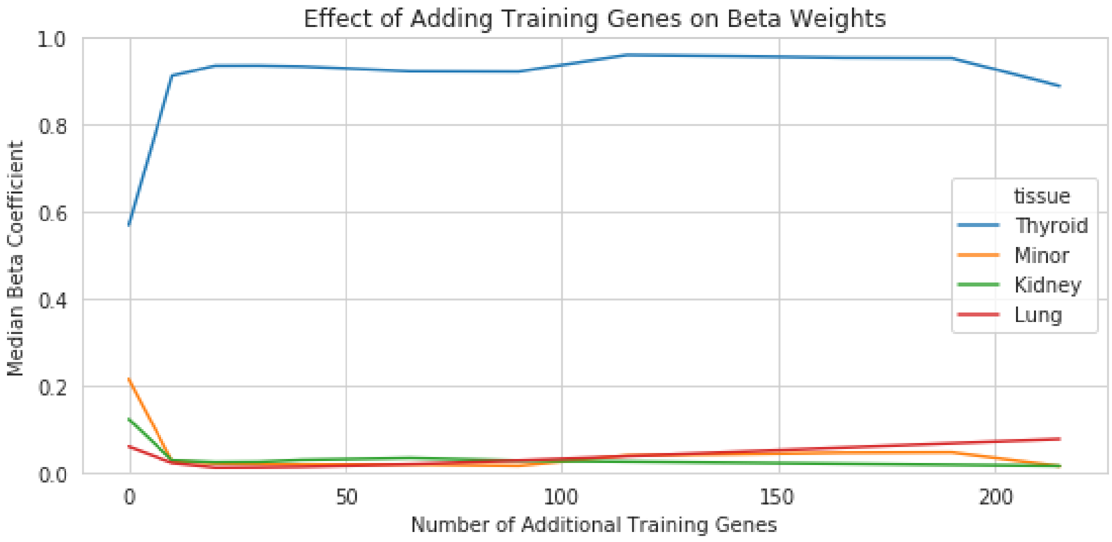

After about ~50 additional training genes, we start losing a little bit of specificity in our model's ability to assign all of the weight to one background tissue, but the model's weight assignment only changes ~5% between the maximum and minimum weight for the matched tissue. Ultimately, we care about this difference in weight assignment on the model parameters.

#### 8.5 Effect of $n$ Genes on $x$ Distribution

Demonstrate the effect of additional training genes on random variates for $x$, which is the expression for a gene/tissue vector. All $x$ distributions are shfited towards the observed N-of-1 sample for that gene, which informs the $x$ distribution during model training.

##### 8.5.1 Random Genes

```
In [4]: ran_genes = np.random.choice(g.genes, 4)
        for gene in ran_genes:
            g.plot_x(gene, 'Thyroid');
```

```
Reading in /mnt/data/outlier/gtex.hd5
Reading in /mnt/data/outlier/tumor.hd5
Loading traces to extract posterior distribution

WARNING (theano.tensor.blas): Using NumPy C-API based implementation for BLAS functions.
```

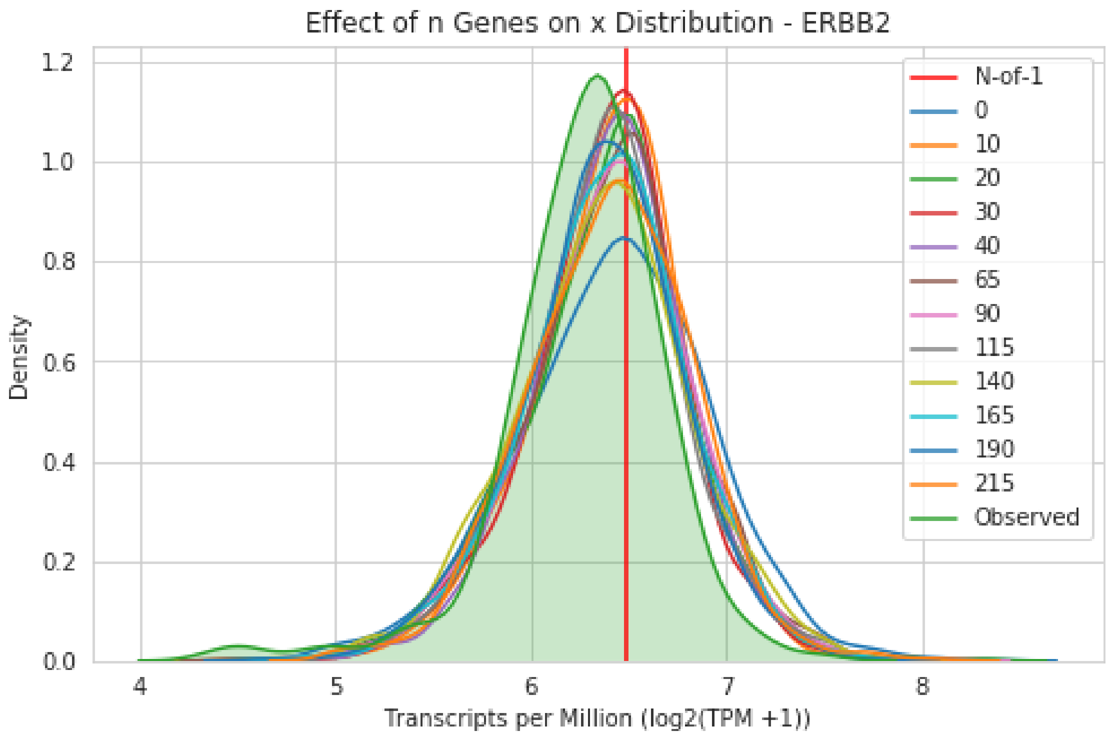
Effect of n Genes on x Distribution - ERBB2
Density
Transcripts per Million (log2(TPM +1))
0.0
0.2
0.4
0.6
0.8
1.0
1.2
4
5
6
7
8
N-of-1
0
10
20
30
40
65
90
115
140
165
190
215
Observed

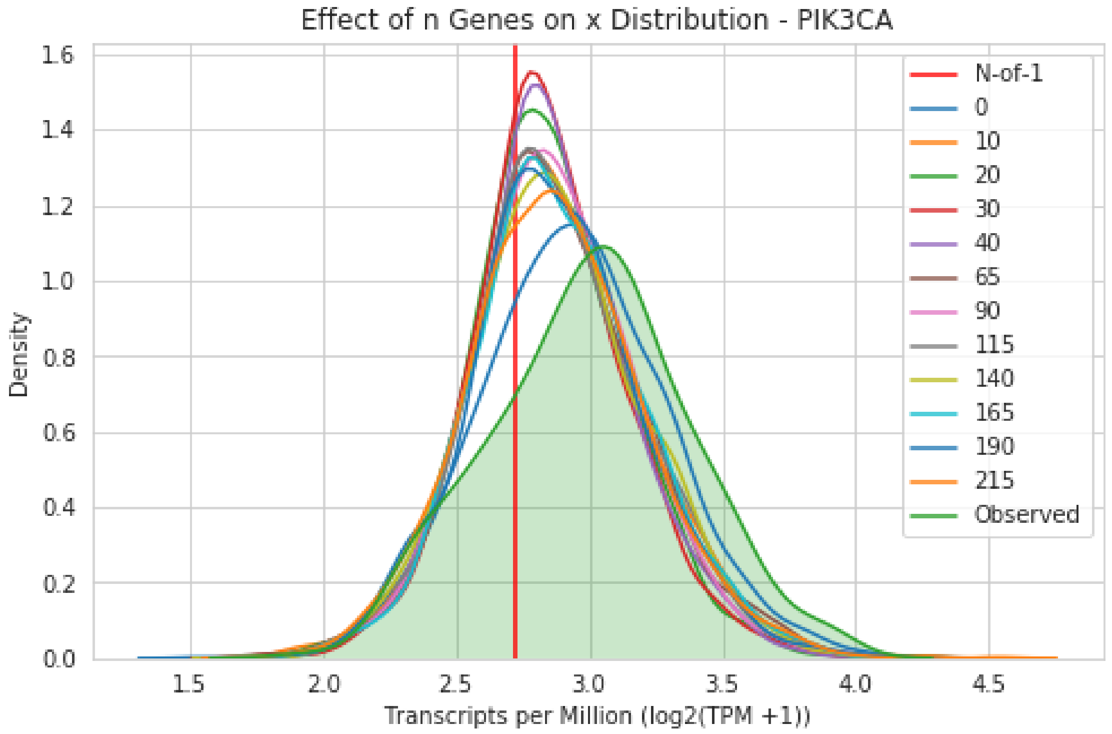
Effect of n Genes on x Distribution - PIK3CA
Density
Transcripts per Million (log2(TPM +1))
0.0
0.2
0.4
0.6
0.8
1.0
1.2
1.4
1.6
1.5
2.0
2.5
3.0
3.5
4.0
4.5
N-of-1
0
10
20
30
40
65
90
115
140
165
190
215
Observed

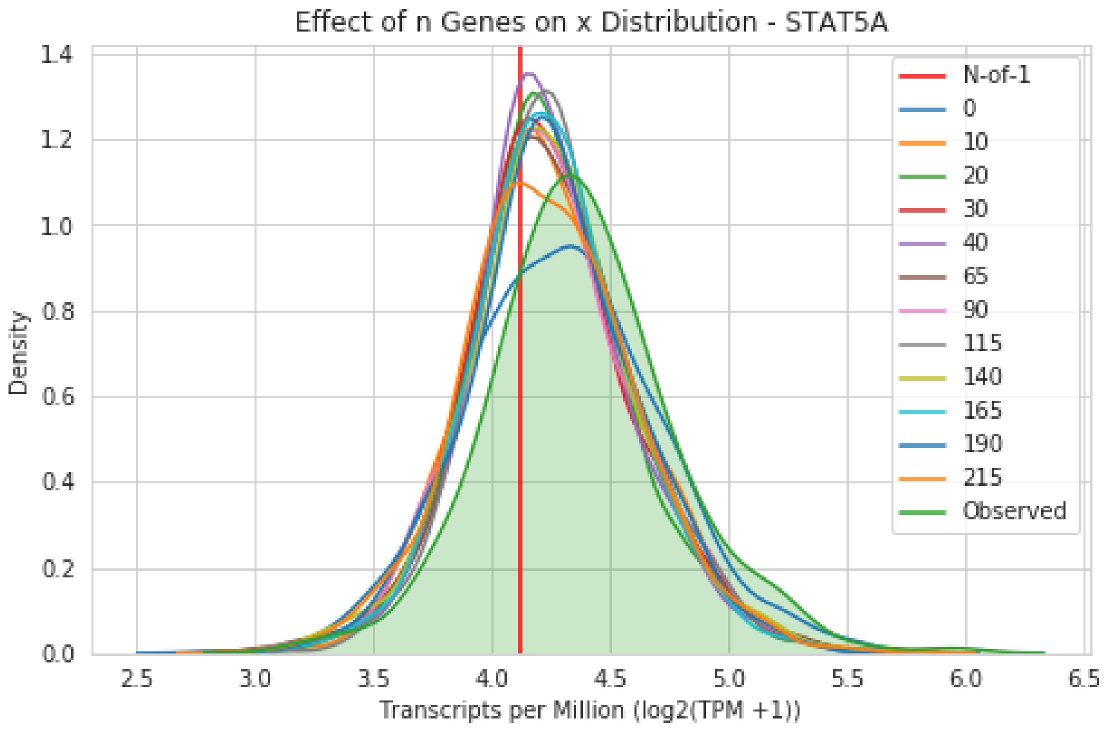
Effect of n Genes on x Distribution - STAT5A
Density
Transcripts per Million (log2(TPM +1))
N-of-1
0
10
20
30
40
65
90
115
140
165
190
215
Observed

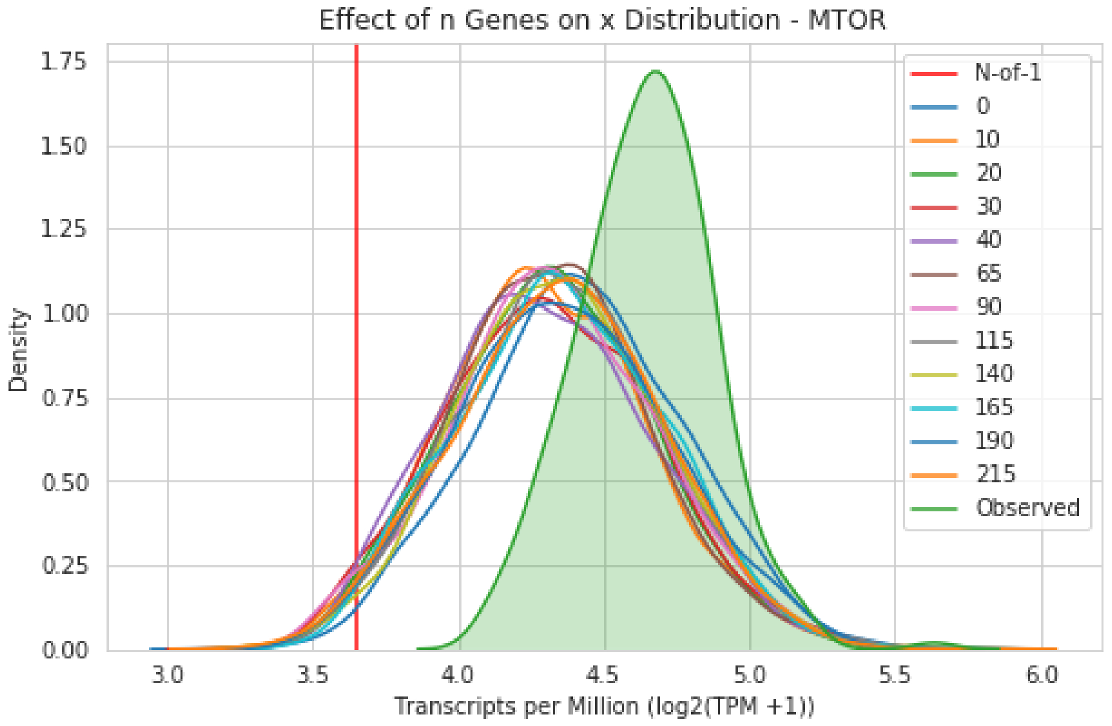
Effect of n Genes on x Distribution - MTOR
Density
Transcripts per Million (log2(TPM +1))
N-of-1
0
10
20
30
40
65
90
115
140
165
190
215
Observed

#### 8.6 Effect of $n$ Genes on Posterior Predictive Distribution

##### 8.6.1 Random Genes

```
In [5]: ran_genes = np.random.choice(g.genes, 4)
        for gene in ran_genes:
            g.plot_posterior(gene, 'Thyroid');
```

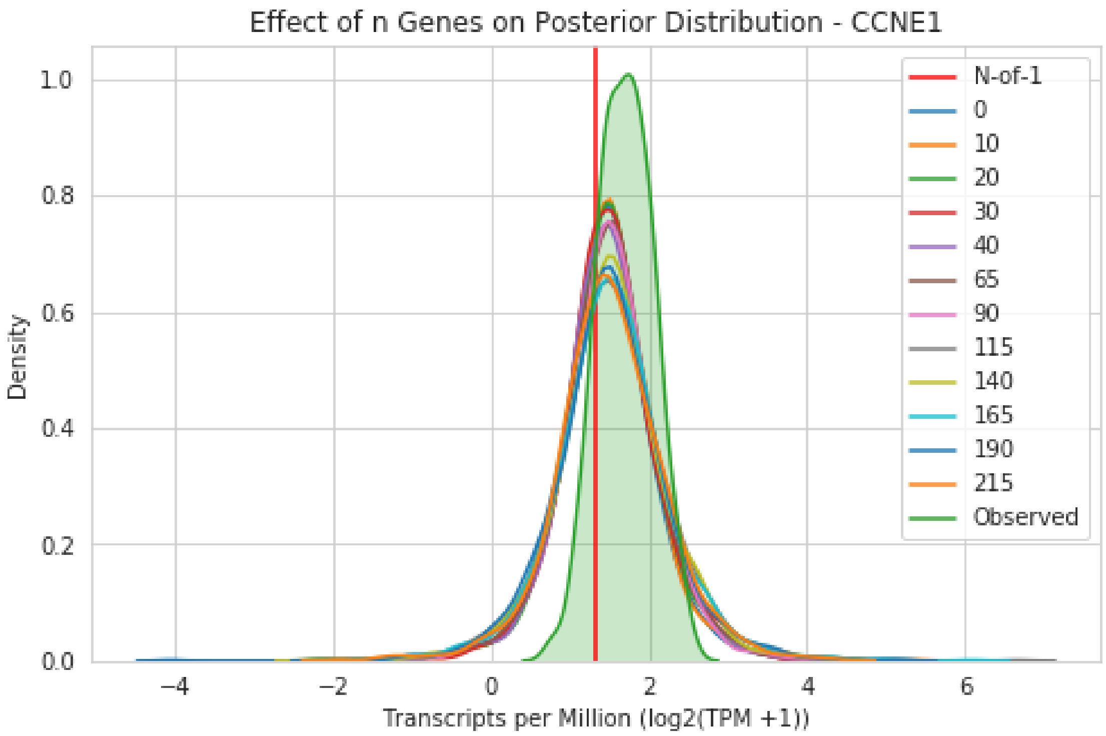

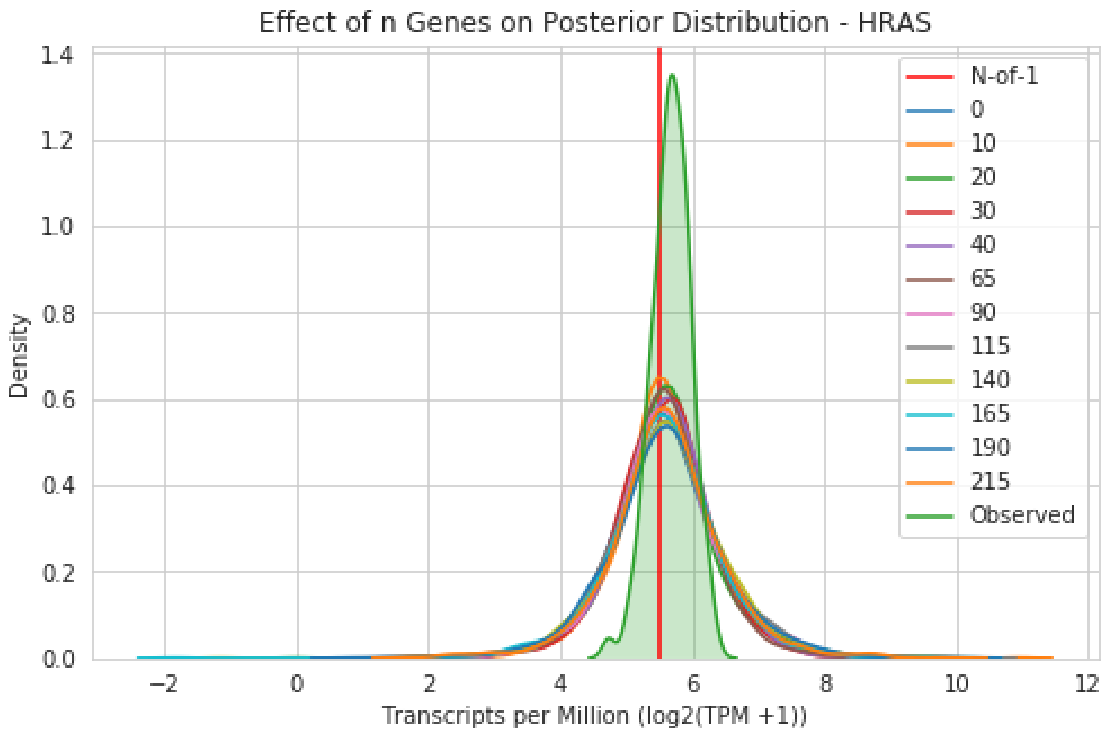
Effect of n Genes on Posterior Distribution - HRAS
Density
Transcripts per Million (log2(TPM +1))
N-of-1
0
10
20
30
40
65
90
115
140
165
190
215
Observed

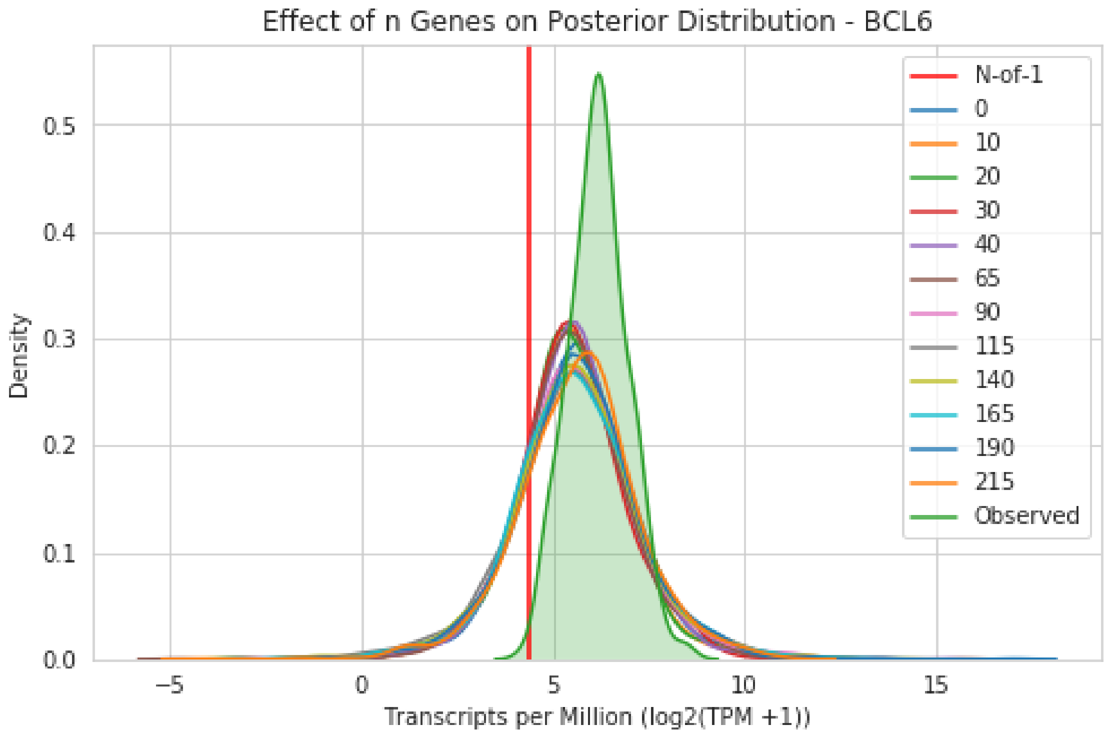
Effect of n Genes on Posterior Distribution - BCL6
Density
Transcripts per Million (log2(TPM +1))
N-of-1
0
10
20
30
40
65
90
115
140
165
190
215
Observed

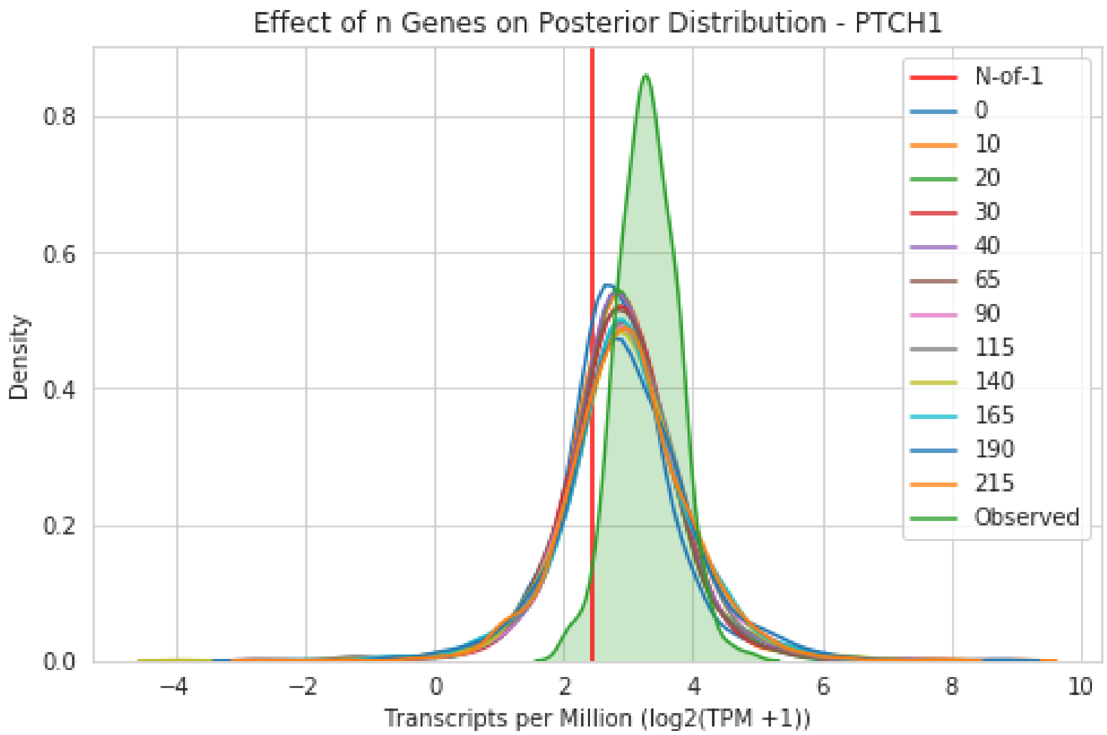

#### 8.7 P-Value Posterior HeatMap

Ultimately, we want to know the effect of additional genes on the p-values the model generates. This can be visualized by calculating the Pearson correlation between all variable runs. For this sample, adding only 10 additional genes (based on ANOVA F-value) produces the same gene p-values as if we had added 215 additional training genes, indicating our model only needs a small number of additional training genes in addition to the gene input list in order to accurately assign beta coefficient weights.

```
In [28]: g.plot_pval_heatmap();
         plt.tight_layout()
         plt.savefig(os.path.join(out_dir, 'heatmap.png'), dpi=300, transparent=True)
```

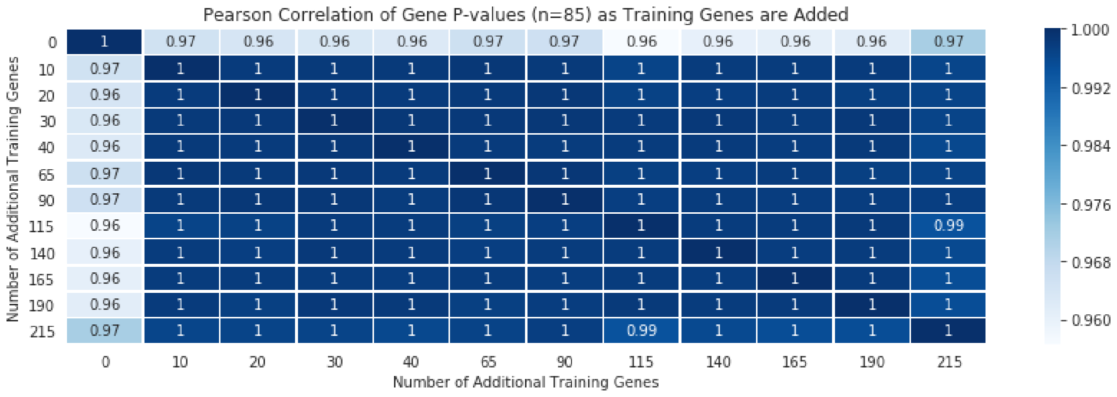

### 9 Effect of $n$ Background Datasets on Model Parameters and Output

This section explores the effect that varying the number of background datasets has on different aspects of model output. We can reuse the same `Genes` class from the previous section.

All data used to generate the following section is avilable at http://courtyard.gi.ucsc.edu/~jvivian/outlier-paper/experiments/variable-backgrounds.tar.gz.

#### 9.1 Select Sample

```
In [2]: # We'll select the same sample as variable-genes here
        sample_id = 'TCGA-DJ-A2PX-01'
```

#### 9.2 Manifest

```
In [3]: out_dir = '/mnt/normsd-outlier-runs/variable-backgrounds/output'
        path = os.path.join(os.path.dirname(out_dir), 'manifest.tsv')
        num_bgs = [1, 2, 3, 4, 5, 6, 7, 8, 9, 10]
        with open(path, 'w') as f:
            f.write('id\tnum_backgrounds\tout_dir\n')
            for x in num_bgs:
                sample_dir = os.path.join(out_dir, str(x))
                f.write(f'{sample_id}\t{x}\t{sample_dir}\n')
```

#### 9.3 Run Samples

```
#!/usr/bin/env bash
source activate toil
python /mnt/gene-outlier-detection/toil/toil-outlier-detection.py \
    --sample /mnt/data/outlier/tumor.hd5 \
    --background /mnt/data/outlier/gtex.hd5 \
    --gene-list /mnt/data/outlier/drug-genes.txt \
    --manifest /mnt/normsd-outlier-runs/variable-backgrounds/manifest.tsv \
    --out-dir /mnt/normsd-outlier-runs/variable-backgrounds/output/ \
    --group tissue \
    --col-skip 5 \
    --num-backgrounds 1 \
    --max-genes 105 \
    --workDir /mnt/ \
    --disable-iter \
    /mnt/jobStore
```

#### 9.4 Background Sets by Weights

```
In [4]: out_dir = '/mnt/normsd-outlier-runs/variable-backgrounds/output'
        g = Genes(
            out_dir,
            '/mnt/data/outlier/gtex.hd5',
            '/mnt/data/outlier/tumor.hd5',
            'TCGA-DJ-A2PX-01',
            '/mnt/data/outlier/drug-genes.txt',
        )
```

```
In [5]: out_dir = '/mnt/figures/variable-backgrounds'
        df = g.w
```

```
plt.figure(figsize=(8, 4))
sns.lineplot(data=df, x='max_genes', y='Median', hue='tissue')
plt.xlabel('Number of Additional Backgrounds')
plt.ylabel('Median Beta Coefficient')
plt.title('Effect of Adding Background Datasets on Beta Weights');
plt.tight_layout()
plt.savefig(os.path.join(out_dir, 'weights.png'), dpi=300, transparent=True)
```

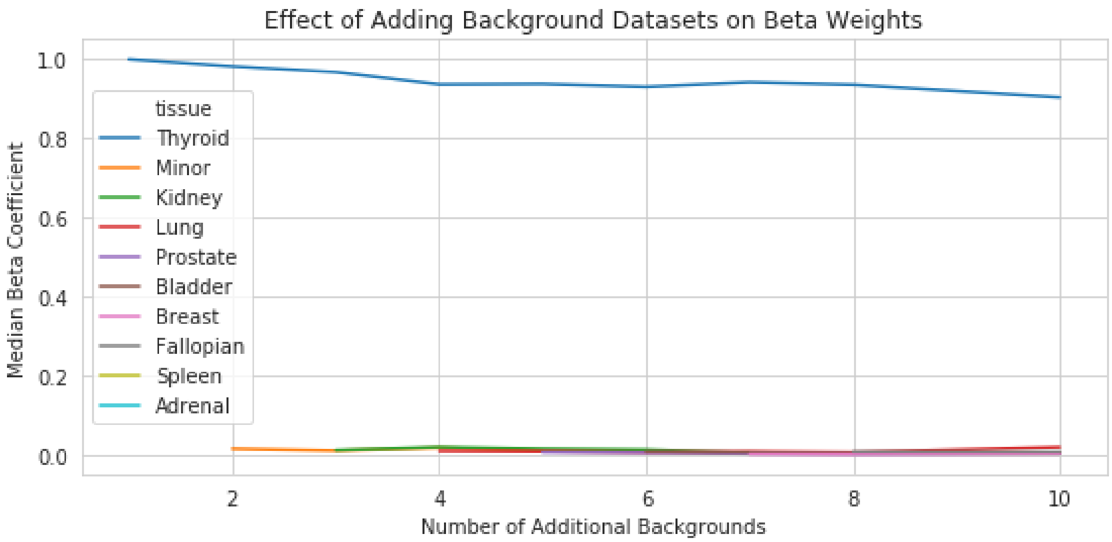

#### 9.5 Effect of $n$ Background Datasets on $x$ Distribution

##### 9.5.1 Random Genes

```
In [15]: ran_genes = np.random.choice(g.genes, 4)
         for gene in ran_genes:
             ax = g.plot_x(gene, 'Thyroid');
             ax.set_title(f'Effect of n Background Datasets on x Distribution - {gene}')
             L = plt.legend()
             for i, n in enumerate(n_bgs):
                 L.get_texts()[i+1].set_text(n)
```

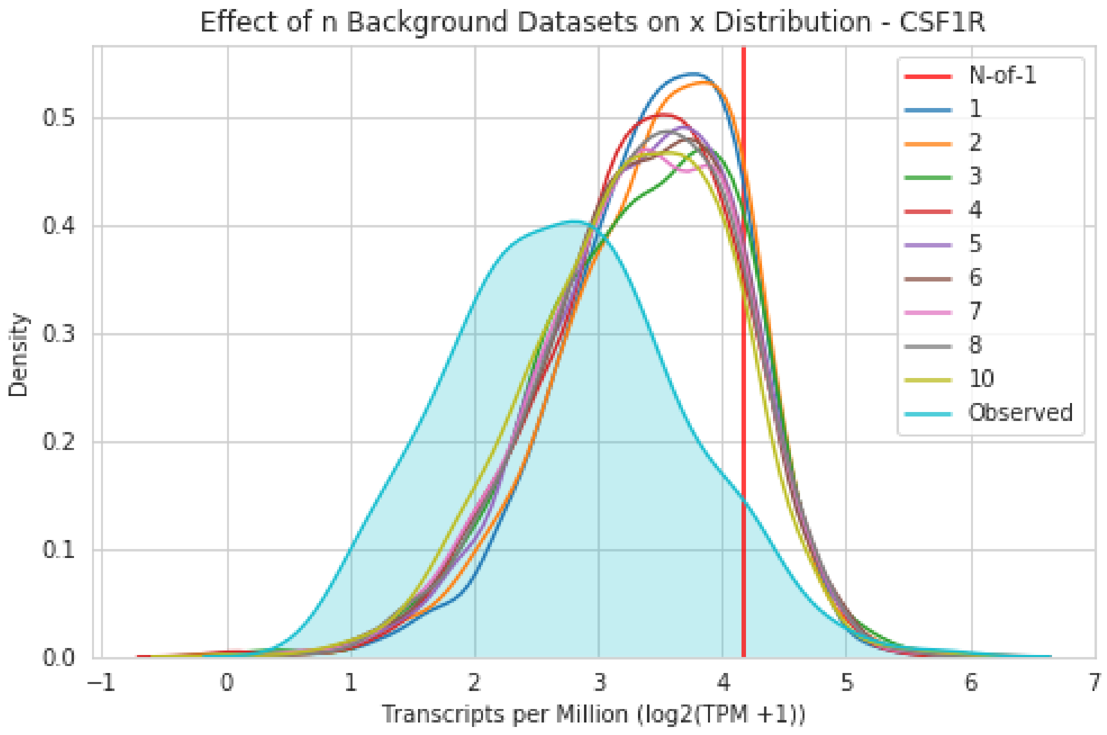
Effect of n Background Datasets on x Distribution - CSF1R
Density
Transcripts per Million (log2(TPM +1))
N-of-1
1
2
3
4
5
6
7
8
10
Observed

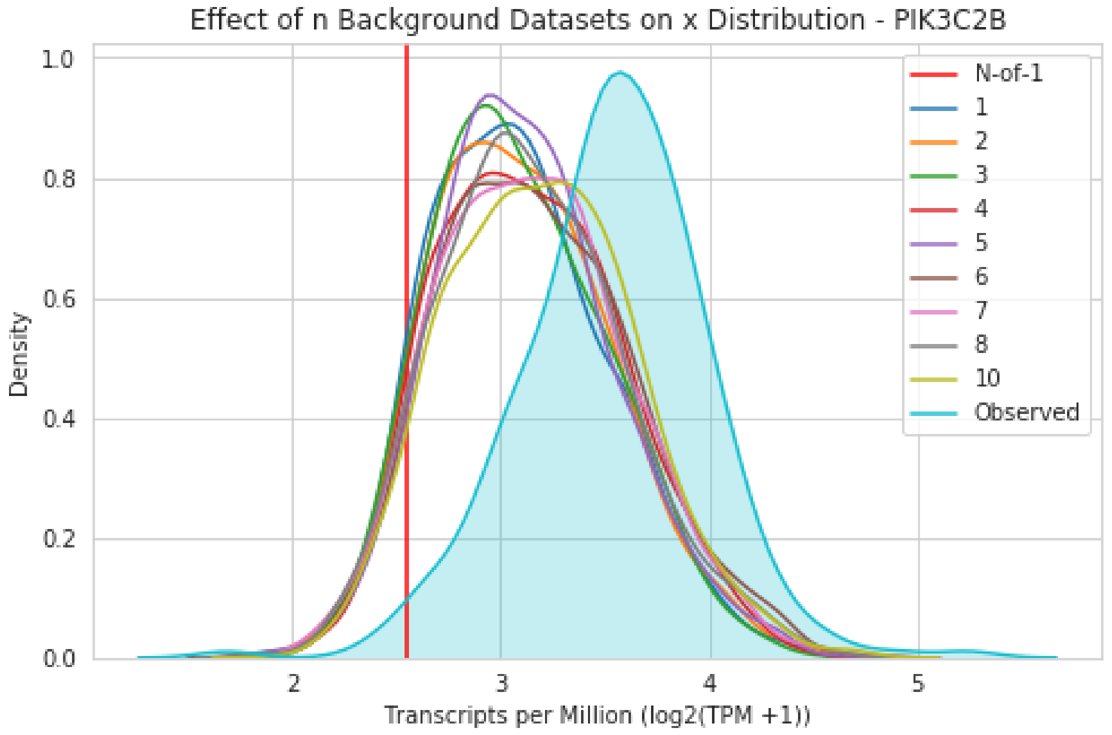
Effect of n Background Datasets on x Distribution - PIK3C2B
Density
Transcripts per Million (log2(TPM +1))
N-of-1
1
2
3
4
5
6
7
8
10
Observed

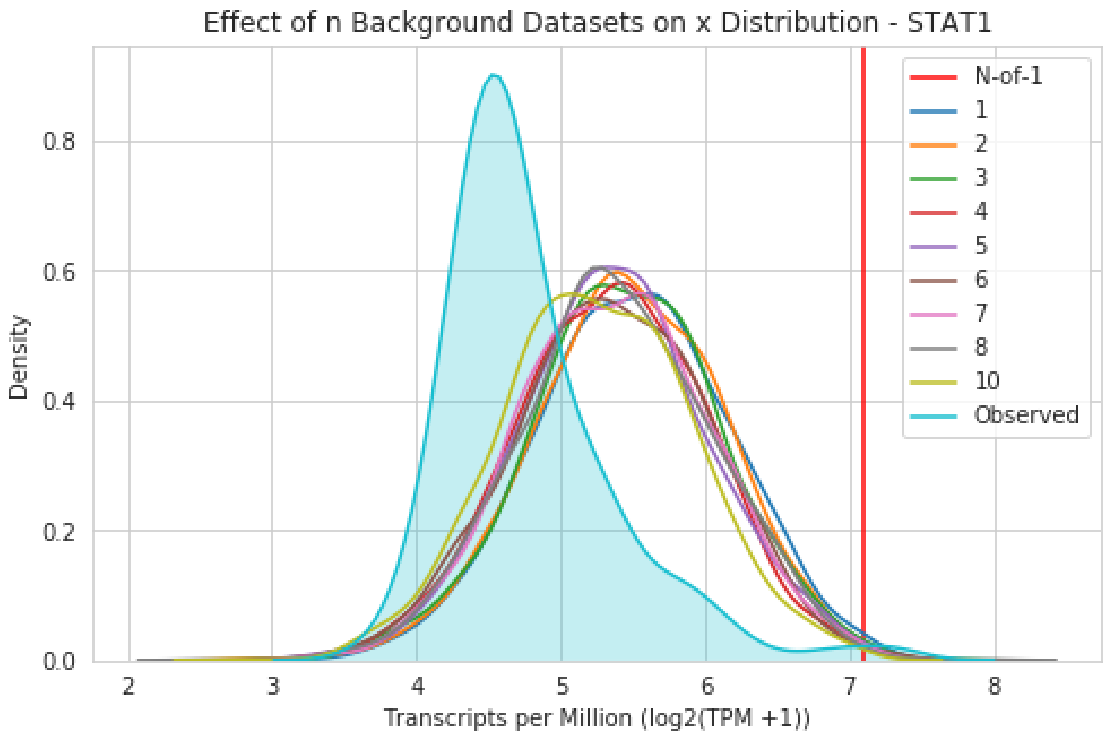
Effect of n Background Datasets on x Distribution - STAT1
Density
Transcripts per Million (log2(TPM +1))
N-of-1
1
2
3
4
5
6
7
8
10
Observed
0.0
0.2
0.4
0.6
0.8
2
3
4
5
6
7
8

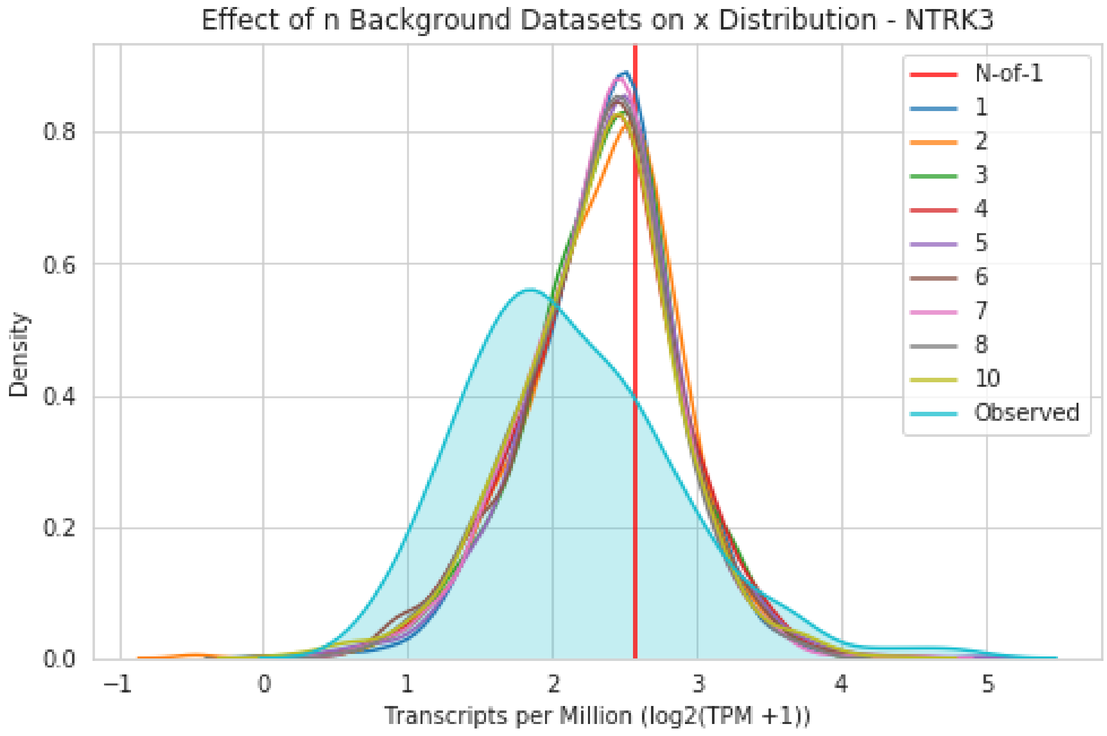
Effect of n Background Datasets on x Distribution - NTRK3
Density
Transcripts per Million (log2(TPM +1))
N-of-1
1
2
3
4
5
6
7
8
10
Observed
0.0
0.2
0.4
0.6
0.8
−1
0
1
2
3
4
5

#### 9.6 Effect of $n$ Background Datasets on Posterior Predictive Distribution

##### 9.6.1 Random Genes

```
In [16]: ran_genes = np.random.choice(g.genes, 4)
         for gene in ran_genes:
             ax = g.plot_posterior(gene, 'Thyroid');
             ax.set_title(f'Effect of n Background Datasets on Posterior Distribution - {gene}')
             L = plt.legend()
             for i, n in enumerate(n_bgs):
                 L.get_texts()[i+1].set_text(n)
                 L.get_texts()[-1].set_text('Posterior')
```

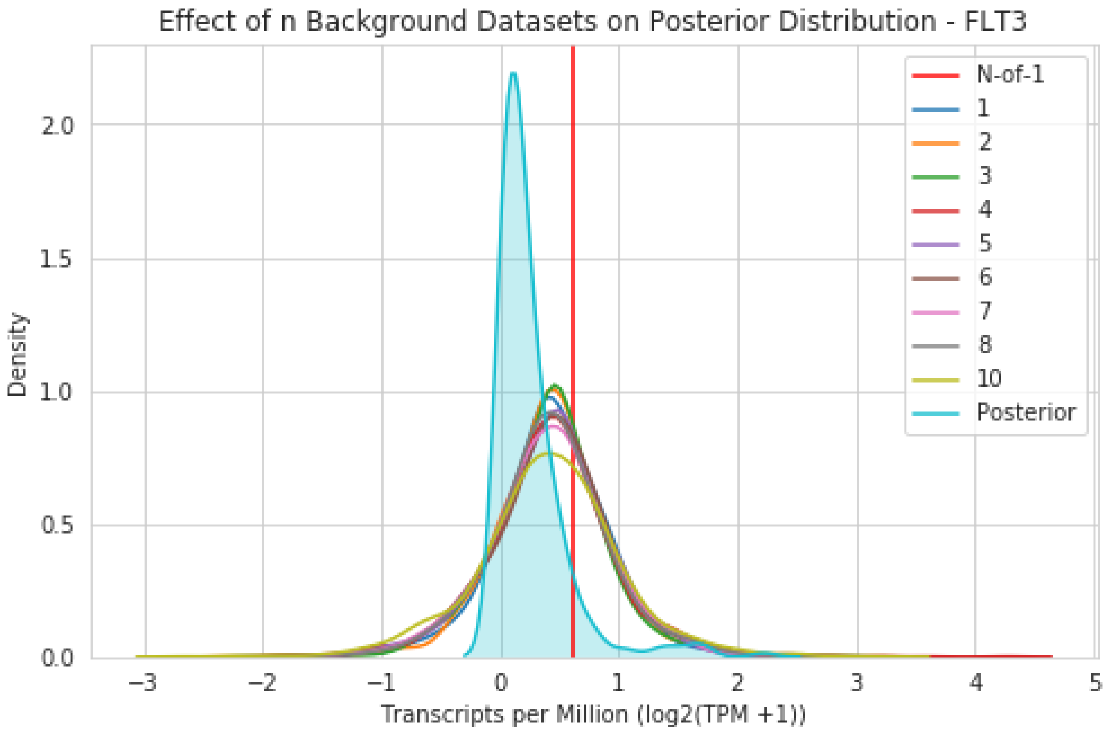

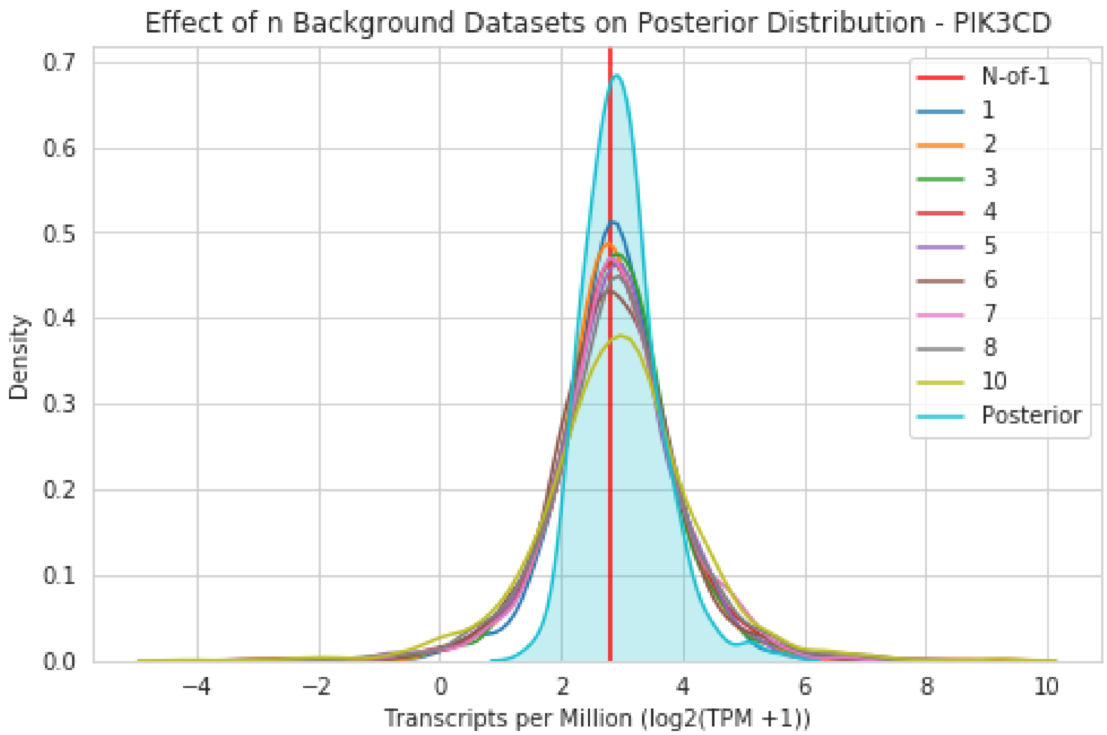

Effect of n Background Datasets on Posterior Distribution - PIK3CD
Density
Transcripts per Million (log2(TPM +1))
N-of-1
1
2
3
4
5
6
7
8
10
Posterior

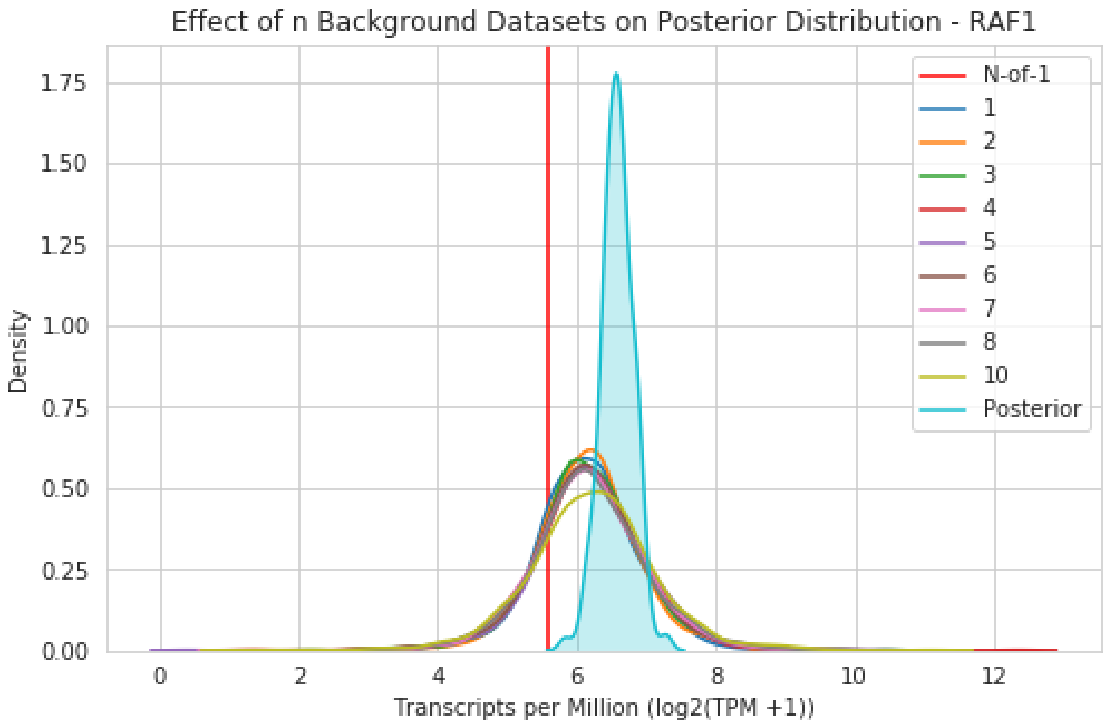

Effect of n Background Datasets on Posterior Distribution - RAF1
Density
Transcripts per Million (log2(TPM +1))
N-of-1
1
2
3
4
5
6
7
8
10
Posterior

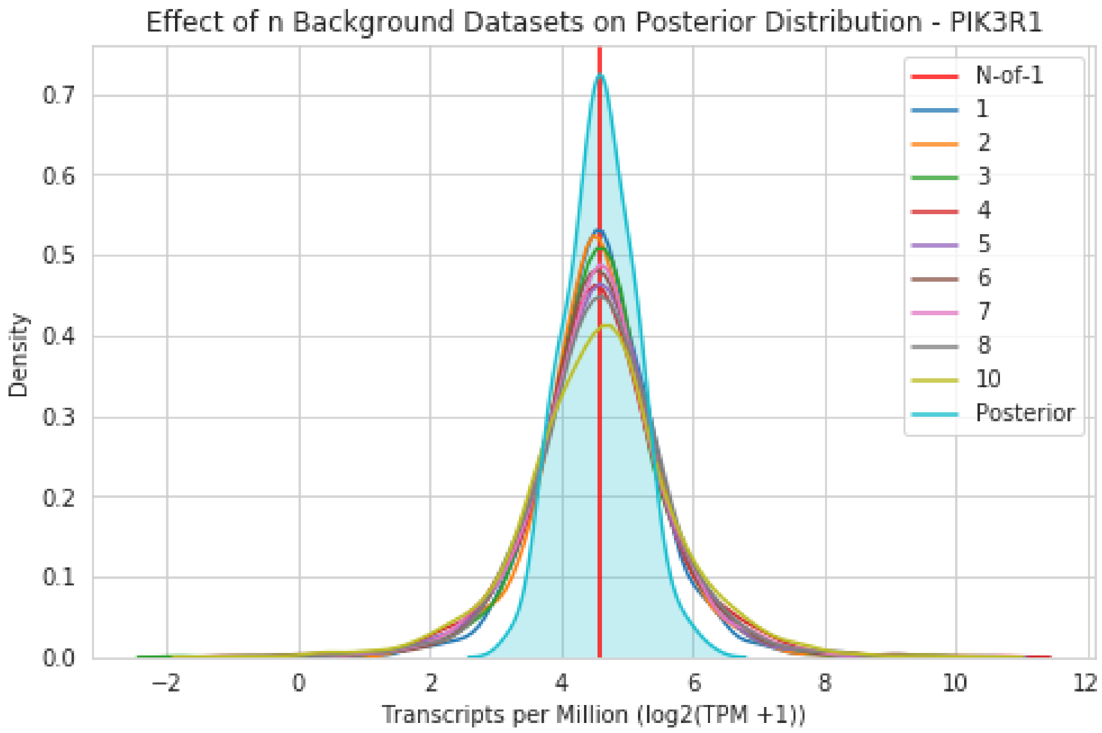

##### 9.7 P-value Posterior Heatmap

Similar to the previous section, we want to know the effect of additional background training sets on the p-values the model generates. This can be visualized by calculating the Pearson correlation between all variable runs. For this sample, the first background dataset was the matched normal from GTEx. Since the model assigns 100% of the weight to the matched tissue, the pearson correlation between gene p-values remains static regardless of the number of additional background datasets that are added.

```
In [17]: matrix = g._pearson_pvalue_matrix()
         matrix.index = range(1, 10)
         matrix.columns = range(1, 10)
```

```
In [18]: plt.figure(figsize=(12, 4))
         sns.heatmap(matrix, cmap='Blues', annot=True, linewidths=0.5)
         plt.xlabel('Number of Background Datasets')
         plt.ylabel('Number of Background Datasets')
         plt.title('Pearson Correlation of Gene P-values (n=85) as Background Datasets are Added');
         plt.tight_layout()
         plt.savefig(os.path.join(out_dir, 'heatmap.png'), dpi=300, transparent=True)
```

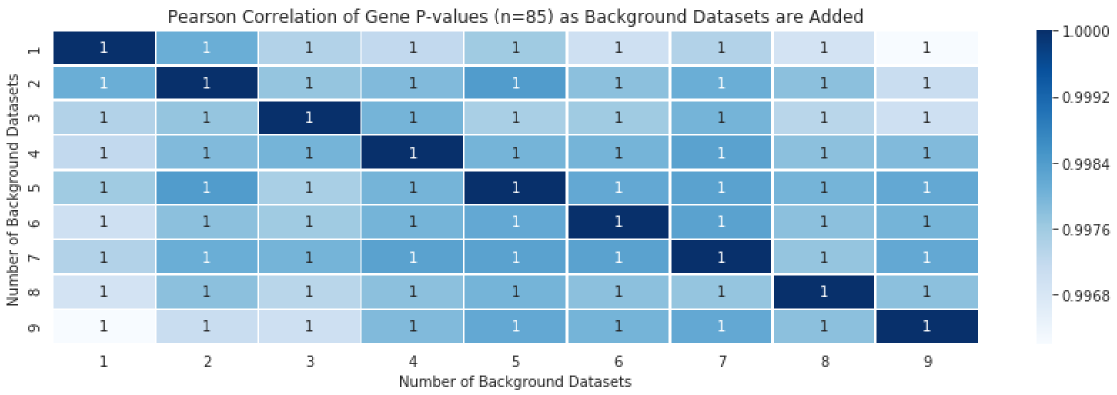

### 10 Software Engineering

The codebase for our outlier detection method is open source (https://github.com/jvivian/gene-outlier-detection/) and available as a Python package (`pip install --pre gene-outlier-detection`), a Docker container (`docker pull jvivian/gene-outlier-detection`), and a Toil workflow (see Github repository). The software provides a plethora of output: ranked pairwise distances of the background datasets as compared to the N-of-1 sample, a serialized PyMC3 trace and model for reproducibility, trace plots for important model parameters to confirm model convergence, $\beta$ weight table and plot to see confidence of model in assignment of weight to the background datasets, and posterior predictive p-values for every gene of interest (Figure 1). The software also includes an **_info** subdirectory that contains Pearson correlations between iterative runs, p-values for those runs, and a **_run_info.tsv** table that contains every parameter setting used in that run of the model in addition to runtime.

The PyMC3 model and trace can be directly examined from the serialized model:

```
import pickle
with open(pkl_path, 'rb') as buff:
    data = pickle.load(buff)
model, trace = data['model'], data['trace']
```

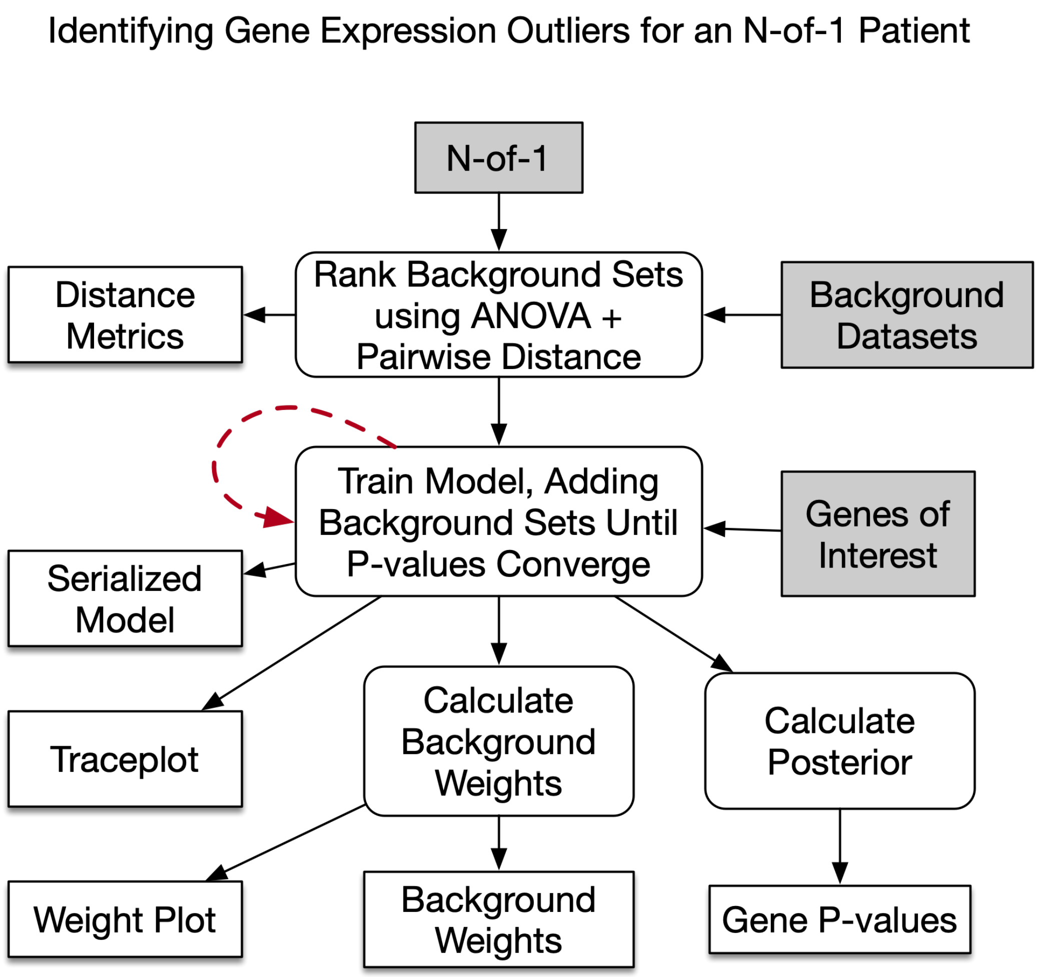

Figure 1: Graph of the outlier modeling software. Dark grey boxes indicate inputs and shadowed boxes indicate outputs. Background datasets are first ranked by calculating pairwise distance from the n-of-1 samples to the top 10% ANOVA genes (section 7). By default, background datasets are iteratively added until the provided genes converge with a p-value $> 0.99$. The software outputs a serialized

#### 11 Data Collection and Processing

The following sections describe how the expression data and metadata were collected and processed.

The easiest way to obtain the expression and metadata used in these experiments is from this URL `http://courtyard.gi.ucsc.edu/~jvivian/outlier-paper/expression-matrices/`. Each HD5 matrix has 5 metadata columns: **id, tissue, subtype, tumor,** and **label**. 'id' corresponds to the sample id, 'tissue' corresponds to the primary tissue type, 'subtype' corresponds to the more granuar tumor or tissue subtype (if available), 'tumor' is a boolean column for whether the sample is a cancer as opposed to normal sample, and 'label' designates which dataset the sample is from. The remaining columns are genes and the values are are expression data from RSEM in the units $log2(TPM + 1)$. The raw table is available from the UC Santa Cruz Xena browser.

#### 11.1 TCGA and GTEx Gene Expression Matrix

Process Xena TPM table for downstream analysis - Subset protein-coding genes - Map genes - Reverse normalization to standard TPM

In [4]:
```
import os
import requests

import numpy as np
import pandas as pd
import rnaseq_lib as r
import holoviews as hv
hv.extension('bokeh', logo=False)
```

##### 11.1.1 Input Data

Read in TPM matrix from Xena

In [ ]:
```
exp_path = '/mnt/data/Expression/TcgaTargetGtex_rsem_gene_tpm'
%time exp = pd.read_csv(exp_path, sep='\t', index_col=0)
```

In [ ]:
```
exp.head()
```

Distribution of Xena normalized TPM mean for genes and samples

In [ ]:
```
%%opts Distribution [xrotation=45]
gene_tpm = hv.Distribution(exp.mean(axis=1).tolist(), kdims='Gene TPM Mean')
sample_tpm = hv.Distribution(exp.mean().tolist(), kdims='Sample TPM Mean')
hv.Layout([gene_tpm, sample_tpm], label='Xena TPM Dataframe')
```

Read in metadata for TCGA and GTEx

In [ ]:
```
met_path = '/mnt/data/Metadata/tcga_gtex_metadata_intersect.tsv'
met = pd.read_csv(met_path, sep='\t', index_col=0)
# Drop duplicate entries
met = met[~met.index.duplicated()]
met.head()
```

##### 11.1.2 Subset for TCGA and GTEx

In [ ]:
```
samples = [x for x in exp.columns if x in met.id.tolist()]
df = exp[samples]
df.shape
```

##### 11.1.3 Reverse XENA Normalization

Reverse Xena normalization (log2(x + 0.001)) to get Transcripts per Million (TPM)

In [ ]:
```
df = df.apply(lambda x: 2**x - 0.001)
df.head(2)
```

TPM Distribution of genes and samples

In [ ]:
```
%%opts Distribution [xrotation=45]
gene_tpm = hv.Distribution(df.mean(axis=1).tolist(), kdims='Gene TPM Mean')
sample_tpm = hv.Distribution(df.mean().tolist(), kdims='Sample TPM Mean')
hv.Layout([gene_tpm, sample_tpm], label='Transcripts per Million')
```

Set all negative values to zero, which exist due to floating point errors

```
In [ ]: df[df < 0] = 0
```

~60,000 gene annotations in gencode v23. TPM should sum to a million per sample.

```
In [ ]: %%opts Distribution [xrotation=45]
        sums = df.sum(axis=0).tolist()
        hv.Distribution(sums, kdims='Sum(TPM)') + hv.BoxWhisker(sums)
```

##### 11.1.4 Map Gene Names

Read in ENSMBL -> HUGO mapping table

```
In [ ]: hugo_path = '/mnt/data/Metadata/ensembl_to_hugo.tsv'
        hugo = pd.read_csv(hugo_path, sep='\t', index_col=0).sort_index()

        # Remove duplicates
        hugo = hugo[~hugo.index.duplicated(keep='first')]
        hugo.head()
```

Reindex dataframe with hugo names

```
In [ ]: df.index = hugo.reindex(df.index).geneName.values
        df = df[df.index.notnull()]
        df.head()
```

Take aggregate sum for duplicate hugo names as multiple ensemble genes map to the same hugo ID

```
In [ ]: df = df.groupby(level=0).aggregate(np.sum)
        df.shape
```

##### 11.1.5 Save as HDF

```
In [ ]: df = df.T
```

Cast types as numpy float32 for speed

```
In [26]: df = df.astype(np.float32)
```

```
In [27]: df.to_hdf('/mnt/data/Expression/tcga_gtex_tpm.hd5', key='exp')
```

#### 11.2 TCGA and GTEx Metadata Collation

Goal: Create a combined metadata table that spans both TCGA and GTEx.
TSVs were obtained from here, and publically rehosted on Synapse.

- TCGA: syn7248855
- GTEx: syn10142937

```
In [245]: %matplotlib inline
          import os
          import pandas as pd
          from pandas.tools.plotting import scatter_matrix
          import numpy as np
          import seaborn as sns
          import matplotlib.pyplot as plt
          import pickle

          sns.set_style('whitegrid')
```

##### 11.2.1 Read Inputs

```
In [222]: tcga = pd.read_csv('input_data/tcga-sample-metadata.tsv', sep='\t',
                             low_memory=False)
          gtex = pd.read_csv('input_data/gtex-sample-metadata.tsv', sep='\t')
```

How many metadata columns are there?

```
In [223]: print 'TCGA: {}\t GTEx: {}'.format(len(tcga.columns),
                                             len(gtex.columns))
```

```
TCGA: 858         GTEx: 301
```

...that's quite a few. Because most columns are likely uninformative, filter the samples by RNA-Seq and examine the remaining columns.

##### 11.2.2 TCGA

Locate what column the sample barcodes are in

```
In [224]: tcga[tcga.apply(
                  lambda r: r.str.contains(
                      'TCGA-DQ-5630', case=False).any(), axis=1)].index
```

```
Out[224]: Int64Index([2470], dtype='int64')
```

Locate column(s) to filter by

```
In [225]: print [str(x) for x in tcga.iloc[2470]].index('RNA-Seq')
```

```
33
```

```
In [226]: tcga.columns[126]   # "Barcode" in original metadata
```

```
Out[226]: 'gdc_cases.samples.portions.analytes.aliquots.submitter_id'
```

```
In [227]: tcga.columns[33]   # filter samples by the term: 'RNA-Seq'
```

```
Out[227]: 'gdc_experimental_strategy'
```

Subset by experimental strategy

```
In [228]: tcga = tcga[tcga['gdc_experimental_strategy'] == 'RNA-Seq']
          print 'Samples: {}\tFeatures: {}'.format(*tcga.shape)
```

```
Samples: 11284        Features: 858
```

Sift through 858 columns using the famous GSD (Graduate Student Descent) algorithm

##### 11.2.3 List of columns to keep for TCGA metadata

Sample Information

- gdc_cases.samples.submitter_id

- reads_downloaded
- paired_end
- mapped_read_count
- auc
- gdc_platform
- gdc_file_size
- gdc_experimental_strategy
- gdc_center.code
- gdc_center.name

- gdc_center.short_name
- gdc_data_category
- gdc_cases.tissue_source_site.name
- gdc_cases.samples.portions.analytes.aliquots.concentration
- gdc_cases.samples.portions.analytes.a260_a280_ratio
- cgc_sample_country_of_sample_procurement

Patient Information

- gdc_cases.demographic.gender
- gdc_cases.demographic.year_of_birth
- gdc_cases.demographic.race
- gdc_cases.demographic.ethnicity
- gdc_cases.samples.initial_weight
- gdc_cases.diagnoses.days_to_death
- gdc_cases.diagnoses.age_at_diagnosis
- gdc_cases.exposures.cigarettes_per_day

- gdc_cases.exposures.alcohol_history
- gdc_cases.exposures.weight
- gdc_cases.exposures.years_smoked
- gdc_cases.exposures.height
- cgc_case_year_of_diagnosis

Tumor Information

- gdc_cases.project.name
- gdc_cases.project.primary_site
- gdc_cases.diagnoses.tumor_stage
- gdc_cases.diagnoses.vital_status
- gdc_cases.samples.sample_type
- cgc_case_new_tumor_event_after_initial_treatment
- cgc_case_pathologic_stage

Select, subset, and rename dataframe

```
In [229]: tcga = tcga[['gdc_cases.samples.submitter_id',
                       'reads_downloaded', 'paired_end',
                       'mapped_read_count', 'auc',
                       'gdc_platform', 'gdc_file_size',
```

```
                       'gdc_experimental_strategy',
                       'gdc_center.short_name',
                       'gdc_data_category',
                       'gdc_cases.tissue_source_site.name',
                       'gdc_cases.samples.portions.analytes.aliquots.concentration',
                       'gdc_cases.samples.portions.analytes.a260_a280_ratio',
                       'cgc_sample_country_of_sample_procurement',
                       'gdc_cases.demographic.gender',
                       'gdc_cases.demographic.year_of_birth',
                       'gdc_cases.demographic.race',
                       'gdc_cases.demographic.ethnicity',
                       'gdc_cases.samples.initial_weight',
                       'gdc_cases.diagnoses.days_to_death',
                       'gdc_cases.diagnoses.age_at_diagnosis',
                       'gdc_cases.exposures.cigarettes_per_day',
                       'gdc_cases.exposures.alcohol_history',
                       'gdc_cases.exposures.weight',
                       'gdc_cases.exposures.years_smoked',
                       'gdc_cases.exposures.height',
                       'cgc_case_year_of_diagnosis',
                       'gdc_cases.project.name',
                       'gdc_cases.project.primary_site',
                       'gdc_cases.diagnoses.tumor_stage',
                       'gdc_cases.diagnoses.vital_status',
                       'gdc_cases.samples.sample_type',
                       'cgc_case_new_tumor_event_after_initial_treatment',
                       'cgc_case_pathologic_stage']]
        tcga.index = tcga['gdc_cases.samples.submitter_id']
        tcga.columns = ['submitter_id', 'reads_downloaded', 'paired_end',
                        'mapped_read_count',
                        'auc', 'platform', 'file_size',
                        'experimental_strategy',
                        'center_short', 'data_category',
                        'tissue_source_site',
                        'aliquots_concentration', 'a260_a280_ratio',
                        'country_of_sample_procurement',
                        'gender', 'year_of_birth', 'race', 'ethnicity',
                        'initial_weight',
                        'days_to_death', 'age_at_diagnosis',
                        'cigarettes_per_day', 'alcohol_history',
                        'weight', 'years_smoked', 'height',
                        'year_of_diagnosis', 'project_name',
                        'primary_site', 'tumor_stage', 'vital_status',
                        'sample_type',
                        'new_tumor_event_after_initial_treatment',
                        'pathologic_stage']
        tcga.index = tcga['submitter_id']
        tcga.index.name = None
```

Drop duplicate rows based on index

```
In [230]: tcga = tcga[~tcga.index.duplicated(keep='first')]
```

```
In [231]: print 'Samples: {}\tFeatures: {}'.format(*tcga.shape)
```

```
Samples: 11190        Features: 34
```

**Data Cleanup** Look for features with values that need NAN replacements, sparse / uninformative, etc

```
In [232]: for c in tcga.columns:
              print c, len(tcga[c].unique())
              if len(tcga[c].unique()) == 1:
                  print '\tdropping: ' + c + '\t' + str(tcga[c][0])
                  tcga.drop(c, axis=1, inplace=True)
```

```
submitter_id 11190
reads_downloaded 11189
paired_end 1
        dropping: paired_end        True
mapped_read_count 11190
auc 11190
platform 2
file_size 11190
experimental_strategy 1
        dropping: experimental_strategy        RNA-Seq
center_short 3
data_category 1
        dropping: data_category        Raw sequencing data
tissue_source_site 217
aliquots_concentration 19
a260_a280_ratio 108
country_of_sample_procurement 33
gender 3
year_of_birth 91
race 7
ethnicity 4
initial_weight 221
days_to_death 1579
age_at_diagnosis 7797
cigarettes_per_day 186
alcohol_history 3
weight 379
years_smoked 64
height 135
year_of_diagnosis 34
project_name 33
primary_site 26
tumor_stage 21
vital_status 4
sample_type 7
new_tumor_event_after_initial_treatment 3
pathologic_stage 21
```

convert string nones to nans

```
In [233]: tcga.replace('not reported', np.nan, inplace=True)
          tcga.replace('None', np.nan, inplace=True)
```

Fix A260 / a280 ratios. Two samples have a260's greater than 10 (180 and 187.5 respectively). We'll set these to NaN since we don't actually know their score should be.

```
In [234]: for i in tcga[tcga['a260_a280_ratio'] > 10].index:
              tcga.loc[i, 'a260_a280_ratio'] = np.nan
```

Convert kg to lb for weight

```
In [235]: tcga['weight'] = tcga['weight'].apply(lambda x: x * 2.2046)
```

Save TCGA metadata

```
In [236]: tcga.to_csv('tcga-metadata-cleaned.tsv', sep='\t')
```

##### 11.2.4 Combined Columns

Metadata that spans both datasets

- ID
  - submitter_id
  - SAMPID
- Reads
  - reads_downloaded
  - spots (??)
- size
  - file_size (bytes?, divide by a million)
  - size_MB
- platform
  - platform
  - Model
- sex
  - gender
  - Sex
- tissue
  - primary_site
  - Body_Site
- Sequencing
  - center_short
  - CenterName
- weight
  - weight
  - WGHT
- height
  - height
  - HGHT
- mapped

- mapped_read_count
  - SMMPPD
- race
  - race
  - RACE (3=white, 2=african american, 1=asian, 98,98,4=other)
- age
  - year_of_birth (2010 - year_of_birth = age)
  - AGE
- ethnicity
  - ethnicity
  - ETHNICITY (don't know code)
- qc (Need transformation factor)
  - a260_a280_ratio
  - SMRIN

Create combined dataframe

```
In [ ]: colnames = ['id', 'reads', 'size_MB', 'platform', 'sex', 'tissue',
                    'seq_site', 'weight', 'height', 'mapped_reads', 'race',
                    'age', 'qc']

        tcga_sub = tcga[['submitter_id', 'reads_downloaded', 'file_size',
                         'platform', 'gender', 'primary_site','center_short',
                         'weight', 'height', 'mapped_read_count',
                         'race', 'year_of_birth', 'a260_a280_ratio']]
        tcga_sub.columns = colnames

        gtex_sub = gtex[['SAMPID', 'spots', 'size_MB', 'Model', 'Sex',
                         'Body_Site', 'CenterName', 'WGHT', 'HGHT', 'SMMPPD',
                         'RACE', 'AGE', 'SMRIN']]
        gtex_sub.columns = colnames

        # TCGA A280 conversion to RIN space
        # Model as a piecewise linear transformation.
        # For values <=2.0    y = 20x - 30
        # For values >2.0     y = -20x + 50
        vals = []
        for v in tcga_sub.qc:
            if v <= 2.0:
                vals.append(20*v - 30)
            else:
                vals.append(-20*v + 50)
        tcga_sub.qc = [x if x > 0 else 0 for x in vals]

        # Convert TCGA height from cm to in
        tcga_sub.height = tcga_sub.height.apply(lambda x: x*1.0 / 2.54)
        # Fix outliers
        tcga_sub.height = [x if x > 40 else np.nan for x in tcga_sub.height]

        # Convert size to MB for TCGA
        tcga_sub.size_MB = tcga_sub.size_MB.apply(lambda x: (x*1.0) / 1024 / 1024)
```

```python
# Convert year of birth to age
tcga_sub.age = tcga_sub.age.apply(lambda x: 2010 - x)

# Truncate last letter of TCGA ID
tcga_sub['id'] = tcga_sub.id.apply(lambda x: x[:-1])

# Create combined manifest
df = pd.concat([tcga_sub, gtex_sub], axis=0)

# Add dataset column
df['dataset'] = ['tcga' if x.startswith('TCGA')
                 else 'gtex' for x in df['id']]

# Add Tumor column
df['tumor'] = ['no' if x.endswith('-11') or x.startswith('GTEX')
               else 'yes' for x in df['id']]

# Clean values for certain columns
df.platform.replace('Illumina HiSeq 2000', 'Illumina HiSeq',
                    inplace=True)

# Generalize tissue labels
df.tissue = df.tissue.apply(lambda x: x.split()[0])
df.tissue.replace('Small', 'Small_intestine', inplace=True)
df.tissue.replace('Soft', 'Soft_tissue', inplace=True)
df.tissue.replace('Bone', 'Bone_marrow', inplace=True)
df.tissue.replace('Colorectal', 'Colon', inplace=True)

# Replace numbers with string words
df.race.replace(3, 'white', inplace=True)
df.race.replace(2, 'black or african american', inplace=True)
df.race.replace(1, 'asian', inplace=True)
df.race.replace(98, 'other', inplace=True)
df.race.replace(99, 'other', inplace=True)
df.race.replace(4, 'other', inplace=True)

# Set index to ID column
df.index = df['id']
df.index.name = None

# Create "Type" label. Disease for tcga and long form label for GTEx
types = []
for sample in df.index:
    if sample.startswith('TCGA'):
        types.append('_'.join(
                tcga[tcga.submitter_id.str.contains(
                        sample)].project_name[0].split()))
    else:
        types.append('_'.join(
                list(gtex_sub[gtex_sub['id'] == sample].tissue)[0].replace(
                    '-', '').split()))
df['type'] = types
```

```
# Save
df.to_csv('tcga-gtex-metadata-intersect.tsv', sep='\t')
```

In [238]:
```
pd.set_option('max_columns', 4)
df.head()
```

Out[238]:
```
                                  id      reads  \
TCGA-CD-8534-01  TCGA-CD-8534-01  240016440
TCGA-ER-A19A-06  TCGA-ER-A19A-06  179705496
TCGA-C5-A1M8-01  TCGA-C5-A1M8-01  195313702
TCGA-D1-A0ZN-01  TCGA-D1-A0ZN-01   30911468
TCGA-EM-A4FF-01  TCGA-EM-A4FF-01  206307190

                                              ...                                \
TCGA-CD-8534-01                               ...
TCGA-ER-A19A-06                               ...
TCGA-C5-A1M8-01                               ...
TCGA-D1-A0ZN-01                               ...
TCGA-EM-A4FF-01                               ...

                tumor  \
TCGA-CD-8534-01   yes
TCGA-ER-A19A-06   yes
TCGA-C5-A1M8-01   yes
TCGA-D1-A0ZN-01   yes
TCGA-EM-A4FF-01   yes

                                                                 type
TCGA-CD-8534-01                                 Stomach_Adenocarcinoma
TCGA-ER-A19A-06                                Skin_Cutaneous_Melanoma
TCGA-C5-A1M8-01  Cervical_Squamous_Cell_Carcinoma_and_Endocervical_Adenoc...
TCGA-D1-A0ZN-01                     Uterine_Corpus_Endometrial_Carcinoma
TCGA-EM-A4FF-01                                      Thyroid_Carcinoma

[5 rows x 16 columns]
```

##### 11.2.5 Missing Data

In [239]:
```
df.isnull().sum()
```

Out[239]:
```
id                 0
reads              0
size_MB            0
platform           0
sex               39
tissue             0
seq_site           0
weight          8342
height          8428
mapped_reads    1048
race            1373
age              210
qc                 0
dataset            0
tumor              0
```

```
type                  0
dtype: int64
```

```
In [240]: print 'Number of samples with missing values: {} of {}'.format(
              sum(df.isnull().sum(axis=1) > 0), df.shape[0])
```

```
Number of samples with missing values: 9544 of 20852
```

```
In [241]: gr = df.columns.to_series().groupby(df.dtypes).groups
          gr = {k.name: v for k, v in gr.items()}
```

```
In [254]: f, ax = plt.subplots(4, 2, figsize=(10, 10))
          ax = ax.flatten()
          i = 0
          for s in gr['float64'] + gr['int64']:
              n = sum(df[s].value_counts())
              title = s + ' - NaNs: {}'.format(df.shape[0] - n)
              fg = sns.FacetGrid(df.dropna(), hue='dataset')
              fg = fg.map(sns.distplot, s, ax=ax[i])
              ax[i].set_title(title)
              i += 1
              plt.close(fg.fig)  # FacetGrid makes its own plot we don't want
          ax[-1].axis('off')
          f.subplots_adjust(hspace=.5)
          plt.tight_layout()
          plt.show()
```

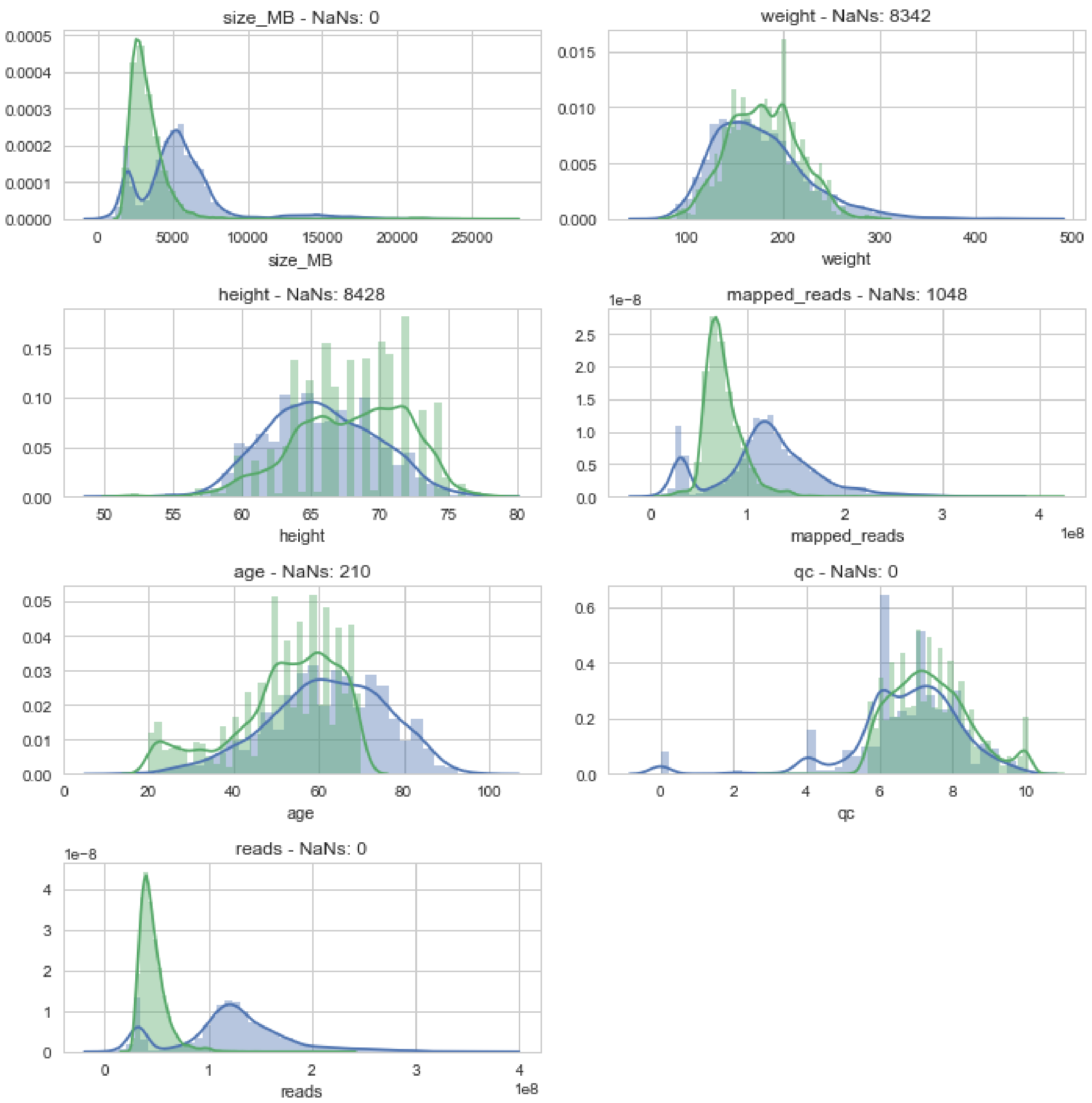

In [258]:
```python
f, ax = plt.subplots(3, 2, figsize=(10, 10))
ax = ax.flatten()
for i, s in enumerate(['platform', 'sex', 'seq_site', 'race', 'tumor']):
    n = sum(df[s].value_counts())
    title = s + ' - NaNs: {}'.format(df.shape[0] - n)
    sns.countplot(data=df, x=s, hue='dataset', ax=ax[i])
plt.setp(ax[3].get_xticklabels(), rotation=45)
plt.tight_layout()
f.subplots_adjust(hspace=.25)
ax[-1].axis('off')
plt.show()
```

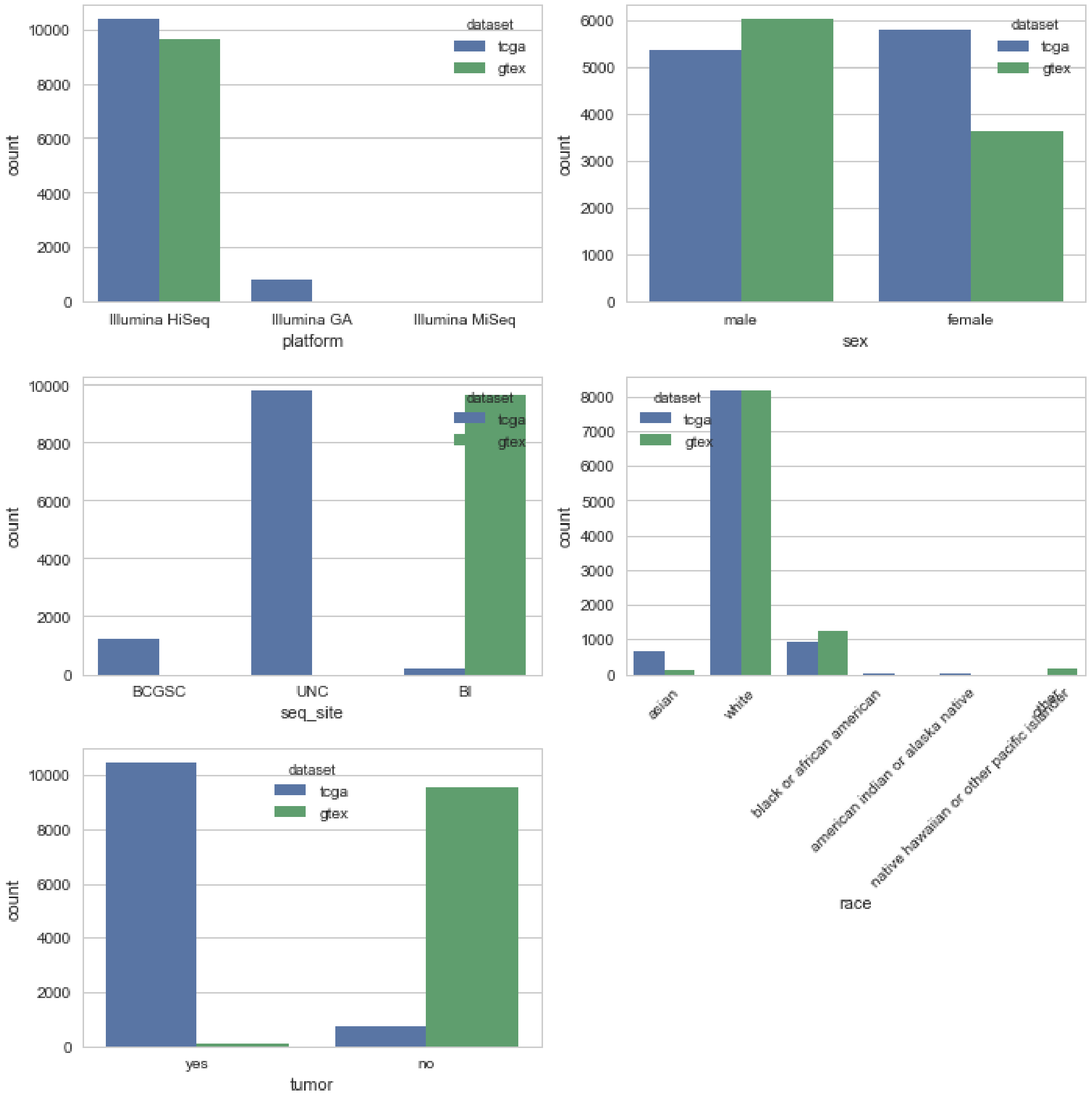

#### 11.3 Consolidated TCGA and GTEx HDF5 Object

contains:

- TCGA and GTEx Expression (Samples by Genes in TPM)
- TCGA Metadata
- GTEx Metadata
- Intersectional Metadata
- Drug Information

In [1]:
```
import pandas as pd

# Synapse ID: syn12117384
exp = pd.read_hdf('/mnt/data/Expression/tcga_gtex_tpm.hd5')
```

```
# Synapse ID: syn12117386
tcga_path = 'Metadata-collation/output/tcga-metadata-cleaned.tsv'
tcga_met = pd.read_csv(tcga_path, sep='\t', index_col=0)

# Synapse ID: syn12117387
gtex_path = 'Metadata-collation/input_data/gtex-sample-metadata.tsv'
gtex_met = pd.read_csv(gtex_path, sep='\t', index_col=0)

# Synapse ID: syn11363589
met_path = '/mnt/data/Metadata/tcga_gtex_metadata_intersect.tsv'
met = pd.read_csv(met_path, index_col=0, sep='\t')

# Synapse ID: syn12117759
drug_path = '/mnt/data/Metadata/drug_biomarkers.tsv'
drug = pd.read_csv(drug_path, index_col=0, sep='\t')

# Synapse ID: syn12246910
pairing = pd.read_csv('data/subtype-pairings.tsv', sep='\t', index_col=0)
```

##### 11.3.1 Gene Expression

```
In [2]: print exp.shape
        exp.head(2)
```

```
(18295, 58581)
```

```
Out[2]:                          5S_rRNA  5_8S_rRNA  7SK       A1BG  A1BG-AS1  A1CF  \
        GTEX-UTHO-1226-SM-3GAEE      0.0        0.0  0.0  11.079876  1.029968  0.00
        GTEX-146FH-1726-SM-5QGQ2     0.0        0.0  0.0   9.469890  2.459924  0.01

                                        A2M    A2M-AS1       A2ML1  A2ML1-AS1  \
        GTEX-UTHO-1226-SM-3GAEE  199.478958  0.609981    0.049999        0.0
        GTEX-146FH-1726-SM-5QGQ2  95.961578  0.869973  433.863281        0.0

                                     ...      snoU2-30  snoU2_19  snoU83B  snoZ196  \
        GTEX-UTHO-1226-SM-3GAEE      ...           0.0       0.0      0.0      0.0
        GTEX-146FH-1726-SM-5QGQ2     ...           0.0       0.0      0.0      0.0

                                 snoZ278  snoZ40  snoZ6  snosnR66    uc_338  \
        GTEX-UTHO-1226-SM-3GAEE      0.0     0.0    0.0       0.0  9.360242
        GTEX-146FH-1726-SM-5QGQ2     0.0     0.0    0.0       0.0  7.369859

                                 yR211F11.2
        GTEX-UTHO-1226-SM-3GAEE         0.0
        GTEX-146FH-1726-SM-5QGQ2        0.0

        [2 rows x 58581 columns]
```

##### 11.3.2 TCGA Metadata

```
In [3]: print tcga_met.shape
        tcga_met.head(2)
```

```
(11190, 31)
```

```
Out[3]:                        submitter_id  reads_downloaded  mapped_read_count  \
        TCGA-CD-8534-01A  TCGA-CD-8534-01A         240016440          207116695
        TCGA-ER-A19A-06A  TCGA-ER-A19A-06A         179705496          178311442

                                  auc        platform     file_size center_short  \
        TCGA-CD-8534-01A  15223006507  Illumina HiSeq  1.570993e+10        BCGSC
        TCGA-ER-A19A-06A   8517492780  Illumina HiSeq  7.741969e+09          UNC

                              tissue_source_site  aliquots_concentration  \
        TCGA-CD-8534-01A                  ILSBio                    0.15
        TCGA-ER-A19A-06A  University of Pittsburgh                  0.17

                          a260_a280_ratio       ...        years_smoked height  \
        TCGA-CD-8534-01A              NaN       ...                 NaN    NaN
        TCGA-ER-A19A-06A              1.7       ...                 NaN    NaN

                          year_of_diagnosis             project_name primary_site  \
        TCGA-CD-8534-01A             2011.0   Stomach Adenocarcinoma      Stomach
        TCGA-ER-A19A-06A             2006.0  Skin Cutaneous Melanoma         Skin

                          tumor_stage  vital_status    sample_type  \
        TCGA-CD-8534-01A     stage ii         alive  Primary Tumor
        TCGA-ER-A19A-06A     stage iv         alive     Metastatic

                          new_tumor_event_after_initial_treatment pathologic_stage
        TCGA-CD-8534-01A                                       NO         Stage II
        TCGA-ER-A19A-06A                                      YES         Stage IV

        [2 rows x 31 columns]
```

###### 11.3.3 GTEx Metadata

```
In [4]: print gtex_met.shape
        gtex_met.head(2)
```

```
(9662, 300)
```

```
Out[4]:                 Run  RailRnaBatchNumber  \
        rail_id
        3864      SRR660824                  19
        3865     SRR2166176                  19

                                                         BigWigPath  dbGaP_Subject_ID  \
        rail_id
        3864     /dcl01/leek/data/gtex/batch_19/coverage_bigwig...            678145
        3865     /dcl01/leek/data/gtex/batch_19/coverage_bigwig...            706551

                 dbGaP_Sample_ID     SUBJID                       SAMPID  \
        rail_id
        3864              914544  GTEX-QMR6       GTEX-QMR6-1926-SM-32PL9
        3865             1664018  GTEX-T5JC  GTEX-T5JC-0011-R11A-SM-5S2RX

                                      SAMPLE_USE   ReleaseDate      LoadDate  \
        rail_id
```

```
3864     Seq_RNA_WTSS; Seq_RNA_Expression  Feb 02, 2013  Oct 03, 2014
3865     Seq_RNA_WTSS; Seq_RNA_Expression  Oct 06, 2015  Sep 11, 2015

          ...     MHTTO012M  MHTTOONP  MHTXCEXP  MHUK8096  MHUREMIA MHWKNSSU  \
rail_id   ...
3864      ...           0.0       0.0       0.0       0.0       0.0      0.0
3865      ...           0.0       0.0       0.0       0.0       0.0      0.0

        MHWNVCT MHWNVHX MHWTLSUA MHWTLSUB
rail_id
3864        0.0     0.0      0.0      0.0
3865        0.0     0.0      0.0      0.0

[2 rows x 300 columns]
```

##### 11.3.4 TCGA/GTEx Intersectional Metadata

```
In [5]: print met.shape
        met.head(2)
```

```
(20852, 16)
```

```
Out[5]:                                 id      reads       size_MB        platform  \
        TCGA-CD-8534-01  TCGA-CD-8534-01  240016440  14982.158203  Illumina HiSeq
        TCGA-ER-A19A-06  TCGA-ER-A19A-06  179705496   7383.316781  Illumina HiSeq

                          sex   tissue seq_site  weight  height  mapped_reads   race  \
        TCGA-CD-8534-01  male  Stomach    BCGSC     NaN     NaN   207116695.0  asian
        TCGA-ER-A19A-06  male     Skin      UNC     NaN     NaN   178311442.0  white

                          age   qc dataset tumor                     type
        TCGA-CD-8534-01  40.0  0.0    tcga   yes   Stomach_Adenocarcinoma
        TCGA-ER-A19A-06  83.0  4.0    tcga   yes  Skin_Cutaneous_Melanoma
```

##### 11.3.5 Drug Information

```
In [6]: drug.head(2)
```

```
Out[6]:              Approved Biomarker(s) [RESPONSIVE]  \
        Drug
        ATEZOLIZUMAB                                NaN
        DURVALUMAB                                  NaN

                     Potential Biomarker(s) [RESPONSIVE]  \
        Drug
        ATEZOLIZUMAB                 CD274 overexpression
        DURVALUMAB                   CD274 overexpression

                     Approved Biomarker(s) [RESISTANT]  \
        Drug
        ATEZOLIZUMAB                               NaN
        DURVALUMAB                                 NaN

                     Potential Biomarker(s) [RESISTANT] Main Target(s)  \
```

```
Drug
ATEZOLIZUMAB                                   NaN   PD-L1, BCL-2
DURVALUMAB                                     NaN          PD-L1

             Other Target(s) Gene(s) Involved  \
Drug
ATEZOLIZUMAB             NaN      CD274, BCL2
DURVALUMAB              B7-1      CD274, CD80

                                         Primary Cancer Type(s)  \
Drug
ATEZOLIZUMAB  Locally advanced or metastatic (M1) urothelial...
DURVALUMAB    Locally advanced or metastatic urothelial carc...

             Secondary Cancer Type(s) Clinical Trial Cancer Type(s)  \
Drug
ATEZOLIZUMAB                      NaN                            NaN
DURVALUMAB                        NaN                            NaN

                         ...             Biomarker Paper  \
Drug                     ...
ATEZOLIZUMAB             ...                         NaN
DURVALUMAB               ...                         NaN

                                          Pharmocology Synopsis  \
Drug
ATEZOLIZUMAB  Atezolizumab is a humanized monoclonal immunog...
DURVALUMAB    Durvalumab is a programmed death-ligand 1 (PD-...

             Grouping of drugs →           Drug Type Subtype/Target Class  \
Drug
ATEZOLIZUMAB                 NaN  Monoclonal antibody                 CAM
DURVALUMAB                   NaN  Monoclonal antibody                 NaN

              Class/Family (if applicable) Mechanism of Action  \
Drug
ATEZOLIZUMAB                           NaN    CD274-inhibition
DURVALUMAB                             NaN    CD274-inhibition

             Pathway(s) Affected Notes Biomarker(s) for Sankey Graph
Drug
ATEZOLIZUMAB      PD-1 signaling   NaN         CD274 overexpression
DURVALUMAB        PD-1 signaling   NaN         CD274 overexpression

[2 rows x 23 columns]
```

##### 11.3.6 Subtype Pairings

In [7]:
```
pairing.head()
```

Out[7]:
```
                                Tumor                                Normal  \
0             Acute_Myeloid_Leukemia                                    NaN
1                                NaN                                    NaN
2                                NaN                                    NaN
3           Adrenocortical_Carcinoma                                    NaN
```

```
4  Pheochromocytoma_and_Paraganglioma  Pheochromocytoma_and_Paraganglioma

                               GTEx
0  Cells_Leukemia_cell_line_(CML)
1            Adipose_Subcutaneous
2      Adipose_Visceral_(Omentum)
3                   Adrenal_Gland
4                   Adrenal_Gland
```

##### 11.3.7 Write Consolidated HDF

**Key: Value** - exp: TCGA and GTEX gene expression (TPM) - tcga: TCGA Metadata - gtex: GTEx Metadata - met: TCGA/GTEx intersectional metadata - drug: Antineoplastic biomarkers and more

```
In [8]: output_hdf = '/mnt/data/Objects/tcga_gtex_data.hd5'
```

```
In [ ]: exp.to_hdf(output_hdf, key='exp')
        tcga_met.to_hdf(output_hdf, key='tcga')
        gtex_met.to_hdf(output_hdf, key='gtex')
        met.to_hdf(output_hdf, key='met')
        drug.to_hdf(output_hdf, key='drug')
        pairing.to_hdf(output_hdf, key='pairing')
```
